# A catalog of *cis*-regulatory mutations in 12 major cancer types

**DOI:** 10.1101/710103

**Authors:** Zhongshan Cheng, Michael Vermeulen, Micheal Rollins-Green, Brian DeVeale, Tomas Babak

**Author notes:** Authors contributed equally, correspondence.

## Abstract

Despite the recent availability of complete genome sequences of tumors from thousands of patients, isolating disease-causing (driver) non-coding mutations from the plethora of somatic variants is notoriously challenging, and only a handful of validated examples exist. By integrating whole-genome sequencing, gene expression, chromatin accessibility, and genetic data from TCGA, we identified 301 non-coding somatic mutations that affect gene expression in *cis*. These mutations cluster into 36 hotspot regions with diverse molecular mechanisms of gene expression regulation. We further show that these mutations have hallmark features of noncoding drivers; namely, that they confer a positive selection on growth, functionally disrupt transcription factor binding sites, and contribute to disease progression reflected in decreased overall patient survival.

## Introduction

Identification of somatic mutations that contribute to tumorigenesis is an essential step to understanding disease prognosis and developing therapies (Gerstung et al., 2015; Kulik, Pel’kis, & Korol, 1989; Verhaak et al., 2010). Despite extensive exome and genome sequencing efforts, a substantial proportion of causal or driver mutations (called *drivers* from here on) are thought to be unknown (Cancer Genome Atlas Research, 2014, 2015; Kandoth et al., 2013; Nik-Zainal et al., 2016). On average, 22.2% of tumor samples within each cancer type do not harbor coding mutations in any of 144 common driver genes (Schroeder, Rubio-Perez, Tamborero, Gonzalez-Perez, & Lopez-Bigas, 2014). Moreover, since multiple drivers are typically involved(Vogelstein et al., 2013), even tumors with well-characterized mutations likely harbor additional causal alterations (Beerenwinkel et al., 2007; Merid, Goranskaya, & Alexeyenko, 2014; Sjoblom et al., 2006; Vogelstein et al., 2013). Mutations in *cis-*regulatory elements (CREs) are postulated to comprise a large fraction of the undiscovered drivers (Sjoblom et al., 2006). However, despite the availability of hundreds of complete tumor genomes, only a few non-coding drivers have been experimentally validated (Table S2).

Distinguishing drivers from passengers outside coding regions requires overcoming several known challenges: the search space is orders of magnitude larger, functional impact cannot be predicted from amino acid changes (especially gain-of-function hotspots), mutation rates are higher (Poulos, Sloane, Hesson, & Wong, 2015), and positive selection pressure on relative growth is relaxed. These challenges have been partially overcome by associating mutations with alterations to transcription factor binding sites (Mathelier et al., 2015; Melton, Reuter, Spacek, & Snyder, 2015; Weinhold, Jacobsen, Schultz, Sander, & Lee, 2014), altered mRNA abundance (Fredriksson, Ny, Nilsson, & Larsson, 2014), clinical data (Smith et al., 2015; Weinhold et al., 2014), and evolutionary conservation (Carter et al., 2009; Foo et al., 2015; Fu et al., 2014; Piraino & Furney, 2017). Combinations of these features have also been weighed to prioritize putative drivers and determine significant mutational hotspots (Fu et al., 2014; Piraino & Furney, 2017; Puente et al., 2015; Weinhold et al., 2014).

Since the tumorigenic role of a noncoding driver is likely exerted through a *cis-* change in gene expression (Khurana et al., 2016), mapping genes whose expression is impacted by *cis-*acting regulatory effects has significant promise. Allele-specific expression (ASE), where one allele of a gene is more highly expressed than the other, is a powerful approach for detecting *cis-*regulatory effects, since *trans*-regulatory effects impact both alleles equally (Fraser, 2011). By comparing ASE in tumors to matched normal ASE (“diffASE”), it is further possible to distinguish somatic from germ-line effects. Ongen *et al*. applied this approach to identify 71 putative driver genes in colorectal cancer (Ongen et al., 2014). In practice, however, the sparse availability of matched tumor and normal gene expression and genetic data poses a significant limitation. Just 7.7% of TCGA tumor samples have matched normal RNA-Seq data (Fig. S1A).

Here we show that the vast majority of differential ASE is acquired in tumors, enabling us to dispense with the matched normal requirement and expand our survey 13-fold. We interrogated all whole-genome sequenced noncoding somatic mutations across 1,165 TCGA patients and identified 36 novel regulatory driver hotspots on the basis of robust association with ASE in tumors. The driver role of these mutations is further supported by elevated variant allele frequencies, functional disruption of transcription factor binding sites, and negative association with overall patient survival. This functional catalog of novel noncoding features significantly expands our knowledge of noncoding tumor driver biology.

### Survey of Breast Invasive Carcinoma (BRCA) reveals that >98% of differential ASE is due to ASE in tumor

We initially focused on BRCA since it is the cancer type with the largest set of matched tumor and normal data accompanied by whole genome sequence (WGS) in TCGA (Fig. S1A). Measuring ASE relies on counting RNA-Seq reads that map over heterozygous single nucleotide polymorphisms (SNPs) (Fig. 1A) detected by genotyping arrays. To maximize our sensitivity, we first imputed and phased SNPs using the 1000 Genome haplotypes (Howie, Donnelly, & Marchini, 2009) (Fig. 1A), which on average increases the number of informative SNPs 5-fold. This is more accurate than relying on WGS where sufficient coverage is not always available (Fig. S1B). Moreover, false-positive SNPs have a disproportionately high impact on estimates of ASE since all reads are assigned to one haplotype. Phasing also allowed us to combine allelic counts across SNPs within the same gene (Babak et al., 2015) (and see Methods for more details). We observed extensive diffASE in BRCA (Fig. 1B).

**Figure 1.**
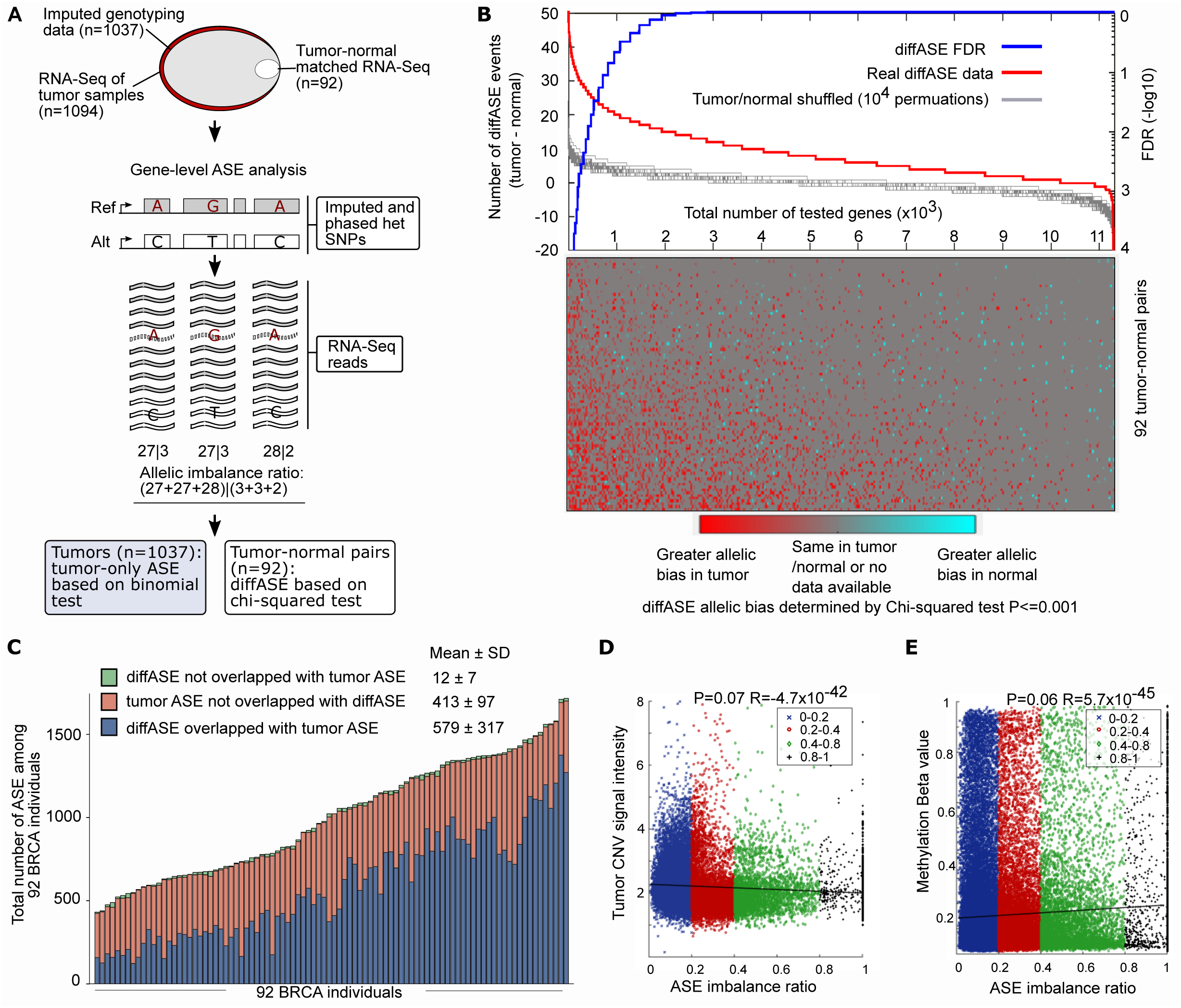
Allelic bias commonly arises in tumors independent of copy number variations (CNVs) and promoter methylation. (A) A schematic of the allele-specific expression (ASE) analysis strategy implemented on TCGA breast cancer samples. In brief, imputed genotyping data, tumor RNA-Seq and cases where tumor-normal matched RNA-seq are assessed for gene-level ASE by calculating the allelic imbalance ratios for imputed and phased heterozygous single-nucleotide polymorphisms (SNPs). We report differential ASE (diffASE) between tumors and matched normals and ASE in tumors (tumorASE) in cases when matched normals are unavailable. (B) diffASE events in breast cancer tumors exceed the background. A diffASE event is called between a tumor and its matched normal when the allelic ratio between them is P<0.001 using a chi-squared test and the skew is greater in the tumor. 632 diffASE events were obtained when the diffASE events calculated with the actual sample identities were compared to the background obtained with 10,000 permutations of randomized normal/tumor identities (FDR<0.05 and greater ASE in the tumor than the matched normal). The FDR reflects the proportion of permutations where the most significant diffASE event was obtained with the actual tumor/normal data. (C) >98% of diffASE events originate in breast cancer tumors. The breakdown of diffASE that overlaps tumorASE (P<0.001, binomial), does not overlap tumorASE, as well as the tumorASE that does not overlap diffASE by individual. (D) ASE does not correlate with CNV in BRCA tumors. The Pearson correlation between linear regression of gene-level ASE (ranging from 0 to 1) and tumor CNV signal intensity is weak and not significant (R=4.7×10^-42^, P=0.07, n=92). This analysis includes every gene exhibiting ASE (binomial P<0.001) in an individual tumor and excludes all others. The CNV signal intensity is obtained from CNV microarrays. (E) Gene-level diffASE is not correlated with the promoter (+/-2Kb from TSS) methylation (Pearson’s linear correlation R=5.7×10^-45^, P=0.06, n=92). As in Fig. 1D, all genes exhibiting ASE (binomial P<0.001) in an individual tumor were included. The beta-value is the ratio of methylated to total probe intensity.

Strikingly, nearly all of the diffASE can be attributed to an increase of ASE in tumors relative to matched controls (Fig. 1B). We reasoned that this trend may be due to higher clonality of tumors relative to matched normal tissue which would be expected to be more complex. We first considered whether loss of heterozygosity (LOH) may be a confounding factor. We reasoned that since all BRCA tumors are female, a comparison of allelic expression between autosomes to the X-chromosome could illuminate the contribution of clonality. X-chromosomes are randomly inactivated across cells comprising normal tissue. Comparison with a clone derived from this tissue (where all cells retain monoallelic expression from the same allele) would yield strong diffASE for any expressed gene on chromosome X. If clonality was the dominant source of greater ASE in tumors, we would expect enrichment of highly ranked X-linked genes when evaluated for diffASE. This enrichment would not be expected if LOH was the dominant source. We indeed observed a high enrichment of X-linked genes (66/100) among the top diffASE genes, suggesting that these tumors are highly clonal (Fig. S1D). When we performed the analysis of ASE using tumor expression data alone (tumorASE), we observed a recapitulation of >98% of the diffASE events (Fig. 1C). Finally, ASE events originating in tumors were not strongly correlated with CNV or methylation (Fig. 1D, E), which we estimate collectively explain less than 1% of ASE in tumors. In summary, we believe that altered *cis-*regulatory mechanisms of gene expression explain the majority of observed ASE in tumors, and that this signal is a valuable starting point for identifying noncoding drivers.

### Identification of mutations that explain ASE in tumors

The availability of WGS data for 113 BRCA RNA-seq samples (Fig. S1A) allowed us to find specific mutations that are associated, and which may explain, the observed ASE in tumors. We evaluated common mutation callers and implemented a robust filtering scheme to yield high confidence somatic variants (see Methods for details). We then asked whether the presence or absence of these variants near a gene is associated with ASE of that gene across BRCA tumor samples. Unfortunately, using the entire region surrounding a gene did not yield associations that survived multiple test correction, even in this heavily surveyed cancer type. The high proportion of neutral mutations relative to genuine noncoding drivers likely explains this result, and necessitates an enrichment strategy for variants that are likely to have a functional impact.

The vast majority of previously validated noncoding driver mutations occur in promoters, enhancers, and CTCF binding sites (Table S1). As these collectively encompass major sites of transcriptional regulation, we focused on somatic variants within these features and refined them using several publically available annotation resources. To comprehensively map genomic regions where transcription is regulated, we also included an aggregate map of TF binding sites (‘ChIP-seq’). For the enrichment analysis, we grouped the somatic mutations in promoters, CTCF and TF binding sites by regulatory feature and asked if they were 10kb upstream of a TSS or gene body of a gene exhibiting ASE. Conversely, since enhancers vary by cell type and frequently regulate non-adjacent genes, we used cancer-specific enhancers and regulatory relationships defined by associations between accessible chromatin and changes in gene-expression (Corces et al., 2018). Putative drivers were identified by positive correlations between gene-level ASE and somatic regulatory mutations (see Methods). This approach revealed novel non-coding driver mutations regulating genes previously implicated in breast cancer by coding variants as well as altered regulation of novel genes.

Using these features, we found somatic mutations in the regulatory elements of seven genes that are enriched for altered *cis-*regulation in breast cancer (FDR<=0.9, n=113). These include mutations in the enhancers of *EGFR, CDC42EP3* and *TIMP3*, variants in the promoters of *UNC5B*, CTCF binding sites of *DAAM1* and *NOTCH1* as well as TF binding sites near *ITPR3* (Fig. 2A). Altered *cis-*regulation of *EGFR* is particularly notable as coding mutations in it are among the most common cancer drivers (Fig. 2B, C) (Bailey et al., 2018; Wee & Wang, 2017). In the 4 tumors harboring enhancer mutations, dysregulation is evident from the ASE ratio of 3.81 compared to 1.68 in tumors where they are not mutated (Fig. 2B, P=3.64×10^-3^, Pearson’s linear correlation).

**Figure 2.**
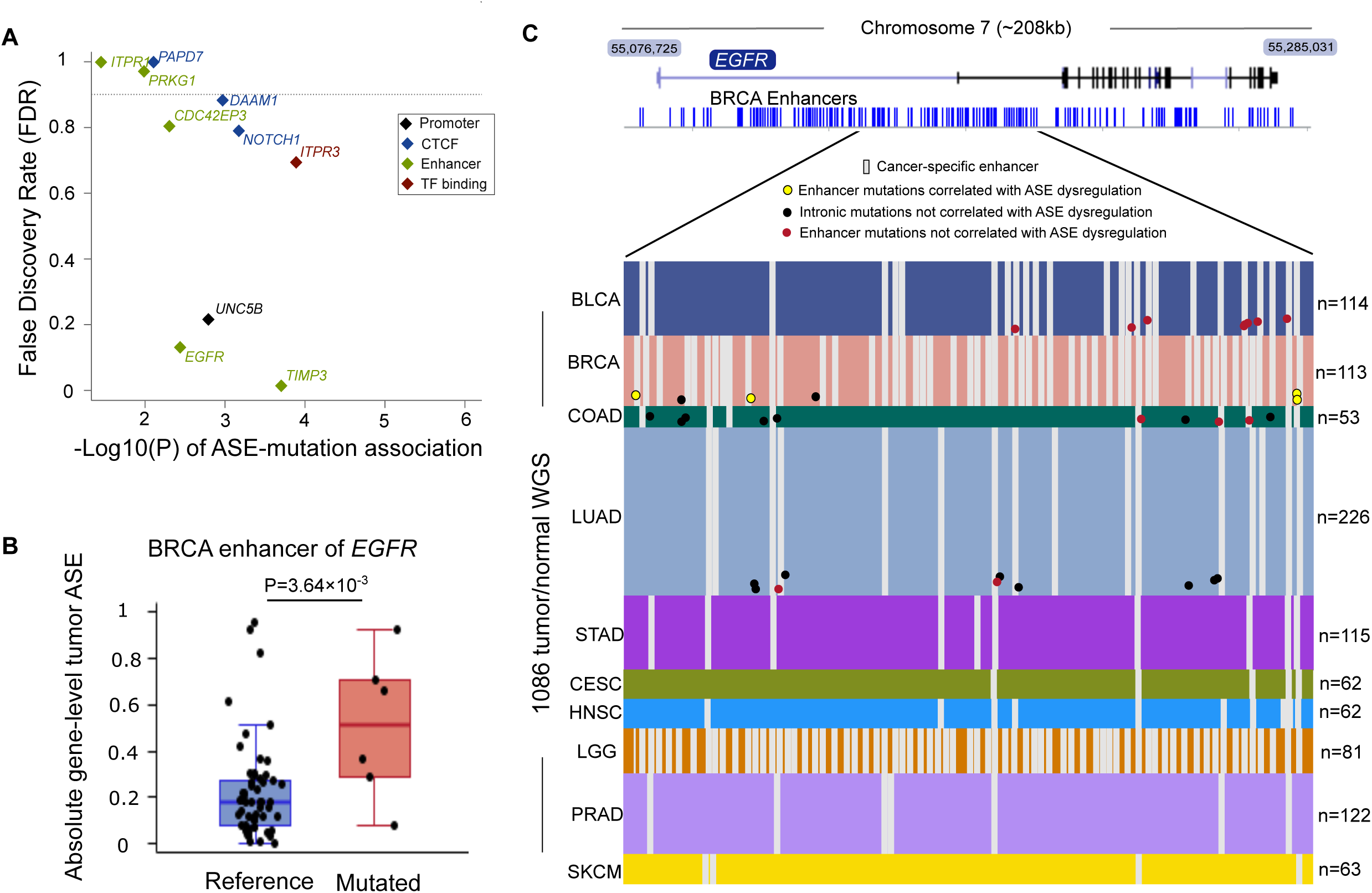
Seven somatically mutated regulatory features are enriched for ASE in breast cancer. (A) The significance of gene-level associations between mutated regulatory features and ASE in relation to FDR in breast cancer. The association of gene-level ASE was evaluated with a Pearson co-efficient of determination. The FDR calculated by permuting the identity of tumor samples for both ASE and mutation values (see methods for detail). Each association was performed genome-wide. (B) The ASE ratio of putative BRCA driver *EGFR* (P=3.64×10 ^-3^, FDR=0.014, n=6). The boxplot is delimited by the first and the third quartile, while the diamonds and dots represent the medians and raw absolute gene-level ASE, respectively. (C) The distribution of somatic mutations in the enhancer of *EGFR* among the 10 cancers analyzed (n=1,086 tumors). Each row represents a sample and each column represents 1 bp. Cancer-specific enhancers were not available for OV and LAML.

### Somatic mutations in regulatory features are enriched for gene-level ASE in diverse tumors

We applied our pipeline to 12 cancer types that had a sufficient number of matched WGS, RNA-Seq, genotyping, and chromatin accessibility data (derived by ATAC-seq (Corces et al., 2018)) (Fig. S1A). We identified 36 putative driver hotspots consisting of 301 mutations (Fig. 3, Table S1), which we will collectively refer to as the “novel driver candidates”. Notable examples of novel drivers based on prevalence include the enhancers of *ME3* in colon adenocarcinoma (COAD; 13.2%; n=7/53), the CTCF bound regions of *CBLB* in acute myeloid leukemia (LAML; 12.2%; n=5/41) and the CTCF bound region of *SEMA4D* in lung squamous cell carcinoma (LUAD; 11.5%; n=26/226). Interestingly dysregulation of *COL4A1* was found in two cancer types, including LUAD and COAD (Table 1 and S1). The majority of genes impacted by our novel drivers have been implicated previously in cancer and compelling cases for their driver mechanistic roles are explored further below (see Discussion).

**Table 1.** Annotated catalog of the 36 putative driver hotspots.

| Cancer | Regulatory feature | Gene symbol | P | FDR | Carriers | ASE-CNV Assoc P |
| --- | --- | --- | --- | --- | --- | --- |
| BLCA | TF binding | *SFMBT2* | $8.8 \times 10^{-4}$ | 0.55 | 4 | 0.78 |
| BLCA | TF binding | *SLC39A11* | $5.1 \times 10^{-5}$ | 0.14 | 7 | 0.61 |
| BLCA | TF binding | *TPCN1* | $1.1 \times 10^{-3}$ | 0.60 | 3 | 0.18 |
| BRCA | Cancer-specific enhancer | *CDC42EP3* | $5.0 \times 10^{-3}$ | 0.80 | 3 | 1.00<sup>#</sup> |
| BRCA | CTCF | *DAAM1* | $1.1 \times 10^{-3}$ | 0.88 | 5 | 0.83 |
| BRCA | Cancer-specific enhancer | *EGFR* | $3.6 \times 10^{-3}$ | 0.13 | 10 | 0.88 |
| BRCA | TF binding | *ITPR3* | $1.3 \times 10^{-4}$ | 0.69 | 5 | 0.72 |
| BRCA | CTCF | *NOTCH1* | $6.7 \times 10^{-4}$ | 0.79 | 4 | 0.21 |
| BRCA | Cancer-specific enhancer | *TIMP3* | $2.0 \times 10^{-4}$ | 0.01 | 8 | 0.47 |
| BRCA | Promoter | *UNC5B* | $1.9 \times 10^{-3}$ | 0.22 | 4 | 1.00<sup>#</sup> |
| CESC | TF binding | *SULF1* | $4.0 \times 10^{-3}$ | 0.22 | 3 | 1.00<sup>#</sup> |
| COAD | Promoter | *COL4A1* | $3.4 \times 10^{-3}$ | 0.50 | 3 | 0.68 |
| COAD | Promoter | *GSTA4* | $2.9 \times 10^{-3}$ | 0.47 | 3 | 1.00<sup>#</sup> |
| COAD | Cancer-specific enhancer | *ME3* | $3.7 \times 10^{-3}$ | 0.81 | 7 | 1.00<sup>#</sup> |
| HNSC | Promoter | *WLS* | $6.2 \times 10^{-3}$ | 0.08 | 3 | $*6.7 \times 10^{-3}$ |
| LAML | CTCF | *CBLB* | $3.7 \times 10^{-4}$ | 0.05 | 4 | 1.00 |
| LUAD | Cancer-specific enhancer | *ATXN1* | $2.6 \times 10^{-4}$ | 0.38 | 9 | $*2.2 \times 10^{-2}$ |
| LUAD | Cancer-specific enhancer | *CACNA1H* | $1.1 \times 10^{-3}$ | 0.47 | 7 | 0.68 |
| LUAD | Cancer-specific enhancer | *COL4A1* | $2.2 \times 10^{-4}$ | 0.36 | 3 | $*4.5 \times 10^{-4}$ |
| LUAD | TF binding | *ERI2* | $1.8 \times 10^{-6}$ | 0.67 | 3 | 0.29 |
| LUAD | Promoter | *GATA3* | $3.4 \times 10^{-6}$ | 0.17 | 4 | 0.95 |
| LUAD | TF binding | *ITGAE* | $8.1 \times 10^{-7}$ | 0.54 | 11 | 0.20 |
| LUAD | Cancer-specific enhancer | *JCAD* | $4.0 \times 10^{-3}$ | 0.89 | 8 | 0.58 |
| LUAD | CTCF | *LPAR1* | $2.5 \times 10^{-5}$ | 0.83 | 24 | 0.60 |
| LUAD | Cancer-specific enhancer | *PIP5K1B* | $1.1 \times 10^{-5}$ | 0.14 | 5 | 0.21 |
| LUAD | CTCF | *SEMA4D* | $1.2 \times 10^{-6}$ | 0.31 | 26 | $*1.4 \times 10^{-2}$ |
| LUAD | TF binding | *SND1* | $7.4 \times 10^{-8}$ | 0.22 | 24 | 0.88 |
| LUAD | Cancer-specific enhancer | *TBCD* | $1.3 \times 10^{-4}$ | 0.32 | 4 | 0.63 |
| LUAD | Cancer-specific enhancer | *TRIM2* | $6.7 \times 10^{-4}$ | 0.45 | 8 | 0.95 |
| OV | TF binding | *PKNOX2* | $2.4 \times 10^{-3}$ | 0.47 | 4 | 0.49 |
| OV | CTCF | *SLC27A1* | $7.7 \times 10^{-3}$ | 0.56 | 4 | $*3.2 \times 10^{-2}$ |
| SKCM | CTCF | *KANSL1* | $8.5 \times 10^{-3}$ | 0.10 | 3 | 1.00<sup>#</sup> |
| STAD | CTCF | *ARSB* | $2.3 \times 10^{-4}$ | 0.27 | 3 | $*2.5 \times 10^{-3}$ |
| STAD | Cancer-specific enhancer | *CCND2* | $5.4 \times 10^{-2}$ | 0.87 | 3 | 0.40 |
| STAD | CTCF | *PAG1* | $5.5 \times 10^{-3}$ | 0.87 | 4 | 0.59 |
| STAD | TF binding | *PBX1* | $2.1 \times 10^{-4}$ | 0.56 | 4 | 0.56 |
| STAD | TF binding | *TSHZ2* | $2.7 \times 10^{-6}$ | 0.10 | 4 | $*1.5 \times 10^{-2}$ |

Note: \*Among all 7 genes demonstrating association between ASE and CNV, no genes have tumors carrying driver mutations and CNV coincidentally, except for *TSHZ2* (STAD) where only one tumor carrying both driver mutation and CNV; however, the association between mutations and ASE remained significant after excluding the tumor ( $P=2.2 \times 10^{-5}$ ). #When a driver gene doesn't have more than 1 sample harboring CNV for the association test between ASE and CNV, the association is assigned as $P = 1$ . CNV: copy number variation; P: ASE-mutation association P-value; ASE-CNV Assoc P: ASE-CNV association P-value; FDR: false discover rate; Carriers: driver mutations carriers.

**Figure 3.**
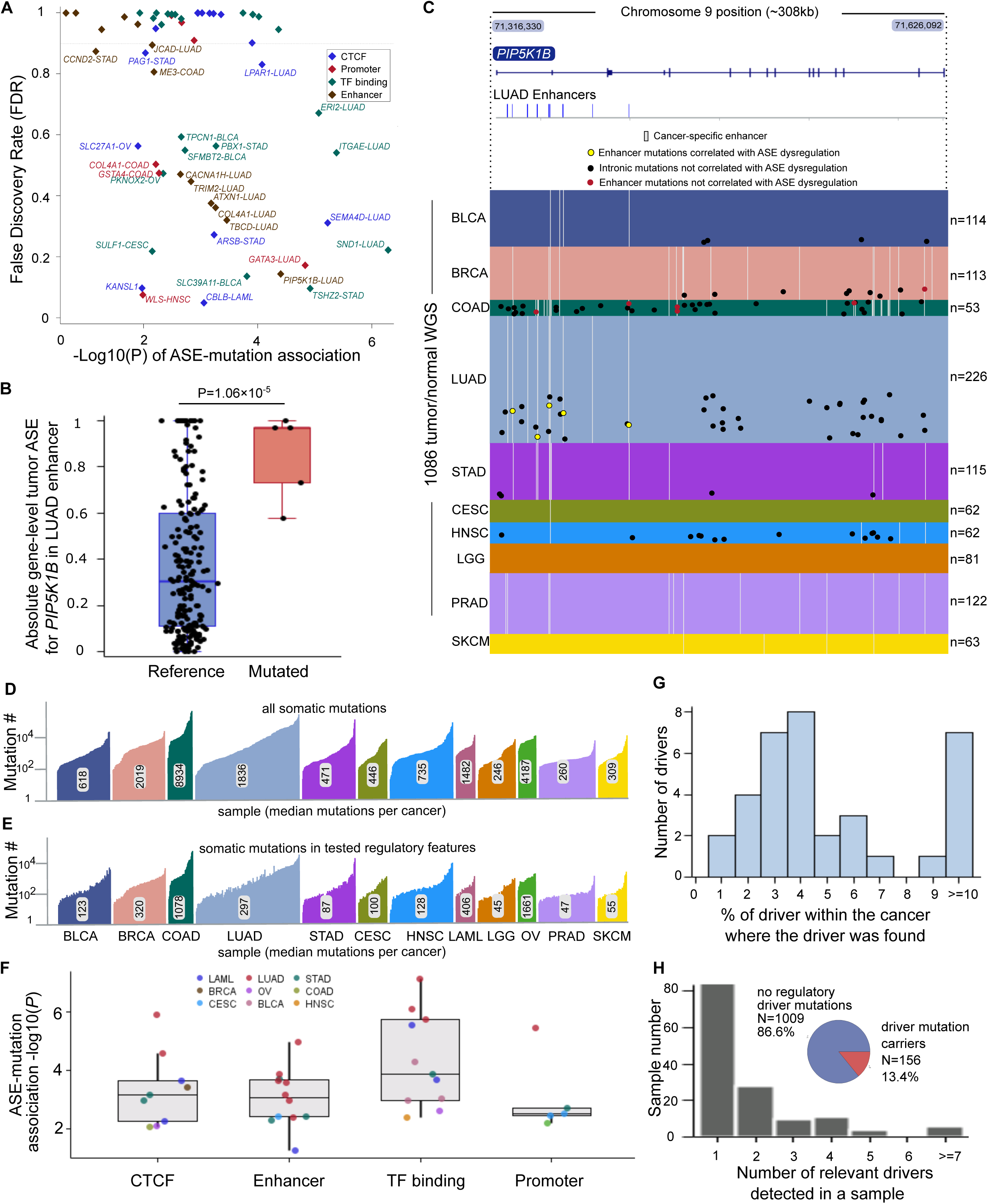
Thirty somatically mutated regulatory features are enriched for ASE across 11 additional cancer types. (A) The significance of gene-level associations between mutated regulatory features and ASE in relation to FDR in 11 additional cancers. The association of gene-level ASE was evaluated with a Pearson co-efficient of determination. The FDR calculated by permuting the identity of tumor samples for both ASE and mutation values (see methods for detail). Each association was performed genome-wide. (B) The ASE ratio of putative LUAD driver *PIP5K1B* (P=1.06×10 ^-5^, FDR=0.14, n=5). The boxplot is delimited by the first and the third quartile, while the diamonds and dots represent the medians and raw absolute gene-level ASE, respectively. (C) The distribution of somatic mutations in the enhancer of *PIP5K1B* among the 10 cancers analyzed (n=1,086 tumors). Each row represents a sample and each column represents 1 bp. Cancer-specific enhancers were not available for OV and LAML. Mutations of *PIP5K1B* enhancers are putative drivers of LUAD. (D) The number of somatic mutations in each tumor as well as the median number for each type of cancer. (E) The number of somatic mutations in regulatory features that were tested in the ASE-Mut association in each tumor as well as the median number for each type of cancer. (F) The distribution of driver significance plotted by the type of regulatory feature that is mutated. (G) The penetrance (% of all tumors) of mutated regulatory features in the cancer type where they are enriched for ASE. (H) The distribution of mutated regulatory features between samples. The inset illustrates that the majority of samples do not harbor driver mutations.

### Novel driver candidates have elevated variant allele frequencies

By definition, driver mutations confer a selective advantage to the cells in which they occur. Variant allele frequency (VAF) measures the fraction of alleles in a sample in which the variant is present. Hence, if a mutation confers a selective advantage to the cell in which it occurs, its VAF would be higher, on average, than passenger mutations that arose coincidentally. A corollary being that mutations with increased VAF occurred early enough during tumor evolution for this selective advantage to manifest as increased VAF. To ask whether driver mutations conferred a selective advantage, we compared the normalized VAF of all putative drivers to all non-coding mutations that were not enriched for ASE (P>0.5). As a positive control we used known coding driver mutations(Schroeder et al., 2014). As expected, we found that the VAF of known coding drivers (n=116) was, on average, higher than background mutations in coding regions (P-value=7.5×10 ^-6^, n=2,971). Importantly we found that the VAF of our novel driver candidates was also higher (Fig. 4A), an effect that is independent of CNV based on the stable ratio of adjacent heterozygous SNPs (Table 1 and S1).

**Figure 4.**
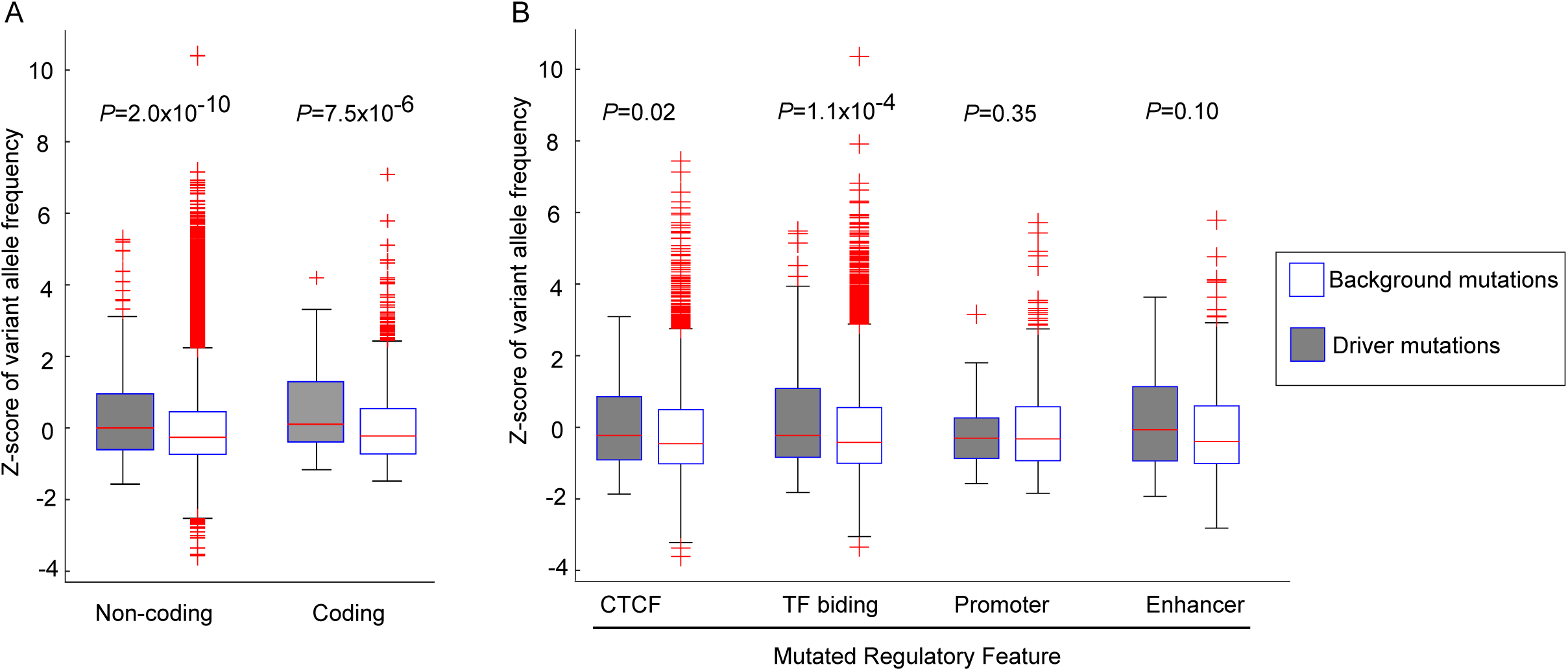
Variant allele frequency (VAF) of noncoding drivers suggests positive selection. VAF was calculated as the fraction of all sequencing reads covering variant with mutation and was normalized against all mutations within each patient to account for differences in tumor heterogeneity. (A) Coding Drivers (n=116) represent all mutations within known driver genes that yield a functional amino acid change and Coding Bcg (n=2,971) represent identically selected mutations in all other coding genes(Schroeder et al., 2014). Noncoding Drivers represent all mutations from Table S1 (n=301) and Noncoding Bcg represent all noncoding mutations not enriched for ASE (n=122,603). Both Coding and Noncoding VAFs are positively shifted relative to background (P=7.5×10 ^-6^, P=2.0×10 ^-10^). Please see Methods for more details. (B) Same as (A) with noncoding drivers divided by feature.

### Novel driver candidates disrupt transcription factor binding motifs

To begin to explore the mechanisms through which our driver mutations may be acting, we asked whether they may impact TF binding. TF binding affinities are typically represented by a generalized position-weight matrix (PWM) that represents a motif and a probability of observing any of the four bases at each position in that motif. These probabilities are typically constructed from observed frequencies of genuine binding events and can be represented as bit-scores. A bit-score of 2 implies that a particular base is always found at that position. The challenge with relying on PWMs exclusively to identify transcription factor binding is that there is typically insufficient information to distinguish genuine binding sites from the many possible motif sequence matches in the genome. To enrich for genuine binding sites we only considered mutations in our functionally annotated regions and required that mutations exceed a minimal degree of evolutionary conservation (Phastcons>0.05;(Siepel et al., 2005)).

If our driver mutations were causing ASE by disrupting binding at one of the alleles, we would expect to see a greater impact on the difference in bit scores (i.e. delta-bit) in ASE (driver) vs. non-ASE (background) mutations. We observed this effect in our promoter and CTCF features (Fig. 5). It is also possible that improving binding (i.e. a positive delta-bit) could cause ASE, although we did not see evidence for this which may be due to power constraints stemming from “functional gain” in binding being less frequent than “functional loss”.

**Figure 5.**
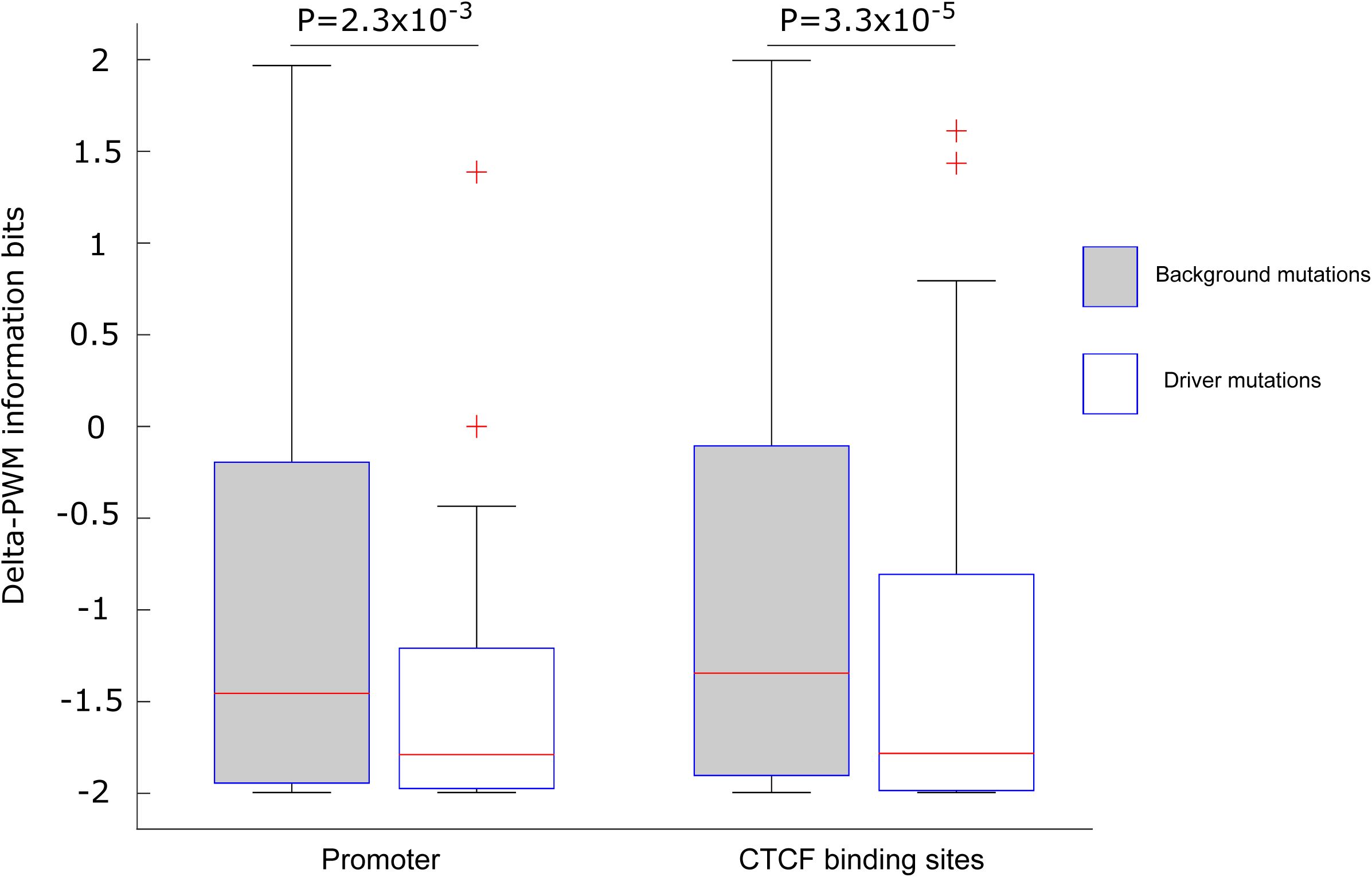
Driver mutations preferentially disrupt transcription factor binding motifs. The damage driver and passenger mutations caused to transcription factor binding sites was compared using delta-bite scores (delta-PWM information bits). Driver mutations were more disruptive to transcription factor binding sites in both promoters (P-value = 2.3×10 ^-3^) and CTCF binding sites (P-value = 3.3×10 ^-5^); Mann-Whitney U tests were performed to determine the difference of delta bit-scores between driver mutations and background. All driver mutations are with FDR<1, and the background mutations’ FDRs are equal to 1. There are 53 and 121 driver mutations and 2,292 and 14,514 background mutations found promoters and CTCF binding sites, respectively.

### Novel driver candidates correlate with reduced survival

To assess the impact of the putative drivers on patients, we asked how they effected survival. For this analysis, we grouped patients across all cancer types and features to enhance statistical power. Since each cancer type alone yields vastly different survival outcomes (J. Liu et al., 2018), we constructed carefully matched background sets with cancer type proportions constrained (see Methods). The overall survival (OS) of patients whose tumors were driven by any of the 36 non-coding drivers (Table 1 and S1) was worse than patients with a matched mutation burden not associated with ASE. The lower OS is also evident when comparing non-coding driver carriers to background versus 5,000 random subsets of the entire cohort (Fig. 6A). The negative impact of these drivers is clear when visualizing survival of patients carrying the non-coding drivers to one random subset of the whole dataset (Fig. 6B). Next we compared the impact of mutations in different classes of regulatory elements in different cancers on survival. The impact of driver mutations in promoters as well as CTCF and TF binding sites was predominantly negative (Fig. 6C). These analyses indicate that mutation of distinct classes of regulatory features may contribute to disease progression.

**Figure 6.**
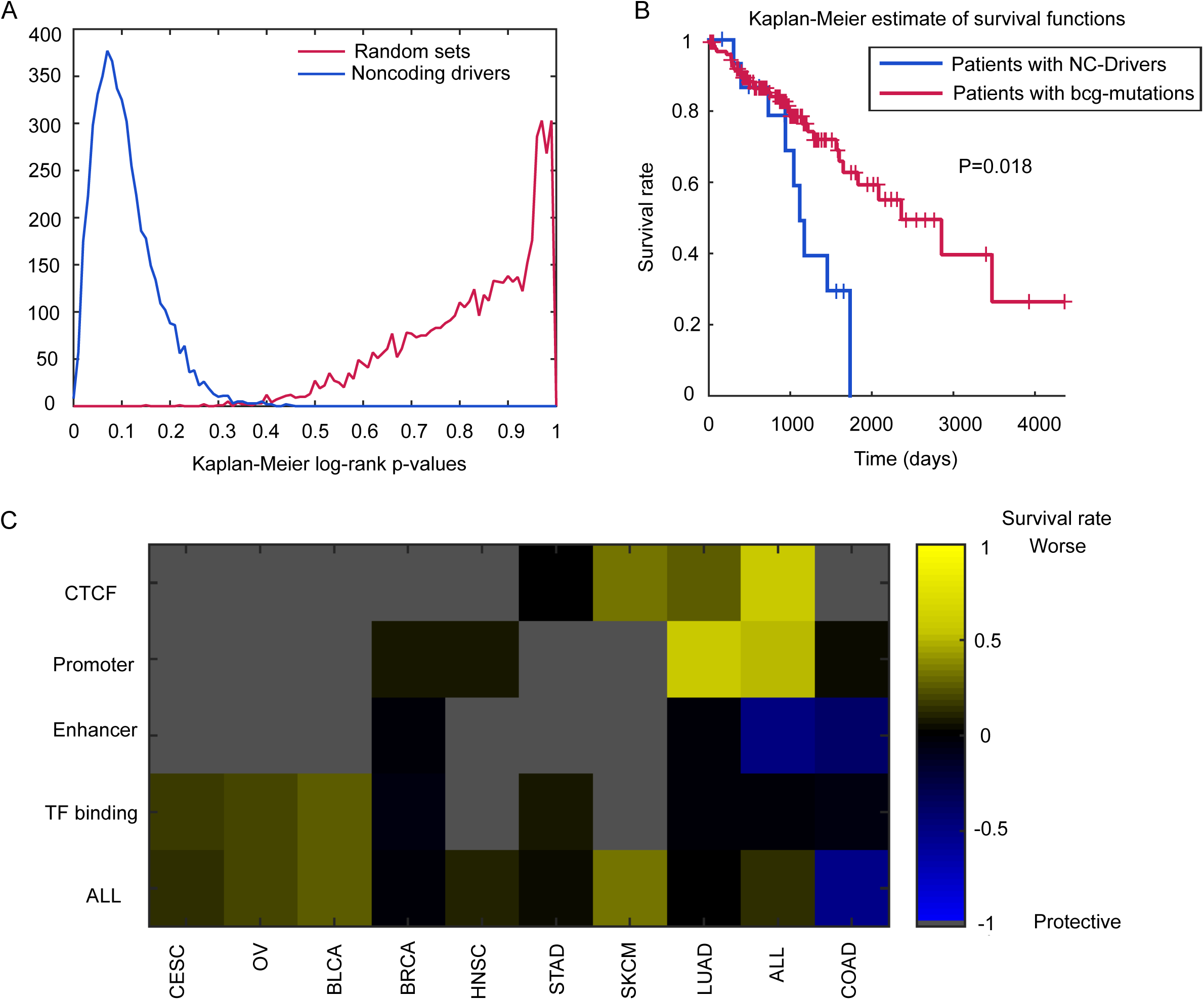
Non-coding drivers correlate with a worse overall survival outcome. (A) Sample Kaplan-Meier survival outcome analysis of patients harboring high-confidence non-coding drivers (Table S1; FDR<0.5) versus patients with mutations not enriched for ASE, matched for cancer and feature type (see Methods). Distribution of log-rank P-values (2-tailed) for 5,000 iterations shown in inset (blue) as well as random sets versus background (magenta). ‘+’ data points indicate censored data. (B) Survival analyses for mutations considered in (A) broken down by cancer and feature type.

## Discussion

Our 36 mutation hotspots associated with ASE significantly expand the landscape of noncoding cancer drivers. The majority of our findings are novel, although there is parital overlap with previous noncoding driver discoveries. For example, we found enriched *cis-*regulatory mutations in the CTCF binding sites of *DAAM1* and *NOTCH1* in BRCA. DAAM1 is a member of the formin protein family activated by Dishevelled binding (W. Liu et al., 2008). It regulates cytoskeletal dynamics through its control of linear actin assembly (D. Li et al., 2011). Regulatory mutations in *DAAM1* were recently implicated in invasiveness of melanoma (Zhang et al., 2018). Similarly, mutations in the 3’ UTR of *NOTCH1* were previously implicated in chronic lymphocytic leukaemia (Puente et al., 2015). Our findings also overlap previous reports in that somatic mutations in other regulatory regions of the same genes in the same type of cancer have been implicated as drivers. For example, mutations in the splice-acceptor site of *GATA3* were previously implicated in LUAD (Hornshoj et al., 2018). Here we implicated promoter mutations in *GATA3* in LUAD. This overlap suggests that the consequences of mutated regulatory features may overlap in these cases, and that combining the association of distinct features that regulate the same gene may increase sensitivity.

Many of the genes impacted by noncoding drivers discovered here (Table 1 and S1) have been previously implicated in cancer biology. The BRCA hotspots illustrate how driver roles clearly tie into the established functions of the dysregulated genes. Binding of a variety of ligands to EGFR promotes cell survival and proliferation via ERK signaling, and its mutation is one of the most common cancer drivers (Bailey et al., 2018; Wee & Wang, 2017). NOTCH1 is a key mediator of signaling between adjacent cells. It is essential in specification of a variety of cell types, maintaining tissue homeostasis and its mutation is a highly prevalent driver in certain cancers, such as T-cell Acute Lymphoblastic Leukemia (Bigas & Espinosa, 2018). ITPR3 mediates the release of intracellular calcium in response to IP3 (Yamamoto-Hino et al., 1994). It was recently implicated as the target of the tumor suppressor BAP1 that triggers apoptosis following exposure to genotoxic stress (Bononi et al., 2017). UNC5B is a netrin family receptor that mediates guidance of the vascular system as well as other cell types (Lu et al., 2004; Tai-Nagara et al., 2017). It encodes a repulsive netrin whose disruption results in aberrant extension, branching and navigation of affected cells (Lu et al., 2004). CDC42EP3 is a Cdc42 effector that regulates septin organization through binding to septin GTPases (Joberty, Perlungher, & Macara, 1999; Joberty et al., 2001). The tumor growth promoting activity of cancer-associated fibroblasts requires *CDC42EP3* (Calvo et al., 2015). Finally, TIMP3 is an inhibitor of matrix metalloproteinases whose upregulation suppresses tumor growth (Anand-Apte et al., 1996).

In contrast with previous reports, somatic mutations in regulatory features, not CNVs or differential methylation, underlie the majority of altered *cis-*regulation in tumors (Mayba et al., 2014). Somatic variants in non-coding regions that are enriched for altered *cis-*regulation were found in 13.4% of the tumors analyzed. This high-prevalence is predicted by multi-hit models as well as divergent phenotypes between tumors with common known drivers. While many of the associations involve genes thought to be involved in tumorigenesis, the implication of specific mutations and regulatory features is novel. Indeed, we are not aware of any of the specific mutated regulatory features reported here previously being implicated as drivers of tumorigenesis.

Although TCGA and other emerging cancer data now include >1000 available genomes, illuminating the complete set of noncoding drivers will require a substantially broader collection. Even with the approach employed here of focusing on functional somatic variants with underlying evidence of gene expression regulation, we found ourselves limited by statistical power, especially in cancer types with fewer than 100 genomes. Deeper genome sequencing with longer reads will also improve driver detection sensitivity by enabling phasing of mutations with the direction of ASE. This would allow more evidence to be used to prioritize genuine drivers (e.g. disruption of an activating transcription factor binding site should reduce expression of that allele). This was generally not possible with the current available data since accurate phasing of somatic variants more than a few hundred base pairs away from the gene would require long-read technology or much deeper coverage. Improved matching of the regulatory features to each cell type will also improve sensitivity. When possible, cellular context was prioritized throughout these analyses to account for context-specific aspects of gene-regulation. For example, enhancers were matched to the cancer type being analyzed (Corces et al., 2018), and each cancer was separately analyzed in parallel, however, enhancer to gene maps are still incomplete and will no doubt improve with more chromatin accessibility readouts expand. In any case, we believe our approach here, made freely available as a dockerized pipeline (see Methods) will be a powerful tool for taking advantage of these emerging resources and building on our discoveries.

## Methods

### Genotyping and imputation

Genome-wide Affymetrix 6.0 genotype array datasets from normal blood samples were downloaded as Birdseed files from GDC Legacy Archive (https://portal.gdc.cancer.gov/legacy-archive/search/f) for all 5,875 patients from 12 cancer types. Among these patients, only the 1,165 where tumor RNA-seq and matched tumor/normal WGS data were available were included in the downstream association between gene-level ASE and mutation occurrence (see Fig. S1A). These datasets were annotated with Affymetrix annotation files and converted into base-level genotypes. To minimize allelic mapping bias we excluded SNPs with more than 2 polymorphisms or those where 2 SNPs conflicted at the same site on the same strand in phased 1000 Genomes Project Phase1 v3 data. Affymetrix 6.0 arrays genotype nearly 1 million SNPs. Typically ∼25% of these sites are heterozygous and only a small fraction fall within expressed regions (mean=12,468). To increase the number of SNPs available to resolve ASE, we imputed and phased genotypes as previously described in Babak and DeVeale *et al*. (Babak et al., 2015). In brief, genotyping data were transformed into PLINK binary format and subjected to pre-phasing with Shape-IT software (v2.r790)(Delaneau, Coulonges, & Zagury, 2008) using the 1000 Genomes Project Phase1 v3 data as the reference, then imputed and phased using Impute2 software (v2.3.2) (Howie et al., 2009). We imputed with default parameters and used phased 1000 Genomes Project Phase1 v3 data as the reference panel. For each individual, heterozygous SNPs with genotype probability ≥ 0.95 were retained, as well as the allelic status within phased haplotypes. This provided an average of 4-fold additional SNPs per individual.

### RNA-seq

Matched tumor/normal RNA-Seq BAM files were downloaded from GDC Legacy Archives for all 1,165 patients across these 12 cancer types. SAMtools (H. Li et al., 2009) mpileup was used to calculate the reads at each heterozygous SNP site in all the RNA-seq BAM files aligned to the GRCh37-lite human reference genome with BWA. Default SAMtools mpileup settings were used except for a maximum read depth of 8,000 in order to reduce the bias of bases showing excessive depth and conserve computational resources.

### Gene-level ASE

To generate gene-level ASE ratios, heterozygous SNPs in each individual were mapped to a custom human transcript track generated by aggregating Ensembl (v80), UCSC and NCBI transcripts. Gene-level ASE was calculated by summing allelic counts from all heterozygous SNPs within the same haplotype by gene. Notably, the increased number of expressed heterozygous SNPs provided by imputation increased the proportion of genes assayable (≥50 reads) for gene-level ASE from 50% to ∼90%.

### Differential ASE

To distinguish somatic from physiological ASE (random monoallelic silencing, imprinting etc.) we performed differential (diff)ASE analysis. diffASE is the difference in gene-level allelic bias between the normal and the tumor expression profiles (e.g. 50:50 versus 60:40, chi-squared p = 0.1). An event was either tumor- or normal-specific, depending on which sample deviated further from 50:50 allelic expression. diffASE events where the ratio between two alleles is more skewed in the tumor than in the normal are of primary interest. The FDR generated with 10,000 permutations of ASE reads from tumor and normal samples for each gene. In cases where matched normal RNA-seq was not available, ‘tumorASE’ was assessed relative to the binomial distribution (P<0.001). The FDR was assessed with 10,000 permutations of the reads at each gene randomly drawn from all samples. To compare the outcome of diffASE and tumorASE, we filtered for genes ≥50 reads in at least half of both the tumor and normal samples. Extensive overlap between diffASE events (chi-squared P<0.001) and tumorASE (binomial distribution P<0.001), indicates that >98% of diffASE originates in tumors (Fig. 1C). Thus we included gene-level ASE from tumor RNA-seq without a matched normal sample to dramatically increase the sample size (Fig. S1A).

### Mutation calling

To identify somatic mutations, we used Varscan in conjunction with custom filters. WGS data of 12 cancer types were downloaded from GDC Legacy Archive as 1,165 matched tumor/normal BAM files (extensive sample information is available in Table S3). We required that WGS samples had matched tumor/normal files, as well as corresponding genotype array, copy-number, and tumor RNA-seq data for inclusion. Additionally, only WGS tumor/normal pairs aligned to the GRCh37 reference build (hg19) were included in our analysis. For SKCM, we used metastatic tumor samples (sample type 6 in the TCGA database), and primary tumor samples for the remaining cancer types (sample type 1). The sequence read counts at each site were obtained from WGS BAM files aligned to the GRCh37-lite human reference genome with the SAMtools (H. Li et al., 2009) mpileup. The base quality alignment (BAQ) computation of SAMtools was turned off with the parameter ‘-B’ as it is too stringent for variant calling, and reads with mapping quality > 0 was set with ‘-q 1’. Single nucleotide substitutions, insertions and deletions were simultaneously called using Varscan2 somatic caller (Koboldt et al., 2012). Data were processed using a 1,052-core Linux cluster at the High-Performance Computing Virtual Laboratory (HPCVL) (Kingston, Ontario).

Somatic mutations were focused on single-nucleotide substitutions, as well as small insertions and deletions (Fig. S2D), not structural variations. Somatic mutations generated by Varscan2 were initially filtered with two criteria: (1) a minimum read depth of 10 for both the tumor and matched normal, and (2) alleles with a mutation frequency exceeding 0.1 in tumor but less than 0.1 in matched normal (Fig. S2A). However, since ∼60% of the mutations called with this approach were rare germline SNPs (Fig. S2B; all SNPs from dbSNP150), we implemented custom filters to deplete them.

The custom filters were chosen based on optimization on 48 randomly selected BRCA tumor/normal WGS. Pearson correlation analysis was performed among 48 BRCA WGS samples to determine which Varscan parameters contributed to dbSNPs being called mutations. These included the number of normal reads, tumor reads, alternative allele frequency (AAF) in tumors (Tumor AAF), AAF in normal (Normal AAF), and Delta AAF (i.e., Tumor AAF – Normal AAF). The extent of germline SNPs contaminating Varscan-called somatic mutations was assessed as a proportion and Pearson correlation R^2^ relative to dbSNP150s from UCSC and mapped by genomic coordinates to both dbSNP alleles. This assessment was run among 48 randomly selected BRCA tumor/normal WGS. Using these metrics of contamination, each parameter (Tumor AAF, Normal AAF and Delta AAF was assessed through a range of values. This analysis revealed that two parameters, Normal AAF (R^2^=0.37, *P*<1×10 ^-4^) and Delta AAF (R^2^=-0.13, *P*<1×10 ^-4^), were primarily responsible for the high proportion of dbSNPs in Varscan.

This optimization supported use of an AAF ≤ 2% in matched normal samples since the fraction of dbSNPs increased when the AAF was higher (Fig. S2C). It also supported a requirement that the AAF exceed 10% in tumors. Finally, we required that the difference between the AAF in tumors and matched normal samples exceed 30% based on the optimization (Fig. S2B). Only 12.8% of variants passed this 30% threshold but it was effective in decreasing the fraction of dbSNPs included in the association as variants (Fig. S2C). Finally, we filtered all known dbSNPs from the mutations that passed these filters.

### Effect of copy number variations on gene-level ASE

To evaluate the effect of CNV on gene-level ASE, raw Affymetrix CNV data were downloaded from GDC Legacy Archive for 1,091 BRCA tumors. CNV data were then annotated with ‘GenomeWideSNP_6.cn.na35.annot.csv.zip’ downloaded from Affymetrix home page and mapped to Ensembl genes. Gene-level CNV signals were calculated by averaging the signals of all CNVs mapped to the gene. Finally, the somatic CNV signal for each gene was correlated with its corresponding value of gene-level allelic imbalance (reads ≥50) to determine the influence of CNV on ASE in tumors. We applied this process to 1,091 tumor ASE and 92 diffASE, calling significant associations between gene-level ASE and CNV signal when P<0.05.

To remove the confounding effect of CNVs among associations between gene-level ASE and regulatory mutations, we assessed the correlation of each gene associated in our analysis with CNV signal. We also determined the association between gene-level ASE and CNV exclusively among driver mutation carriers to differentiate the effect of driver mutations from that of CNV on gene-level ASE. We applied this filter for BRCA and other 11 cancer types, and found that none of driver genes displayed significant association (P<0.05) with CNV when only the samples containing driver mutations were considered.

To ask whether the CNV contributed to the association between mutated regulatory features and ASE of individual genes, we asked if CNV and ASE were correlated for each putative driver (Table 1 and S1). 30 of 36 drivers were not correlated with CNV. For the 7 genes where there was an association, we asked if it was dependent on CNV. The only putative driver where mutations and CNV coincided was *TSHZ2* (STAD). Hence the association between mutations and ASE occurs independent of CNV in 35/36 putative drivers. Even for *TSHZ2* (STAD), the mutations in TF binding sites and CNV only coincided in 1 tumor, and the association between mutations and ASE remained significant after excluding it (P=2.2×10 ^-5^).

### Effect of methylation on gene-level ASE

To determine the effect of methylation on gene-level ASE, we downloaded methylation Beta values for 1,091 BRCA tumors from GDC Legacy Archive. These methylation data were converted into bed format and mapped to Ensembl genes. The average methylation score was determined for each gene including a 2kb region upstream and downstream of each gene to encompass the promoter. ASE imbalance values were then correlated with average methylation levels on a gene-by-gene basis to determine the influence of methylation on gene-level ASE. This analysis was applied to all genes for 92 tumor/normal RNA-seq samples. Significant correlations between gene-level ASE and methylation Beta value were called at P<0.05.

### Selection of *cis-*regulatory features

We surveyed major *cis-*regulatory features for *cis-*regulatory variants. These included TF binding sites (Encode ChIP-seq peaks clustered V3, 2013) and CTCF binding sites (from GM12878 cell line) both obtained from the UCSC database. We derived a single track of TF binding sites by collapsing multiple ChIP-seq maps of TF binding (Table S4). We also interrogated promoters (Roadmap Project) and cancer-specific enhancers (Corces et al., 2018). These were defined based on the presence of peaks: promoters were defined as H3K4me3+ regions (signal in ≥10/127 tissues/cell types from the NIH Roadmap Epigenomics Mapping Consortium), while cancer-specific enhancers were defined by association between accessible chromatin and gene-expression changes in specific cancers (Corces et al., 2018).

### Association of mutations and gene-level ASE

Only samples with corresponding WGS, genotyping array and tumor RNA-seq data were included in the ASE-mutation association analysis. To test association between mutations and gene-level ASE across various regulatory features, these data were imported into MATLAB 2014a (The MathWorks Inc., Natick, MA, 2014).

First, the somatic mutations were mapped based on proximity to promoters and enhancers as well as TF and CTCF binding sites. Using these annotations, promoters comprise 1.6% of the genome, enhancers ∼1.4% (ranging from 0.8% in LGG to 2.7% in BRCA), TF binding sites 13.2% and CTCF binding 6.0% of the genome. Somatic mutations were binned as present (=1) or absent (=0) among the regulatory features. The overlap among these regulatory features ranges from 0.04% to 85% (Fig. S3A). For example, CTCF and TF binding sites occupy 20-70% of the enhancer feature, while the enhancers occupy <10% of the CTCF binding sites.

Second, somatic mutations, including single-nucleotide alterations, insertions and deletions were mapped to nearby genes. The genomic coordinates of each gene were defined as beginning 10kb upstream of the TSS and gene body of each gene. These settings were applied to test mutation association within each regulatory region (see Fig. S3C-D for a summary of the number of mutations mapped to the different regulatory features). Third, each gene (*i*) containing somatic mutations and also with summed heterozygous SNP allelic counts ≥50 reads was analyzed for gene-level allelic imbalance (*P*) by using the read counts of the two haplotypes (Ha, Hb) of each gene (*i*), in each sample (*n*), with the following formula, *P*_i_=|2×H _a_/(H_a_+H_b_) – 1|. The gene-level ASE varies considerably between different cancer types (Fig. S3B, binomial P-value<0.001). To increase sample size in the association test, we included all gene-level ASE, regardless of their binomial P-values.

Finally, the correlation significance (Pearson r^2^ value; coefficient of determination) between a gene’s allelic imbalance and mutations in each annotated region was determined with MATLAB. To obtain robust results we only ran the association when both mutation carriers (n>=3) and non-carriers (n>=3) had gene-level imbalance values derived from summing ≥50 reads from all heterozygous sites. Only ASE events positively correlated with mutations were retained in the analysis to focus on allelic imbalance resulting from dysregulation. All negative correlation were assigned P value =1.

We permutated the data to determine the false discovery rate (FDR) for the association between gene-level ASE and the occurrence of somatic mutations in each genomic feature. For mutations residing in specific genomic regions, all pairs of gene-level ASE and mutations were randomized 1,000 times to generate association P-values that reflect the distribution of the data. The FDR for each gene was then calculated through comparison of the actual association P-value to all genes’ minimum P-values derived from 1,000 random permutation as: sum(real P<minimum(1,000 permutations of P-values))/total number of genes. Regulatory features that could be associated with multiple genes were included in all possible associations. When independent mutations were found within the same feature of the same sample, they were collapsed to a single mutation for the association. Finally, if multiple regulatory features were enriched for ASE of the same gene, only the most significant association with the smallest FDR was retained.

### VAF

VAF was calculated independently for each mutation that we identified in WGS data as the fraction of all sequencing reads covering the variant that were mutated. VAF was z-scaled within each patient using all mutations detected in that patient in order to normalize out the effect of tumor heterogeneity. Noncoding driver and background sets were selected on basis of association with ASE. P>0.5 and FDR=1 were used as criteria for lack of significant association with ASE. Significance (2-tailed) was assessed using an unpaired t-test for unequal variance.

### Survival Analysis

Unique patients IDs with predicted noncoding drivers were extracted from Table S1 (ASE-mutation association FDR<0.5). Clinical data for TCGA was downloaded from BROAD Firehose (https://gdac.broadinstitute.org/) on April 21, 2017. Null distributions of patients were selected on basis of 1) having matching mutations in the same set of features in the same cancer type, 2) being powered to detect ASE (i.e. presence of RNA-seq data and phased SNPs), and 3) association between ASE and mutation having no significance (i.e. FDR=1). Probability of a difference in survival was assessed using the Kaplan Meier approach with censored data implemented in the MATLAB script KMPLOT (Cardillo, 2008). Since P-values are dependent on random selection of background and the limitations above imposed an upper limit on the number of background subjects that fit all criteria. Bootstrapping with 5,000 iterations was conducted to compare with P-values generated from random sets of background comparisons.

### Effect on Transcription Factor Binding Sites

The human genome was scanned using HOMER (scanMotifGenomeWide.pl using default settings for 392 motifs in the HOMER package (Heinz et al., 2010) to map putative binding sites for each factor. The default log-odds detection thresholds included with the package were used. Somatic mutations were overlapped with each motif and the difference in bit-scores (i.e. “delta-bit” using the PWM) between the reference and mutated bases was calculated, with 2 is the maximum bit-score. Delta-bit scores associated with genes under ASE versus no-ASE were compared. Only single nucleotide substitutions at positions with phastcons scores greater than 0.05 were considered for this analysis. A mutation could be considered more than once if two or more transcription factor binding motifs were present.

### Driver-ASE

We have implemented our analysis methods for gene-level ASE and somatic mutation calling into a Perl package named Driver-ASE, which is available at GitHub (https://github.com/MichealRollins-Green/Driver-ASE). All MATLAB scripts to test association between mutations and gene-level ASE are also included in Driver-ASE. All of the dependencies required to run Driver-ASE are contained in a Docker (http://www.docker.com) image found here: https://hub.docker.com/r/mikegreen24/driver-ase. Docker is required to run Driver-ASE and the instructions Docker installation can be found here: https://docs.docker.com/engine/installation. Instructions to set up a Docker image are in the description section of the Docker page.

## Data availability

Driver-ASE uses data or software provided by the following websites: UCSC Genome Browser (https://genome.ucsc.edu/cgi-bin/hgTables), Cancer Genomics Hub (https://cghub.ucsc.edu), Genomic Data Commons (https://gdc.cancer.gov), The Cancer Genome Atlas (TCGA) (http://cancergenome.nih.gov), PLINK (www.cog-genomics.org/plink), NIH Roadmap Epigenomics Mapping Consortium (www.roadmapepigenomics.org), SAMtools (www.htslib.org), overlapSelect (http://hgdownload.soe.ucsc.edu/admin/exe/linux.x86_64.v287), VarScan (http://massgenomics.org/varscan), impute2 (https://mathgen.stats.ox.ac.uk/impute/impute_v2.html) and shapeit (https://mathgen.stats.ox.ac.uk/genetics_software/shapeit/shapeit.html). All raw gene-level ASE and somatic mutations called by Varscan can be freely accessed via DRYAD (https://datadryad.org/review?doi=doi:10.5061/dryad.fp3573m).

## Acknowledgements

This study was supported by Canadian cancer CBCF grant BC-RG-15-2 (PI: Tomas Babak). No funding sources were involved in study design, data collection and interpretation, or the decision to submit the work for publication.

## Author Contributions

T.B. and B.D. designed the study. T.B, B.D., Z.C., M.V., and M.R. analyzed the data and wrote the manuscript.

## Supplementary Data

**Figure S1.**
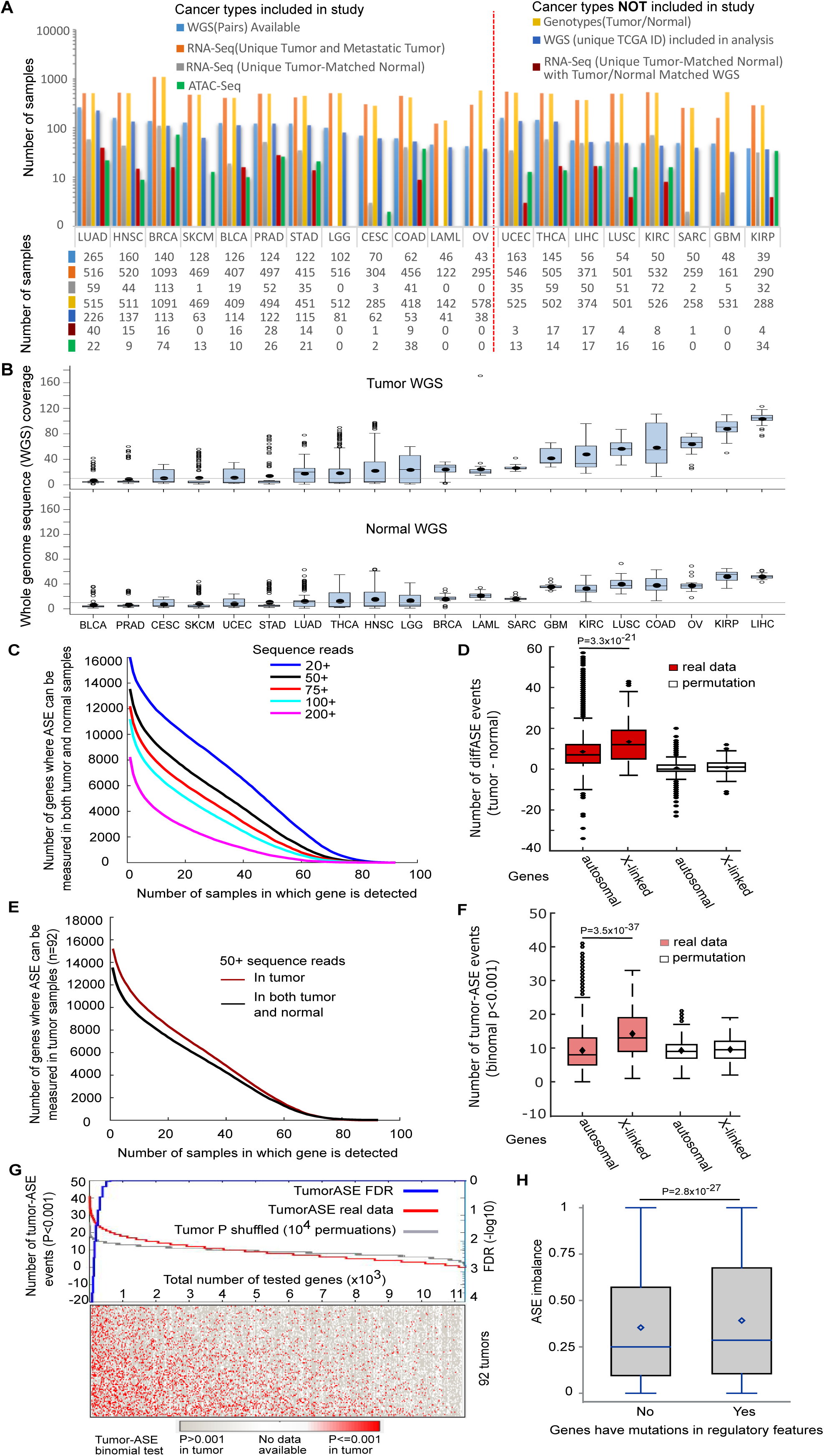
diffASE and tumorASE events exceed background at the chosen thresholds. (A) A summary of relevant TCGA samples available for analysis. (B) Whole genome sequence (WGS) coverage for tumor and matched normal across 20 cancer types (estimated based on bam size of WGS). (C) The number of genes where diffASE can be calculated per BRCA sample (n=92 tumor/normal RNA-seq) at various read thresholds. (D) X-linked diffASE events are enriched relative to autosomal events. The number of X-linked diffASE events exceeds the number of autosomal events in real, but not permuted data (P=3.3×10^-21^, ANOVA). (E) A comparison of the number of genes where diffASE and tumorASE (n=92 tumor/normal RNA-seq) can be calculated per BRCA sample at a threshold of 50 reads. (F) X-linked tumorASE events for the 92 tumor RNA-seq samples are enriched relative to autosomal events. The number of X-linked and autosomal tumorASE events (P=3.5×10^-37^, ANOVA). (G) tumorASE events in BRCA (n=92 RNA-seq) exceed the number expected by chance due to the distribution of the data. tumorASE events are those where the allelic ratio is P<0.001 using a binomial test. 241 tumorASE events with FDR<0.05 were obtained when the tumorASE events calculated with the actual sample identities were compared to the background obtained with 10,000 permutations of randomized tumor identities. The FDR reflects the proportion of permutations where the most significant tumorASE event was obtained with the actual tumor data. (H) Gene-level ASE is imbalanced among those harboring mutations in CTCF or TF binding sites, promoters, and cancer-specific enhancers (2.8×10^-27^, ANOVA, n=113 samples where tumor RNA-seq and matched tumor/normal WGS was available). Only 16 of these samples overlap with those used to compute diffASE in Fig. 1.

**Figure S2.**
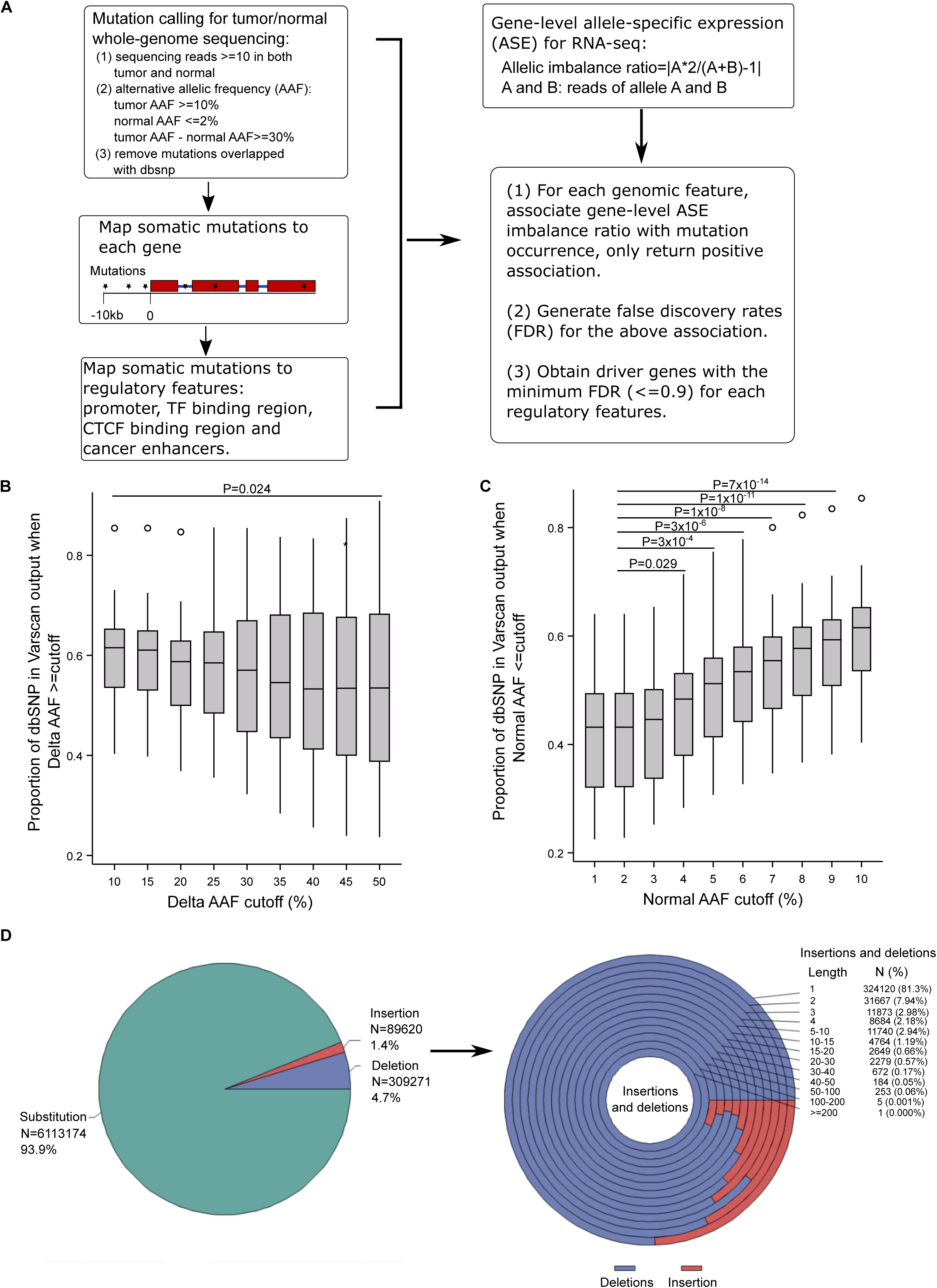
Optimized parameters deplete somatic mutation calls of rare germline SNPs. (A) A schematic of the analysis workflow used to associate allelic imbalance with mutations in regulatory regions. Our analyses identified 36 drivers at a false discovery rate (FDR)<0.9 (see Table S1 for detailed output). 12 tumors were analyzed: breast cancer (BRCA), Bladder Urothelial *Carcinoma* (BLCA), Cervical Squamous Cell *Carcinoma* (CESC), Colon Adenocarcinoma (COAD), Head and Neck Squamous Cell *Carcinoma* (HNSC), Acute Myeloid Leukemia (LAML), Low Grade Glioma (LGG), Lung Adenocarcinoma (LUAD), Ovarian cancer (OV), Prostate Adenocarcinoma (PRAD), Skin Cutaneous Melanoma (SKCM), Stomach Adenocarcinoma (STAD). (B, C) The effect of filtering parameters on the fraction of false-positive mutations (SNPs) called by Varscan was determined using BRCA WGS samples (n=48 matched tumor/normal BRCA WGS samples) (B) The proportion of SNPs among mutations called by Varscan decreases when the delta alternative allele frequency (delta AAF) between tumor and matched normal whole genome sequence (WGS) increases (≥10 versus ≥50, P=0.024, n=48 BRCA samples, Duncan’s new multiple range test). C) The proportion of false positive mutations called by Varscan (i.e. those that are actually SNPs) increases when normal AAF increases (normal AAF at a threshold of 2% was significantly different when compared to normal AAF thresholds ranging from 4% to 10%, n=48 BRCA WGS samples, Duncan’s new multiple range test). (D) The total number of somatic mutations, including substitutions, insertions, and deletions for all 12 cancer types (left panel). The insertions and deletions grouped by the length of insertion and deletion (right panel). All of these somatic mutations were obtained by applying the optimized WGS filters listed in (A).

**Figure S3.**
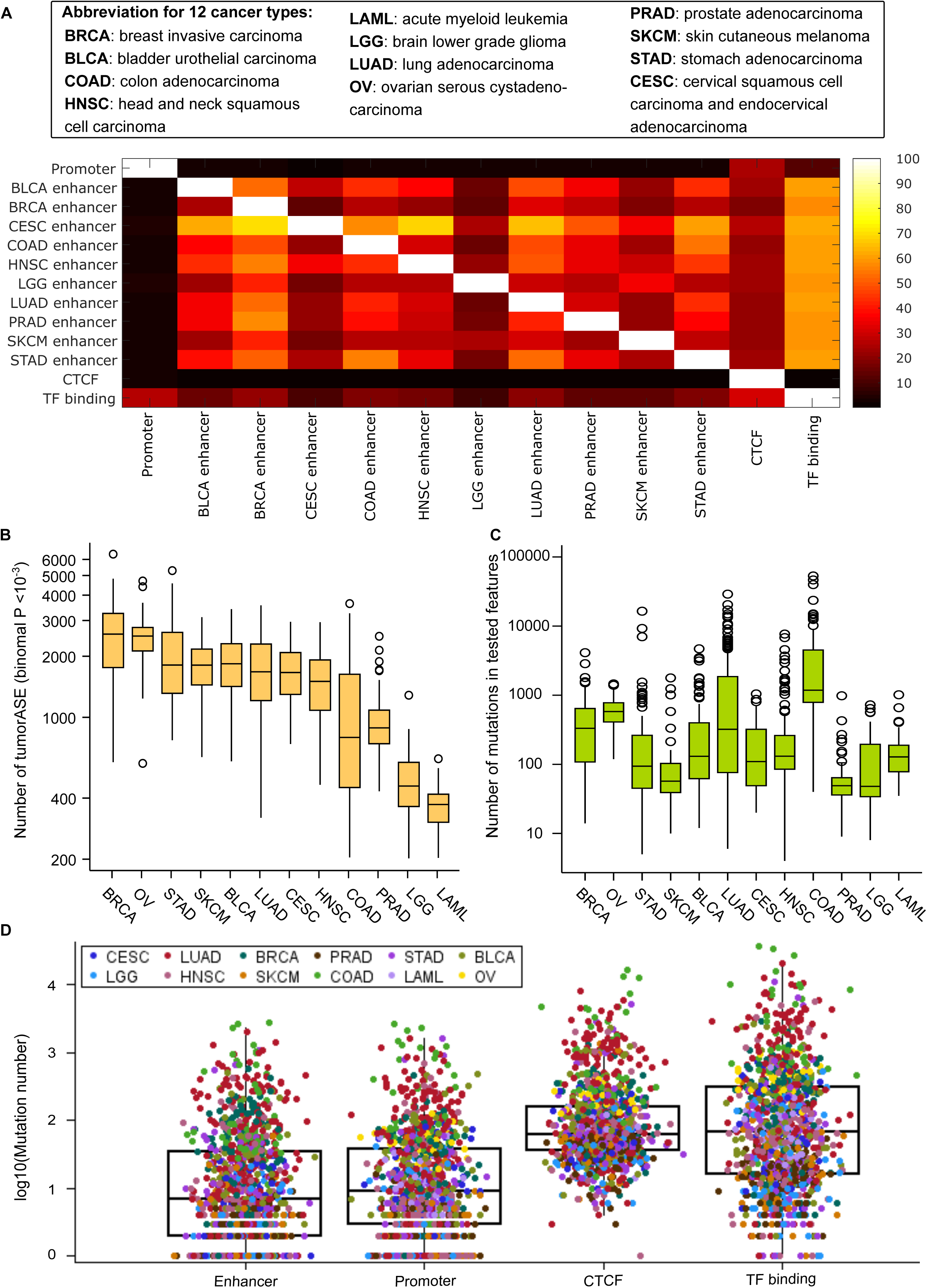
Characterizing four regulatory features and tumor ASE events, as well as mutations residing in these regulatory features, for 12 cancer types. (A) The heatmap displays the pairwise overlap among the four regulatory features analyzed: CTCF, TF binding sites (ChIP-seq), promoters, and cancer-specific enhancers. Reading to the right of the diagonal indicates how much of the feature listed in each row is overlapped by the feature listed in each column. Conversely, to the left of the diagonal indicates how much of the feature listed in each column is overlapped by the feature listed in each row. For example, 20-70% of each enhancer track is overlapped by the CTCF and TF binding sites, but these enhancers occupy <10% of the CTCF and TF binding sites. (B) The average number of tumorASE (binomial P<10^-3^) events per tumor among 1,165 tumor RNA-seq samples plotted by the 12 types of cancers. (C) The average number of somatic mutations residing in the four regulatory features among 1,165 tumor/normal WGS divided into the 12 cancer types. (D) The number of somatic mutations in each regulatory feature distinguished by different colors for each of the 12 cancer types. In each boxplot, the horizontal line represents the median. The boxes are delimited by the first and the third quartile.

### Supplementary Tables

**Table S1. Annotated catalog of the 301 potential driver mutations among 36 putative driver hotspots.**

**Table S2. A table of known non-coding drivers.**

**Table S3. Annotation of the samples used in this analysis.**

**Table S4. Annotation of the samples used to generate the meta-track of TF binding sites.**

## REFERENCES

Anand-Apte, B., Bao, L., Smith, R., Iwata, K., Olsen, B. R., Zetter, B., & Apte, S. S. (1996). A review of tissue inhibitor of metalloproteinases-3 (TIMP-3) and experimental analysis of its effect on primary tumor growth. Biochem Cell Biol, 74(6), 853–862.

Babak, T., DeVeale, B., Tsang, E. K., Zhou, Y., Li, X., Smith, K. S., … Fraser, H. B. (2015). Genetic conflict reflected in tissue-specific maps of genomic imprinting in human and mouse. Nat Genet, 47(5), 544–549. doi:10.1038/ng.3274

Bailey, M. H., Tokheim, C., Porta-Pardo, E., Sengupta, S., Bertrand, D., Weerasinghe, A., … Ding, L. (2018). Comprehensive Characterization of Cancer Driver Genes and Mutations. Cell, 174(4), 1034–1035. doi:10.1016/j.cell.2018.07.034

Beerenwinkel, N., Antal, T., Dingli, D., Traulsen, A., Kinzler, K. W., Velculescu, V. E., … Nowak, M. A. (2007). Genetic progression and the waiting time to cancer. PLoS Comput Biol, 3(11), e225. doi:10.1371/journal.pcbi.0030225

Bigas, A., & Espinosa, L. (2018). The multiple usages of Notch signaling in development, cell differentiation and cancer. Curr Opin Cell Biol, 55, 1-7. doi:10.1016/j.ceb.2018.06.010

Bononi, A., Giorgi, C., Patergnani, S., Larson, D., Verbruggen, K., Tanji, M., … Carbone, M. (2017). BAP1 regulates IP3R3-mediated Ca(2+) flux to mitochondria suppressing cell transformation. Nature, 546(7659), 549–553. doi:10.1038/nature22798

Calvo, F., Ranftl, R., Hooper, S., Farrugia, A. J., Moeendarbary, E., Bruckbauer, A., … Sahai, E. (2015). Cdc42EP3/BORG2 and Septin Network Enables Mechano-transduction and the Emergence of Cancer-Associated Fibroblasts. Cell Rep, 13(12), 2699–2714. doi:10.1016/j.celrep.2015.11.052

Cancer Genome Atlas Research, N. (2014). Comprehensive molecular profiling of lung adenocarcinoma. Nature, 511(7511), 543–550. doi:10.1038/nature13385

Cancer Genome Atlas Research, N. (2015). The Molecular Taxonomy of Primary Prostate Cancer. Cell, 163(4), 1011–1025. doi:10.1016/j.cell.2015.10.025

Cardillo, G. (2008). KMPLOT: Kaplan-Meier estimation of the survival function. doi:http://www.mathworks.com/matlabcentral/fileexchange/22293

Carter, H., Chen, S., Isik, L., Tyekucheva, S., Velculescu, V. E., Kinzler, K. W., … Karchin, R. (2009). Cancer-specific high-throughput annotation of somatic mutations: computational prediction of driver missense mutations. Cancer Res, 69(16), 6660–6667. doi:10.1158/0008-5472.CAN-09-1133

Corces, M. R., Granja, J. M., Shams, S., Louie, B. H., Seoane, J. A., Zhou, W., … Chang, H. Y. (2018). The chromatin accessibility landscape of primary human cancers. Science, 362(6413). doi:10.1126/science.aav1898

Delaneau, O., Coulonges, C., & Zagury, J. F. (2008). Shape-IT: new rapid and accurate algorithm for haplotype inference. BMC Bioinformatics, 9, 540. doi:10.1186/1471-2105-9-540

Foo, J., Liu, L. L., Leder, K., Riester, M., Iwasa, Y., Lengauer, C., & Michor, F. (2015). An Evolutionary Approach for Identifying Driver Mutations in Colorectal Cancer. PLoS Comput Biol, 11(9), e1004350. doi:10.1371/journal.pcbi.1004350

Fraser, H. B. (2011). Genome-wide approaches to the study of adaptive gene expression evolution: systematic studies of evolutionary adaptations involving gene expression will allow many fundamental questions in evolutionary biology to be addressed. Bioessays, 33(6), 469–477. doi:10.1002/bies.201000094

Fredriksson, N. J., Ny, L., Nilsson, J. A., & Larsson, E. (2014). Systematic analysis of noncoding somatic mutations and gene expression alterations across 14 tumor types. Nat Genet, 46(12), 1258–1263. doi:10.1038/ng.3141

Fu, Y., Liu, Z., Lou, S., Bedford, J., Mu, X. J., Yip, K. Y., … Gerstein, M. (2014). FunSeq2: a framework for prioritizing noncoding regulatory variants in cancer. Genome Biol, 15(10), 480. doi:10.1186/s13059-014-0480-5

Gerstung, M., Pellagatti, A., Malcovati, L., Giagounidis, A., Porta, M. G., Jadersten, M., … Boultwood, J. (2015). Combining gene mutation with gene expression data improves outcome prediction in myelodysplastic syndromes. Nat Commun, 6, 5901. doi:10.1038/ncomms6901

Heinz, S., Benner, C., Spann, N., Bertolino, E., Lin, Y. C., Laslo, P., … Glass, C. K. (2010). Simple combinations of lineage-determining transcription factors prime cis-regulatory elements required for macrophage and B cell identities. Mol Cell, 38(4), 576–589. doi:10.1016/j.molcel.2010.05.004

Hornshoj, H., Nielsen, M. M., Sinnott-Armstrong, N. A., Switnicki, M. P., Juul, M., Madsen, T., … Pedersen, J. S. (2018). Pan-cancer screen for mutations in non-coding elements with conservation and cancer specificity reveals correlations with expression and survival. NPJ Genom Med, 3, 1. doi:10.1038/s41525-017-0040-5

Howie, B. N., Donnelly, P., & Marchini, J. (2009). A flexible and accurate genotype imputation method for the next generation of genome-wide association studies. PLoS Genet, 5(6), e1000529. doi:10.1371/journal.pgen.1000529

Joberty, G., Perlungher, R. R., & Macara, I. G. (1999). The Borgs, a new family of Cdc42 and TC10 GTPase-interacting proteins. Mol Cell Biol, 19(10), 6585–6597.

Joberty, G., Perlungher, R. R., Sheffield, P. J., Kinoshita, M., Noda, M., Haystead, T., & Macara, I. G. (2001). Borg proteins control septin organization and are negatively regulated by Cdc42. Nat Cell Biol, 3(10), 861–866. doi:10.1038/ncb1001-861

Kandoth, C., McLellan, M. D., Vandin, F., Ye, K., Niu, B., Lu, C., … Ding, L. (2013). Mutational landscape and significance across 12 major cancer types. Nature, 502(7471), 333–339. doi:10.1038/nature12634

Khurana, E., Fu, Y., Chakravarty, D., Demichelis, F., Rubin, M. A., & Gerstein, M. (2016). Role of non-coding sequence variants in cancer. Nat Rev Genet, 17(2), 93–108. doi:10.1038/nrg.2015.17

Koboldt, D. C., Zhang, Q., Larson, D. E., Shen, D., McLellan, M. D., Lin, L., … Wilson, R. K. (2012). VarScan 2: somatic mutation and copy number alteration discovery in cancer by exome sequencing. Genome Res, 22(3), 568–576. doi:10.1101/gr.129684.111

Kulik, G. I., Pel’kis, F. P., & Korol, V. I. (1989). [Adaptation of the body to alkylating anti-tumor substances]. Eksp Onkol, 11(6), 34–38.

Li, D., Hallett, M. A., Zhu, W., Rubart, M., Liu, Y., Yang, Z., … Shou, W. (2011). Dishevelled-associated activator of morphogenesis 1 (Daam1) is required for heart morphogenesis. Development, 138(2), 303–315. doi:10.1242/dev.055566

Li, H., Handsaker, B., Wysoker, A., Fennell, T., Ruan, J., Homer, N., … Durbin, R. (2009). The Sequence Alignment/Map format and SAMtools. Bioinformatics, 25(16), 2078–2079.

Liu, J., Lichtenberg, T., Hoadley, K. A., Poisson, L. M., Lazar, A. J., Cherniack, A. D., … Hu, H. (2018). An Integrated TCGA Pan-Cancer Clinical Data Resource to Drive High-Quality Survival Outcome Analytics. Cell, 173(2), 400–416 e411. doi:10.1016/j.cell.2018.02.052

Liu, W., Sato, A., Khadka, D., Bharti, R., Diaz, H., Runnels, L. W., & Habas, R. (2008). Mechanism of activation of the Formin protein Daam1. PNAS, 105(1), 210–215. doi:10.1073/pnas.0707277105

Lu, X., Le Noble, F., Yuan, L., Jiang, Q., De Lafarge, B., Sugiyama, D., … Eichmann, A. (2004). The netrin receptor UNC5B mediates guidance events controlling morphogenesis of the vascular system. Nature, 432(7014), 179–186. doi:10.1038/nature03080

Mathelier, A., Lefebvre, C., Zhang, A. W., Arenillas, D. J., Ding, J., Wasserman, W. W., & Shah, S. P. (2015). Cis-regulatory somatic mutations and gene-expression alteration in B-cell lymphomas. Genome Biol, 16, 84. doi:10.1186/s13059-015-0648-7

Mayba, O., Gilbert, H. N., Liu, J., Haverty, P. M., Jhunjhunwala, S., Jiang, Z., … Zhang, Z. (2014). MBASED: allele-specific expression detection in cancer tissues and cell lines. Genome Biol, 15(8), 405. doi:10.1186/s13059-014-0405-3

Melton, C., Reuter, J. A., Spacek, D. V., & Snyder, M. (2015). Recurrent somatic mutations in regulatory regions of human cancer genomes. Nat Genet, 47(7), 710–716. doi:10.1038/ng.3332

Merid, S. K., Goranskaya, D., & Alexeyenko, A. (2014). Distinguishing between driver and passenger mutations in individual cancer genomes by network enrichment analysis. BMC Bioinformatics, 15, 308. doi:10.1186/1471-2105-15-308

Nik-Zainal, S., Davies, H., Staaf, J., Ramakrishna, M., Glodzik, D., Zou, X., … Stratton, M. R. (2016). Landscape of somatic mutations in 560 breast cancer whole-genome sequences. Nature, 534(7605), 47–54. doi:10.1038/nature17676

Ongen, H., Andersen, C. L., Bramsen, J. B., Oster, B., Rasmussen, M. H., Ferreira, P. G., … Dermitzakis, E. T. (2014). Putative cis-regulatory drivers in colorectal cancer. Nature, 512(7512), 87–90. doi:10.1038/nature13602

Piraino, S. W., & Furney, S. J. (2017). Identification of coding and non-coding mutational hotspots in cancer genomes. BMC Genomics, 18(1), 17. doi:10.1186/s12864-016-3420-9

Poulos, R. C., Sloane, M. A., Hesson, L. B., & Wong, J. W. (2015). The search for cis-regulatory driver mutations in cancer genomes. Oncotarget, 6(32), 32509–32525. doi:10.18632/oncotarget.5085

Puente, X. S., Bea, S., Valdes-Mas, R., Villamor, N., Gutierrez-Abril, J., Martin-Subero, J. I., … Campo, E. (2015). Non-coding recurrent mutations in chronic lymphocytic leukaemia. Nature, 526(7574), 519–524. doi:10.1038/nature14666

Schroeder, M. P., Rubio-Perez, C., Tamborero, D., Gonzalez-Perez, A., & Lopez-Bigas, N. (2014). OncodriveROLE classifies cancer driver genes in loss of function and activating mode of action. Bioinformatics, 30(17), i549–555. doi:10.1093/bioinformatics/btu467

Siepel, A., Bejerano, G., Pedersen, J. S., Hinrichs, A. S., Hou, M., Rosenbloom, K., … Haussler, D. (2005). Evolutionarily conserved elements in vertebrate, insect, worm, and yeast genomes. Genome Res, 15(8), 1034–1050. doi:10.1101/gr.3715005

Sjoblom, T., Jones, S., Wood, L. D., Parsons, D. W., Lin, J., Barber, T. D., … Velculescu, V. E. (2006). The consensus coding sequences of human breast and colorectal cancers. Science, 314(5797), 268–274. doi:10.1126/science.1133427

Smith, K. S., Yadav, V. K., Pedersen, B. S., Shaknovich, R., Geraci, M. W., Pollard, K. S., & De, S. (2015). Signatures of accelerated somatic evolution in gene promoters in multiple cancer types. Nucleic Acids Res, 43(11), 5307–5317. doi:10.1093/nar/gkv419

Tai-Nagara, I., Yoshikawa, Y., Numata, N., Ando, T., Okabe, K., Sugiura, Y., … Kubota, Y. (2017). Placental labyrinth formation in mice requires endothelial FLRT2/UNC5B signaling. Development, 144(13), 2392–2401. doi:10.1242/dev.149757

Verhaak, R. G., Hoadley, K. A., Purdom, E., Wang, V., Qi, Y., Wilkerson, M. D., … Cancer Genome Atlas Research, N. (2010). Integrated genomic analysis identifies clinically relevant subtypes of glioblastoma characterized by abnormalities in PDGFRA, IDH1, EGFR, and NF1. Cancer Cell, 17(1), 98–110. doi:10.1016/j.ccr.2009.12.020

Vogelstein, B., Papadopoulos, N., Velculescu, V. E., Zhou, S., Diaz, L. A., Jr., & Kinzler, K. W. (2013). Cancer genome landscapes. Science, 339(6127), 1546–1558. doi:10.1126/science.1235122

Wee, P., & Wang, Z. (2017). Epidermal Growth Factor Receptor Cell Proliferation Signaling Pathways. Cancers (Basel), 9(5). doi:10.3390/cancers9050052

Weinhold, N., Jacobsen, A., Schultz, N., Sander, C., & Lee, W. (2014). Genome-wide analysis of noncoding regulatory mutations in cancer. Nat Genet, 46(11), 1160–1165. doi:10.1038/ng.3101

Yamamoto-Hino, M., Sugiyama, T., Hikichi, K., Mattei, M. G., Hasegawa, K., Sekine, S., … et al. (1994). Cloning and characterization of human type 2 and type 3 inositol 1,4,5-trisphosphate receptors. Receptors Channels, 2(1), 9–22.

Zhang, W., Bojorquez-Gomez, A., Velez, D. O., Xu, G., Sanchez, K. S., Shen, J. P., … Ideker, T. (2018). A global transcriptional network connecting noncoding mutations to changes in tumor gene expression. Nat Genet, 50(4), 613–620. doi:10.1038/s41588-018-0091-2

