## Supplementary material for "A catalog of *cis*-regulatory mutations in 12 major cancer types": Table S1

| **Genes/Driver Region** | **Regulatory Region Affected** | **Mutation** | **Cancer Type** | **mRNA expression** | **Reference (PMID)** | **Experimentally validated** |
| --- | --- | --- | --- | --- | --- | --- |
| *CD274* | 3' UTR |  | STAD | Increase | 22190470 | Mutation at the 3'-UTR of CD274 mRNA led to CD274 overexpression by disrupting the miR-570 binding |
| *BCL6* | 3' UTR |  |  |  | 27064257 | Only analysis |
| *AFF4* | 3' UTR |  |  |  | 27064257 | Only analysis |
| *NOTCH1* | 3' UTR (miRNA binding site) |  | chronic lymphocytic leukemia |  | 26200345 | 3′ UTR mutation affect NOTCH1 expression and the time-to-treatment and survival. |
| *RPS27* | 5' UTR | Chr1: 153 963 239 C>T | SKCM | Increase | 24913145 | Only analysis |
| *MRPS31* | 5' UTR |  | SKCM |  | 26091043 | Only analysis |
| *LMO1* | 5' UTR | chr11:8289481G>A | T-cell acute lymphoblastic leukemia | Increase | 28408461 | Validated by Crisper/casp9. |
| *RP1-28O17.1* | CTCF |  | COAD |  | 26053496 | Only analysis |
| *DGKI* | CTCF |  | COAD |  | 26053496 | Only analysis |
| *RP11-248E9.7 (UP 27531bp) : RPS4XP15 (DOWN 96436bp)* | CTCF |  | COAD |  | 26053496 | Only analysis |
| *BAI3* | CTCF |  | COAD |  | 26053496 | Only analysis |
| *LINC00616* | CTCF |  | COAD |  | 26053496 | Only analysis |
| *RIMS1 (UP 9248bp) : RP11-135M8__A.1 (DOWN 110250bp)* | CTCF |  | COAD |  | 26053496 | Only analysis |
| *RPL7L1P9 (UP 28078bp) : TNS1 (DOWN 10420bp)* | CTCF |  | COAD |  | 26053496 | Only analysis |
| *RP11-99C10.1 (UP 70962bp) : CNTN5 (DOWN 4912bp)* | CTCF |  | COAD |  | 26053496 | Only analysis |
| *CTC-255N20.1* | CTCF |  | COAD |  | 26053496 | Only analysis |
| *RP11-977P2.1 (UP 263191bp) : AC012150.1 (DOWN 31761bp)* | CTCF |  | COAD |  | 26053496 | Only analysis |
| *LINC00290* | CTCF |  | COAD |  | 26053496 | Only analysis |
| *RP11-297L6.1 (UP 54797bp) : RNA5SP358 (DOWN 22587bp)* | CTCF |  | COAD |  | 26053496 | Only analysis |
| *DCLK1* | CTCF |  | COAD |  | 26053496 | Only analysis |
| *U1 (UP 10722bp) : RP11-152C15.1 (DOWN 15039bp)* | CTCF |  | COAD |  | 26053496 | Only analysis |
| *IL1RAPL2* | CTCF |  | COAD |  | 26053496 | Only analysis |
| *DYNC1I1* | CTCF |  | COAD |  | 26053496 | Only analysis |
| *RYR2* | CTCF |  | COAD |  | 26053496 | Only analysis |
| *EFCAB2* | CTCF |  | COAD |  | 26053496 | Only analysis |
| *RP11-747D18.1 (UP 27860bp) : SLITRK3 (DOWN 798bp)* | CTCF |  | COAD |  | 26053496 | Only analysis |
| *RP11-382A20.4* | CTCF |  | COAD |  | 26053496 | Only analysis |
| *DHX35 (UP 41641bp) : RP4-705O1.1 (DOWN 132413bp)* | CTCF |  | COAD |  | 26053496 | Only analysis |
| *AP4B1-AS1* | CTCF |  | COAD |  | 26053496 | Only analysis |
| *Z73964.1 (UP 7177bp) : RAB9B (DOWN 4637bp)* | CTCF |  | COAD |  | 26053496 | Only analysis |
| *KYNU* | CTCF |  | COAD |  | 26053496 | Only analysis |
| *MYO16* | CTCF |  | COAD |  | 26053496 | Only analysis |
| *RP11-348F1.2* | CTCF |  | COAD |  | 26053496 | Only analysis |
| *DCAF8L2* | CTCF |  | COAD |  | 26053496 | Only analysis |
| *CRISP1* | CTCF |  | COAD |  | 26053496 | Only analysis |
| *OR4S1 (UP 806bp) : OR4C3 (DOWN 16962bp)* | CTCF |  | COAD |  | 26053496 | Only analysis |
| *RAB40A (UP 14105bp) : TCEAL4 (DOWN 42637bp)* | CTCF |  | COAD |  | 26053496 | Only analysis |
| *INPP4B* | CTCF |  | COAD |  | 26053496 | Only analysis |
| *SCGB1D5P (UP 19701bp) : RP11-443J23.1 (DOWN 10330bp)* | CTCF |  | COAD |  | 26053496 | Only analysis |
| *U6 (UP 45460bp) : AC104441.1 (DOWN 585047bp)* | CTCF |  | COAD |  | 26053496 | Only analysis |
| *RGS21 (UP 78749bp) : AL136987.1 (DOWN 45094bp)* | CTCF |  | COAD |  | 26053496 | Only analysis |
| *SPTLC3* | CTCF |  | COAD |  | 26053496 | Only analysis |
| *AC007189.2 (UP 30111bp) : FSHR (DOWN 15516bp)* | CTCF |  | COAD |  | 26053496 | Only analysis |
| *LSAMP* | CTCF |  | COAD |  | 26053496 | Only analysis |
| *CHRDL1 (UP 51451bp) : PAK3 (DOWN 96776bp)* | CTCF |  | COAD |  | 26053496 | Only analysis |
| *AC009264.1* | CTCF |  | COAD |  | 26053496 | Only analysis |
| *RP11-202G18.1 (UP 14690bp) : OR2K2 (DOWN 27957bp)* | CTCF |  | COAD |  | 26053496 | Only analysis |
| *RP11-665G4.1* | CTCF |  | COAD |  | 26053496 | Only analysis |
| *ZNF217* | CTCF |  | BRCA | Increase | 29354286 | Only analysis |
| *MAPRE* | CTCF |  |  |  | 29354286 | Only analysis |
| *DPYSL5* | CTCF |  |  |  | 29354286 | Only analysis |
| *PTPRK* | CTCF |  | LIHC | Increase | 29354386 | Only analysis |
| *CDC6* | CTCF |  | BRCA | Increase | 29354386 | Only analysis |
| *PAX5* | Enhancer |  | CLL |  | 26200345 | Enhancer mutation correlated with PAX5 expression, validated by CRISPR/Cas9 experiment. |
| *TAL1* | Enhancer |  | T-ALL (T-cell acute lymphoblastic leukemia) |  | 25394790 | Mutation in the aberrantly formed super-enhancer drives monoallelic TAL1 expression, which is validated by ChIP-seq. |
| *TCF3* | Enhancer |  | B-cell lymphoma |  | 25607463 | Only analysis |
| *SP1* | Enhancer |  | B-cell lymphoma |  | 25607463 | Only analysis |
| *IKZF1* | Enhancer |  | B-cell lymphoma |  | 25607463 | Only analysis |
| *APOH* | Enhancer |  |  | Decrease | 27064257 | Only analysis |
| *PRKCA* | Enhancer |  |  | Increase | 27064257 | Only analysis |
| *TOX3* | Enhancer |  |  |  | 29354286 | Only analysis |
| *CCND1* | Enhancer | Chr11: HIF-binding motif at 11q13.3 | KIRC (renal cancer) | Increase | 22406644 | The enhancer affects the allelic imbalance of cyclin D1 expression. |
| *GATA3* | intron; disrupt the acceptor splice site | indel | BRCA |  | 29354286 | Only analysis |
| *MALAT1* | lncRNA |  | BRCA, LUAD, BLCA |  | 28128360, 23153939 | Gene over-expression experiment in 22722193; gene knockdown experiment in 23153939. |
| *NEAT1* | lncRNA |  | Multiple tumor types |  | 28128360 | Only analysis |
| *SAMMSON* | lncRNA |  | STAD |  | 28128360 | Only analysis |
| *NFKBIZ* | miRNA binding site |  | diffuse large B-cell lymphoma |  | 30275490 | 3' UTR mutations altered the RNA structure. |
| *WDR74* | Promoter |  | Multiple tumor types |  | 25261935 | Only analysis |
| *NFKBIE* | Promoter | Chr6: 44 233 400 C>T (clustered C>T from chr6: 44 233 379 to –44 233 439) | SKCM | Unknown | 26343386 | Only analysis |
| *TFPI2* | Promoter |  |  | Decrease | 27064257 | Only analysis |
| *MED16* | Promoter |  |  |  | 27064257 | Only analysis |
| *WDR4* | Promoter |  |  |  | 27064257 | Only analysis |
| *LHX8* | Promoter |  | pancreatic ductal adenocarcinoma |  | 28481342 | Only analysis |
| *BMP7* | Promoter |  | pancreatic ductal adenocarcinoma |  | 28481342 | Only analysis |
| *LINC1194* | Promoter |  | pancreatic ductal adenocarcinoma |  | 28481342 | Only analysis |
| *LHX8* | Promoter |  | pancreatic ductal adenocarcinoma |  | 28481342 | Only analysis |
| *DUSP22* | Promoter |  | pancreatic ductal adenocarcinoma |  | 28481342 | Only analysis |
| *REREP3* | Promoter |  | pancreatic ductal adenocarcinoma |  | 28481342 | Only analysis |
| *LMX1B* | Promoter |  | pancreatic ductal adenocarcinoma |  | 28481342 | Only analysis |
| *PAX6* | Promoter |  | pancreatic ductal adenocarcinoma |  | 28481342 | Only analysis |
| *ZIC4* | Promoter |  | pancreatic ductal adenocarcinoma |  | 28481342 | Only analysis |
| *FANK1* | Promoter |  | pancreatic ductal adenocarcinoma |  | 28481342 | Only analysis |
| *ST8S1A4* | Promoter |  | pancreatic ductal adenocarcinoma |  | 28481342 | Only analysis |
| *MIR21* | Promoter |  | pancreatic ductal adenocarcinoma |  | 28481342 | Only analysis |
| *VMP1* | Promoter |  | pancreatic ductal adenocarcinoma |  | 28481342 | Only analysis |
| *DMRTA2* | Promoter |  | pancreatic ductal adenocarcinoma |  | 28481342 | Only analysis |
| *VAX2* | Promoter |  | pancreatic ductal adenocarcinoma |  | 28481342 | Only analysis |
| *ZIC4* | Promoter |  | pancreatic ductal adenocarcinoma |  | 28481342 | Only analysis |
| *DUSP22* | Promoter |  | pancreatic ductal adenocarcinoma |  | 28481342 | Only analysis |
| *MALAT1* | Promoter |  | pancreatic ductal adenocarcinoma |  | 28481342 | Only analysis |
| *ZNF595* | Promoter |  | pancreatic ductal adenocarcinoma |  | 28481342 | Only analysis |
| *ZNF718* | Promoter |  | pancreatic ductal adenocarcinoma |  | 28481342 | Only analysis |
| *CDH15* | Promoter |  | pancreatic ductal adenocarcinoma |  | 28481342 | Only analysis |
| *CDH8* | Promoter |  | pancreatic ductal adenocarcinoma |  | 28481342 | Only analysis |
| *PCF11* | Promoter |  |  |  | 29354286 | Only analysis |
| *ETV1* | Promoter |  | COAD |  | 30224643 | Reduction of endogenous ETV1 level using siRNA was associated with decreased cell viability and cell proliferation. |
| *OXNAD1* | Promoter | Chr3: 16 306 504 C>T Chr3: 16 306 505 C>T/A | SKCM | Increase | 26416425 | Reporter assays carried out in one melanoma cell line for DPH3 and OXNAD1 orientations showed statistically significant increased promoter activity due to -8/-9CC > TT tandem mutations; although, no effect of the mutations on DPH3 and OXNAD1 transcription in tumors was observed. |
| *DPH3* | Promoter | Chr3: 16 306 504 C>T Chr3: 16 306 505 C>T/A | SKCM | Increase | 26416425 | Reporter assays carried out in one melanoma cell line for DPH3 and OXNAD1 orientations showed statistically significant increased promoter activity due to -8/-9CC > TT tandem mutations; although, no effect of the mutations on DPH3 and OXNAD1 transcription in tumors was observed. |
| *NDUFB9* | Promoter | Chr8: 125 551 344 C>T | SKCM, KIRC |  | 26082173 | Validated by reporter gene assay. |
| *TERT* | Promoter | Chr5: 1 295 250 C>T; Chr5: 1 295 250 C>T | Most tumor types | Increase | 23348503, 23348506, 28230921, 23530248, 25574106, 23887589 | Validated by reporter gene assays, caused up to twofold increase in transcription of TERT. |
| *SDHD* | Promoter | Chr11: 111 957 523 C>T; Chr11: 111 957 541 C>T | SKCM | Decrease | 28108517, | Validated by reporter gene assays. |
| *FOXA1* | Promoter |  | BRCA | Increase | 28658208 | Validated in reporter assays. |
| *RMRP* | Promoter |  | BRCA |  | 28658208 | Validated in reporter assays. |
| *NEAT1* | Promoter |  | BRCA |  | 28658208 | Validated in reporter assays. |
| *TBC1D12* | Promoter |  | BRCA |  | 28658208 | Validated in reporter assays. |
| *ZNF143* | Promoter |  | BRCA |  | 28658208 | Validated in reporter assays. |
| *RMRP*/CCDC107 | Promoter |  | BRCA |  | 28658208 | Validated in reporter assays. |
| *ALDOA* | Promoter |  | BRCA |  | 28658208 | Validated in reporter assays. |
| *LEPROTL1* | Promoter |  | BRCA |  | 28658208 | Validated in reporter assays. |
| *CITED2* | Promoter |  | BRCA |  | 28658208 | Validated in reporter assays. |
| *CTNNB1* | Promoter |  | BRCA |  | 28658208 | Validated in reporter assays. |
| *NEAT1* |  |  | Liver cancer |  | 27064257 | Only analysis |
| *MALAT1* |  |  |  |  | 27064257 | Only analysis |
| *CDH10* |  |  | Pan-cancer |  | 29354286 | Only analysis |
| *CDC6* |  |  |  |  | 29354286 | Only analysis |
