## Supplementary material for "A catalog of *cis*-regulatory mutations in 12 major cancer types": Table S2

| 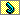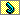**Cancer** | **Regulatory feature** | **Gene symbol** | **Sample IDs** | **Mutation list (hg38)** |
| --- | --- | --- | --- | --- |
| BLCA | TF binding | *SFMBT2* | TCGA-DK-A1A7 \| TCGA-DK-A1AE \| TCGA-DK-A2HX \| TCGA-DK-A1A3 | chr10:7281642( C->G ) \| chr10:7268020( C->G ) \| chr10:7264485( C->T ) \| chr10:7412633( G->A ) |
| BLCA | TF binding | *SLC39A11* | TCGA-BT-A20Q \| TCGA-BT-A3PH \| TCGA-CF-A27C \| TCGA-DK-A1A7 \| TCGA-DK-A3IT \| TCGA-FD-A3SO \| TCGA-BL-A3JM | chr17:72775507( G->A ) \| chr17:72666366( C->G ), chr17:72666416( C->G ), chr17:72667273( C->G ) \| chr17:72702960( G->A ) \| chr17:72727107( A->C ) \| chr17:72661785( G->T ), chr17:72661982( G->C ) \| chr17:72725546( A->C ) \| chr17:72707739( C->G ) |
| BLCA | TF binding | *TPCN1* | TCGA-BT-A3PH \| TCGA-DK-A1A6 \| TCGA-BL-A0C8 | chr12:113246752( G->A ) \| chr12:113267893( C->T ) \| chr12:113276502( A->G ) |
| BRCA | Cancer-specific enhancer | *CDC42EP3* | TCGA-AN-A04D \| TCGA-EW-A1J5 \| TCGA-A7-A13D | chr2:37684074( A->C ) \| chr2:37683992( G->C ) \| chr2:37617798( C->G ) |
| BRCA | CTCF | *DAAM1* | TCGA-AO-A0J6 \| TCGA-AO-A124 \| TCGA-B6-A0I2 \| TCGA-EW-A1PH \| TCGA-AO-A0J2 | chr14:59258774( G->T ), chr14:59264283( G->T ) \| chr14:59330300( A->C ) \| chr14:59278659( A->G ) \| chr14:59250581( C->-T ) \| chr14:59292921( T->C ) |
| BRCA | Cancer-specific enhancer | *EGFR* | TCGA-A8-A07I \| TCGA-A8-A08L \| TCGA-AR-A256 \| TCGA-B6-A0I1 \| TCGA-E2-A109 \| TCGA-E2-A14P \| TCGA-EW-A1J5 \| TCGA-EW-A1PC \| TCGA-GI-A2C9 \| TCGA-A2-A04T | chr7:55181853( C->T ) \| chr7:55133770( G->A ) \| chr7:55006831( G->A ) \| chr7:55090003( C->T ) \| chr7:55097001( T->A ) \| chr7:55080777( C->G ) \| chr7:55044839( G->C ) \| chr7:55066699( T->C ) \| chr7:55133739( T->A ) \| chr7:55181385( C->G ) |
| BRCA | TF binding | *ITPR3* | TCGA-A8-A08L \| TCGA-AO-A03N \| TCGA-C8-A12L \| TCGA-EW-A1J5 \| TCGA-A2-A0EY | chr6:33695721( C->T ) \| chr6:33668064( G->A ) \| chr6:33618415( A->G ) \| chr6:33691995( G->C ) \| chr6:33634084( G->C ) |
| BRCA | CTCF | *NOTCH1* | TCGA-AR-A256 \| TCGA-B6-A0IJ \| TCGA-GM-A3XL \| TCGA-A7-A13D | chr9:136538423( G->A ) \| chr9:136538798( G->C ) \| chr9:136543498( G->C ) \| chr9:136546832( A->G ) |
| BRCA | Cancer-specific enhancer | *TIMP3* | TCGA-A8-A08L \| TCGA-A8-A094 \| TCGA-AN-A0AT \| TCGA-AO-A0J6 \| TCGA-AO-A124 \| TCGA-E2-A15K \| TCGA-EW-A1J5 \| TCGA-A2-A0D0 | chr22:32812606( C->G ) \| chr22:32855080( G->A ) \| chr22:32843654( A->G ) \| chr22:32819786( C->A ) \| chr22:32810033( A->T ) \| chr22:32849480( G->A ) \| chr22:32852854( C->T ) \| chr22:32818293( G->A ) |
| BRCA | Promoter | *UNC5B* | TCGA-AO-A0J4 \| TCGA-D8-A27F \| TCGA-E2-A1LG \| TCGA-A8-A08L | chr10:71216278( C->T ) \| chr10:71212987( T->-GGGGGCCCGGAGCGGAGCTC ) \| chr10:71216508( G->A ) \| chr10:71212358( G->A ) |
| CESC | TF binding | *SULF1* | TCGA-DG-A2KJ \| TCGA-FU-A3TX \| TCGA-C5-A1M9 | chr8:69614558( G->A ) \| chr8:69489264( T->A ), chr8:69624474( G->A ) \| chr8:69458121( C->T ) |
| COAD | Promoter | *COL4A1* | TCGA-AA-3977 \| TCGA-CA-6718 \| TCGA-AA-3516 | chr13:110305196( C->T ) \| chr13:110309939( C->T ) \| chr13:110306834( C->T ), chr13:110308082( G->T ) |
| COAD | Promoter | *GSTA4* | TCGA-AA-3555 \| TCGA-D5-6540 \| TCGA-AA-3516 | chr6:52996489( T->C ) \| chr6:52995013( T->-A ) \| chr6:52994968( C->A ) |
| COAD | Cancer-specific enhancer | *ME3* | TCGA-A6-5659 \| TCGA-AA-3516 \| TCGA-AA-3555 \| TCGA-AA-3977 \| TCGA-AA-A01X \| TCGA-CA-6718 \| TCGA-A6-5656 | chr11:86582920( A->G ) \| chr11:86663831( C->-A ) \| chr11:86663912( C->A ) \| chr11:86574140( T->G ) \| chr11:86593091( G->A ) \| chr11:86464801( G->T ) \| chr11:86583029( G->C ) |
| HNSC | Promoter | *WLS* | TCGA-CN-6011 \| TCGA-DQ-5629 \| TCGA-CN-5374 | chr1:68229338( A->G ) \| chr1:68231897( C->T ) \| chr1:68231223( G->C ) |
| LAML | CTCF | *CBLB* | TCGA-AB-2982 \| TCGA-AB-2992 \| TCGA-AB-2999 \| TCGA-AB-2978 | chr3:105658778( C->A ) \| chr3:105845516( G->T ) \| chr3:105772506( T->A ) \| chr3:105685065( T->C ) |
| LUAD | Cancer-specific enhancer | *ATXN1* | TCGA-53-7624 \| TCGA-53-7626 \| TCGA-55-8085 \| TCGA-62-A46O \| TCGA-67-3771 \| TCGA-71-6725 \| TCGA-78-7152 \| TCGA-86-8358 \| TCGA-35-5375 | chr6:16690681( C->G ), chr6:16726488( G->A ) \| chr6:16713103( T->A ), chr6:16741145( G->C ) \| chr6:16689408( G->A ) \| chr6:16359044( C->A ) \| chr6:16677245( C->A ) \| chr6:16335090( A->+T ) \| chr6:16421511( C->G ) \| chr6:16331294( C->A ) \| chr6:16420010( G->A ) |
| LUAD | Cancer-specific enhancer | *CACNA1H* | TCGA-53-7624 \| TCGA-55-8510 \| TCGA-55-8619 \| TCGA-69-8255 \| TCGA-78-7149 \| TCGA-78-7150 \| TCGA-44-7670 | chr16:1199049( A->C ) \| chr16:1163541( G->T ) \| chr16:1199150( T->C ) \| chr16:1199125( A->C ) \| chr16:1199034( A->C ) \| chr16:1199132( T->C ) \| chr16:1182631( T->-C ) |
| LUAD | Cancer-specific enhancer | *COL4A1* | TCGA-73-4659 \| TCGA-86-8358 \| TCGA-50-5930 | chr13:110209932( A->T ) \| chr13:110206433( C->T ) \| chr13:110197311( C->A ) |
| LUAD | TF binding | *ERI2* | TCGA-55-5899 \| TCGA-73-4670 \| TCGA-44-7660 | chr16:20804758( G->T ) \| chr16:20806812( A->T ) \| chr16:20806778( G->T ) |
| LUAD | Promoter | *GATA3* | TCGA-62-A46O \| TCGA-78-7155 \| TCGA-91-8499 \| TCGA-53-7624 | chr10:8049397( G->T ) \| chr10:8057026( G->T ) \| chr10:8060487( G->T ) \| chr10:8059651( T->G ) |
| LUAD | TF binding | *ITGAE* | TCGA-50-5931 \| TCGA-50-6590 \| TCGA-55-8507 \| TCGA-62-A46O \| TCGA-64-1678 \| TCGA-78-7149 \| TCGA-78-7150 \| TCGA-78-7155 \| TCGA-86-8358 \| TCGA-91-6831 \| TCGA-44-7670 | chr17:3754897( G->A ) \| chr17:3763693( G->C ) \| chr17:3723203( T->C ) \| chr17:3723239( C->A ) \| chr17:3760032( G->T ) \| chr17:3738574( G->T ) \| chr17:3770919( G->A ) \| chr17:3724822( G->T ) \| chr17:3798715( C->A ) \| chr17:3770895( G->A ) \| chr17:3770757( C->A ) |
| LUAD | Cancer-specific enhancer | *JCAD* | TCGA-44-2659 \| TCGA-44-7670 \| TCGA-50-5931 \| TCGA-55-A4DF \| TCGA-62-A46P \| TCGA-67-3771 \| TCGA-78-7155 \| TCGA-44-2656 | chr10:29995077( G->A ) \| chr10:29996123( C->A ) \| chr10:29996155( G->T ) \| chr10:30002669( C->T ) \| chr10:30136118( G->C ) \| chr10:29958117( T->C ) \| chr10:29947761( C->A ) \| chr10:30152108( C->T ) |
| LUAD | CTCF | *LPAR1* | TCGA-44-2655 \| TCGA-44-2659 \| TCGA-44-7669 \| TCGA-44-7670 \| TCGA-44-8117 \| TCGA-44-8120 \| TCGA-49-4512 \| TCGA-50-5939 \| TCGA-55-8507 \| TCGA-55-A48X \| TCGA-55-A4DF \| TCGA-62-8399 \| TCGA-62-A46P \| TCGA-67-3771 \| TCGA-73-4670 \| TCGA-75-6214 \| TCGA-75-7031 \| TCGA-78-7156 \| TCGA-78-7535 \| TCGA-78-7536 \| TCGA-86-8673 \| TCGA-97-7937 \| TCGA-97-8179 \| TCGA-05-4250 | chr9:111010999( C->A ) \| chr9:110940880( G->A ), chr9:110963945( C->A ), chr9:111020369( C->A ) \| chr9:111003737( C->T ) \| chr9:111013977( C->A ) \| chr9:110973103( C->A ) \| chr9:110949032( C->A ), chr9:110996363( C->T ), chr9:111018111( T->C ) \| chr9:110935413( T->C ) \| chr9:110884715( T->A ) \| chr9:110910586( C->A ), chr9:110951168( G->T ), chr9:111013061( C->A ), chr9:111015747( G->T ) \| chr9:110989248( G->T ) \| chr9:110911450( G->A ) \| chr9:110875233( C->T ), chr9:110922356( C->A ) \| chr9:110933516( C->A ) \| chr9:110924163( C->A ), chr9:111004996( A->G ) \| chr9:110882021( C->A ) \| chr9:110883810( G->A ), chr9:110932318( G->T ), chr9:110948169( T->A ) \| chr9:110900614( G->C ), chr9:110977207( G->T ) \| chr9:110901165( C->T ), chr9:110904886( C->A ), chr9:110963525( C->T ) \| chr9:110956980( A->G ) \| chr9:110990680( A->C ), chr9:110998306( C->T ) \| chr9:110958761( G->A ) \| chr9:110919593( C->A ), chr9:110996676( T->C ) \| chr9:110895881( C->A ), chr9:110895951( C->A ) \| chr9:111026284( T->A ) |
| LUAD | Cancer-specific enhancer | *PIP5K1B* | TCGA-55-8507 \| TCGA-78-7150 \| TCGA-78-7155 \| TCGA-86-7955 \| TCGA-44-7660 | chr9:68795123( G->T ) \| chr9:68749892( C->G ) \| chr9:68714716( C->A ) \| chr9:68740078( A->G ) \| chr9:68731967( G->T ) |
| LUAD | CTCF | *SEMA4D* | TCGA-05-4397 \| TCGA-38-4630 \| TCGA-44-2659 \| TCGA-44-3919 \| TCGA-44-7669 \| TCGA-44-7670 \| TCGA-49-4512 \| TCGA-49-6743 \| TCGA-50-5931 \| TCGA-50-6590 \| TCGA-55-7281 \| TCGA-55-7570 \| TCGA-55-8208 \| TCGA-55-8507 \| TCGA-62-8399 \| TCGA-62-A46P \| TCGA-62-A470 \| TCGA-67-3771 \| TCGA-69-8255 \| TCGA-73-4659 \| TCGA-78-7158 \| TCGA-86-7955 \| TCGA-86-8358 \| TCGA-91-6847 \| TCGA-95-7948 \| TCGA-05-4250 | chr9:89374260( C->T ), chr9:89385157( T->G ), chr9:89411782( C->A ), chr9:89414795( C->A ) \| chr9:89430198( G->C ) \| chr9:89427984( C->A ) \| chr9:89361574( T->G ), chr9:89445305( T->G ) \| chr9:89394173( C->A ), chr9:89396441( C->A ), chr9:89396442( C->A ) \| chr9:89394063( T->A ), chr9:89409793( G->C ), chr9:89409938( G->C ), chr9:89425771( A->T ), chr9:89463920( C->A ) \| chr9:89450718( T->G ) \| chr9:89404840( T->C ) \| chr9:89372328( A->G ) \| chr9:89414560( C->A ) \| chr9:89396437( C->A ) \| chr9:89408011( C->G ) \| chr9:89372142( T->G ) \| chr9:89454619( C->A ), chr9:89460342( C->A ) \| chr9:89465278( C->A ) \| chr9:89385281( C->A ) \| chr9:89424005( T->C ) \| chr9:89384536( G->A ), chr9:89412910( T->A ), chr9:89414486( T->A ) \| chr9:89419677( C->A ) \| chr9:89429124( C->A ) \| chr9:89382859( C->A ) \| chr9:89482456( C->A ) \| chr9:89399745( C->T ), chr9:89445005( C->A ), chr9:89454883( C->A ) \| chr9:89457123( A->C ) \| chr9:89487771( G->A ) \| chr9:89383136( A->G ), chr9:89446179( A->C ) |
| LUAD | TF binding | *SND1* | TCGA-38-4631 \| TCGA-44-2656 \| TCGA-44-4112 \| TCGA-44-5643 \| TCGA-44-7660 \| TCGA-44-7670 \| TCGA-50-5930 \| TCGA-53-7624 \| TCGA-55-5899 \| TCGA-62-A46O \| TCGA-64-5781 \| TCGA-67-3771 \| TCGA-69-8255 \| TCGA-75-6214 \| TCGA-75-7031 \| TCGA-78-7143 \| TCGA-78-7155 \| TCGA-78-8640 \| TCGA-86-7955 \| TCGA-86-8358 \| TCGA-91-A4BC \| TCGA-97-7937 \| TCGA-J2-A4AD \| TCGA-05-4250 | chr7:127865628( G->T ) \| chr7:127654838( G->A ) \| chr7:127844920( G->T ), chr7:127844921( G->T ) \| chr7:127651997( C->A ) \| chr7:127678449( T->A ), chr7:127845051( T->C ) \| chr7:128055886( G->T ), chr7:128065935( G->A ) \| chr7:127736660( G->T ) \| chr7:127902189( G->C ) \| chr7:127651964( C->A ) \| chr7:127683507( G->A ) \| chr7:127955514( G->T ), chr7:127998939( G->C ) \| chr7:127822876( A->G ) \| chr7:127998953( G->T ) \| chr7:127850487( G->T ) \| chr7:127707531( A->T ) \| chr7:127705497( T->C ) \| chr7:127892039( G->T ), chr7:127908238( A->+T ) \| chr7:127652132( G->T ), chr7:127908484( C->G ) \| chr7:127895637( G->A ) \| chr7:127684361( G->C ) \| chr7:127739047( G->T ) \| chr7:128067276( C->T ) \| chr7:127777361( G->T ) \| chr7:128047218( A->C ) |
| LUAD | Cancer-specific enhancer | *TBCD* | TCGA-55-A4DF \| TCGA-78-7155 \| TCGA-MP-A5C7 \| TCGA-05-4420 | chr17:82894131( G->A ) \| chr17:82861153( G->C ) \| chr17:82876350( G->A ) \| chr17:82874538( C->T ) |
| LUAD | Cancer-specific enhancer | *TRIM2* | TCGA-55-1596 \| TCGA-55-A4DF \| TCGA-62-A46O \| TCGA-62-A46P \| TCGA-64-1678 \| TCGA-86-7955 \| TCGA-86-8358 \| TCGA-35-5375 | chr4:153217519( T->A ) \| chr4:153144226( G->C ), chr4:153217798( C->G ) \| chr4:153128118( T->C ) \| chr4:153274978( G->T ) \| chr4:153223006( G->T ) \| chr4:153134817( A->G ) \| chr4:153223005( C->T ), chr4:153223148( G->T ) \| chr4:153205252( G->A ) |
| OV | TF binding | *PKNOX2* | TCGA-24-1103 \| TCGA-24-1548 \| TCGA-36-1571 \| TCGA-23-1110 | chr11:125276797( G->T ) \| chr11:125376291( G->C ) \| chr11:125188094( A->C ) \| chr11:125163735( G->A ) |
| OV | CTCF | *SLC27A1* | TCGA-24-1557 \| TCGA-24-2290 \| TCGA-25-1319 \| TCGA-13-0890 | chr19:17461705( G->T ), chr19:17461706( G->T ) \| chr19:17503231( G->T ) \| chr19:17467268( G->T ) \| chr19:17463073( G->T ) |
| SKCM | CTCF | *KANSL1* | TCGA-FS-A1ZZ \| TCGA-GN-A262 \| TCGA-FS-A1ZF | chr17:46199069( C->A ) \| chr17:46185584( G->A ), chr17:46186359( G->T ), chr17:46189165( G->A ), chr17:46192360( G->A ), chr17:46193443( A->G ), chr17:46199899( A->C ) \| chr17:46186359( G->T ) |
| STAD | CTCF | *ARSB* | TCGA-D7-6815 \| TCGA-HU-A4H0 \| TCGA-BR-4280 | chr5:78894452( T->G ) \| chr5:78888943( C->T ) \| chr5:78906862( A->-T ) |
| STAD | Cancer-specific enhancer | *CCND2* | TCGA-F1-6177 \| TCGA-HF-7136 \| TCGA-CG-4442 | chr12:4279234( G->T ) \| chr12:4308008( C->G ) \| chr12:4279400( T->C ) |
| STAD | CTCF | *PAG1* | TCGA-CG-4472 \| TCGA-D7-6520 \| TCGA-F1-6177 \| TCGA-BR-8373 | chr8:80989949( A->C ), chr8:81100587( A->C ) \| chr8:81111576( C->G ) \| chr8:81089577( A->G ), chr8:81097722( A->-AAAG ) \| chr8:81113156( T->+AC ) |
| STAD | TF binding | *PBX1* | TCGA-CG-4465 \| TCGA-D7-6822 \| TCGA-F1-6177 \| TCGA-BR-4267 | chr1:164600784( C->T ) \| chr1:164592116( G->C ) \| chr1:164771569( C->A ), chr1:164831220( T->C ) \| chr1:164595500( A->G ) |
| STAD | TF binding | *TSHZ2* | TCGA-D7-6525 \| TCGA-D7-6822 \| TCGA-F1-6177 \| TCGA-D7-5578 | chr20:53142425( A->G ) \| chr20:53489700( A->G ) \| chr20:53003896( C->-T ), chr20:53070773( T->C ), chr20:53413396( G->A ) \| chr20:53149222( C->A ) |
