## Supplementary material for "A catalog of *cis*-regulatory mutations in 12 major cancer types": Table S3

| **Cancer Type** | **Individual** | **File ID (Tumor)** | **File ID (matched normal)** | **Number of Somatic Variants** |
| --- | --- | --- | --- | --- |
| BRCA | TCGA-A1-A0SM | d1b482b0-0e66-4620-bc89-bfc93541735d | 96a41a4f-52b2-4699-9a58-6e8e2be5e409 | 925 |
| BRCA | TCGA-A2-A04P | 680a94d4-95ae-4e50-b380-28f9f3c3be71 | 38b3f71f-fbf5-46cd-91b9-a6233ac79a89 | 6,737 |
| BRCA | TCGA-A2-A04Q | 77a8d1fa-9421-4333-b128-8a4b8fddfc6f | 4f762d03-b511-4e74-a7af-6633100d33cd | 295 |
| BRCA | TCGA-A2-A04T | 6dd42df3-896f-418a-9f13-468ac66d69bc | f9051235-b44d-460e-91f0-71118754ad79 | 5,620 |
| BRCA | TCGA-A2-A04X | 055e605f-56b9-4ca6-8856-1f32ab8e34a6 | c942bc80-18ce-4573-869b-4fc96677f775 | 1,503 |
| BRCA | TCGA-A2-A0CM | a6c27294-bf51-4eac-a2b6-885f5757b32b | 5aa2407a-3abf-42c8-817f-5de4731a1bc7 | 3,148 |
| BRCA | TCGA-A2-A0D0 | 25ddf041-bc97-4ced-aa1a-cf0339d8c23b | d5cf3968-3a25-4d98-9af5-206f96cce6db | 4,556 |
| BRCA | TCGA-A2-A0D1 | 6d37aa75-54cd-423e-987a-342e9a2def7a | a74be239-af6b-4564-9546-58302824434c | 2,899 |
| BRCA | TCGA-A2-A0D2 | 221d4b81-a70e-4647-8a9a-ce4866538392 | 956f0d4d-6f36-47f6-b9b0-f418ba4544c5 | 3,870 |
| BRCA | TCGA-A2-A0D4 | ac8422e5-dfeb-40c6-ab1f-dddd9f7fb015 | 231cf234-5a6f-486a-805f-8e2f12139cb6 | 1,603 |
| BRCA | TCGA-A2-A0EU | f8f8a691-8930-46b3-9cd7-1b8ac6895baf | bbc2932c-4507-4111-a367-ba914dfe257b | 205 |
| BRCA | TCGA-A2-A0EY | 79cc669f-41ce-4384-ac00-fe94ad0c8cd5 | 20d69bac-1191-4446-9c05-469bef595edd | 7,599 |
| BRCA | TCGA-A2-A0YG | 64d3bdd6-809b-4c4e-bb57-71f60383b777 | 0fe624ef-b6b8-45dc-8e70-c51c6a2e31d7 | 1,200 |
| BRCA | TCGA-A2-A259 | 767c5435-e3e1-4cdb-92e0-2b0961610118 | d2a43dcc-9fe7-4ca8-9f36-dcffd1ac50b3 | 472 |
| BRCA | TCGA-A2-A25B | 372d2951-a381-41f0-88d7-72cc9da6b6db | 639ddd7e-f28c-4b27-ae48-82d236b2f872 | 2,560 |
| BRCA | TCGA-A2-A3KC | 449d2c68-674e-4e86-89ff-574170845aa1 | 0dee69dc-a2f3-404c-b1d3-0155e3df4b39 | 715 |
| BRCA | TCGA-A2-A3Y0 | f1452ca5-f44b-4f35-a5eb-a8b1f9c44ef7 | 2a32c16c-432c-401f-9725-63857230380b | 5,593 |
| BRCA | TCGA-A7-A0CE | d744c7c6-690c-4f5e-968e-532936b3696a | 2ff7263f-3be1-441e-a60d-2fb82d7e3478 | 4,546 |
| BRCA | TCGA-A7-A0D9 | 43dc32db-2202-486d-82f4-388e37224379 | 4d84158c-282c-48bc-ae3d-55ada9b30f29 | 193 |
| BRCA | TCGA-A7-A13D | 3c8a206a-59d4-412a-97c1-9dd31cb9b7c0 | b80530e4-9ddc-40fa-a763-ea5010e21813 | 4,912 |
| BRCA | TCGA-A7-A26G | f7140aea-8ed7-4d5d-adfb-bd360e319805 | b1afd1ab-e46c-4616-8f8e-70509f3f1c32 | 757 |
| BRCA | TCGA-A7-A26J | ba14aedd-0b9a-4d6c-b5ed-80e7c732f0fa | d005c059-0532-4608-80e5-94bfb3de1e5a | 1,949 |
| BRCA | TCGA-A8-A07I | 6453f2d2-c0d0-46ee-a131-4a59788bb4c3 | 21138e96-7fba-4eca-b82b-98786a6858c3 | 2,879 |
| BRCA | TCGA-A8-A08B | f0967810-7b67-40b6-9995-a392153f2fba | ae70c8e9-45c8-4382-966a-e8787009508e | 1,715 |
| BRCA | TCGA-A8-A08L | 941e2188-e858-40d6-9f12-5faa6425a67f | aa5093d9-f4ec-4b1c-8e9b-eabedd7ee5e8 | 15,925 |
| BRCA | TCGA-A8-A08S | b026f426-bb3a-4c0e-b90d-6ec93bde0f53 | 88281394-1a54-4fd8-8b8c-7c6fd3134331 | 2,385 |
| BRCA | TCGA-A8-A092 | 3a2d527a-e52b-426a-aad8-0a7691cf835b | 9b123116-1002-41ca-a335-c49ced491c8b | 3,783 |
| BRCA | TCGA-A8-A094 | d706598d-75e8-4df1-94b7-da37b3361303 | 2cd07b47-50c8-4208-9637-21b2ec86670d | 8,516 |
| BRCA | TCGA-A8-A09I | 767c9f80-f27b-4abe-91c6-394a72835d7a | 0fdbc528-9b3a-4f41-b5e4-9e0f68b6823e | 2,949 |
| BRCA | TCGA-A8-A09X | 8327f8db-6932-4661-a116-5909cd8e4c81 | 9f917b23-c2a4-434b-87fa-4ea1453473db | 983 |
| BRCA | TCGA-AC-A2BK | 53c5c882-8386-48e7-b7dd-8815bcf6b316 | 0cdbbe0e-0c6f-4320-a72f-e80cf5faeb8d | 5,985 |
| BRCA | TCGA-AN-A04D | 8c35cbda-44bd-4a5f-9c1c-1a36d4636e93 | 4a1b29e7-a50b-4f6a-a8a5-0e1727c4647e | 6,502 |
| BRCA | TCGA-AN-A0AT | 5285cd99-09bc-48e5-abcb-11230d5f40f0 | 37554d25-cb93-436e-9379-74d3111e16c7 | 5,453 |
| BRCA | TCGA-AN-A0G0 | 293a4f6c-4b2b-4391-b076-ebebaa31e56d | 31f37360-2e47-4710-b5b5-57fc2b3a88e1 | 2,445 |
| BRCA | TCGA-AN-A0XR | 19e1e7d4-80dd-4107-98b2-6796fe22947f | 5820a5ab-6b80-4a87-8140-d96b70cd48d4 | 2,611 |
| BRCA | TCGA-AO-A03L | e374014a-934d-4eee-b1e3-53ec285ea08e | 03cb8dc7-1f67-4e48-a45f-f95f827694cd | 880 |
| BRCA | TCGA-AO-A03N | a198d001-d09b-4a43-9704-060db667151f | 6299b302-ec55-4baf-a73d-0da84bb6b964 | 2,503 |
| BRCA | TCGA-AO-A0J2 | 46cbf954-a4fb-4060-818c-a60666ae41d3 | 67104b13-f236-4501-aa7e-6a7eab9ec28e | 5,257 |
| BRCA | TCGA-AO-A0J4 | b518df9f-2eae-4c82-92e3-3eb196cdd3a5 | 4d68d5de-7417-49a8-adf6-12b3ccc612c0 | 3,456 |
| BRCA | TCGA-AO-A0J6 | 71b1d18c-e0ab-473a-b4ee-55ebb68cfda8 | 6de10403-e69b-4372-a6f5-e74b01a70129 | 7,132 |
| BRCA | TCGA-AO-A0JF | 031ab51d-07d0-4afe-a9dd-222889f5c32b | ecbc57df-ef58-47a5-9ae5-83e394d14191 | 90 |
| BRCA | TCGA-AO-A0JJ | 6d955c96-b34d-4aea-961b-378e856ae8cb | 335753c7-066c-41ad-8616-f9ec5f6b22e8 | 156 |
| BRCA | TCGA-AO-A0JL | b39a31b9-dd85-403d-ac11-d7b6dd2703a6 | 4db7f9bb-a3b8-4b87-b98c-8ba5be69963f | 295 |
| BRCA | TCGA-AO-A0JM | 456682bf-18d1-4383-96ba-6e06fe7dd669 | ef8744a1-5a99-4907-a1ad-fd44bb54635d | 1,396 |
| BRCA | TCGA-AO-A124 | 6ca7df84-fd75-4ecb-852b-76236b5781d9 | df77d00f-d0ce-435b-8776-c9043d199f1c | 10,328 |
| BRCA | TCGA-AO-A12H | 55b85b3a-3eed-4a89-ab63-56b85e180571 | 3acf783b-164a-4cee-bc63-deedb1ebc450 | 1,460 |
| BRCA | TCGA-AQ-A04J | c57fe789-7ef3-4dcb-bc21-b7ccb5ff208e | daa58157-819b-4aa9-b5f7-a7db1fb9acba | 1,657 |
| BRCA | TCGA-AR-A0TU | fb34c291-904a-4613-80cd-9788ff446157 | 79a26532-d93b-48b3-8756-9508d69f56ad | 88 |
| BRCA | TCGA-AR-A0TX | e9fddd42-1948-4df8-ac61-4db49de8123f | b17edad3-432f-495b-8595-03a17d1d0544 | 1,039 |
| BRCA | TCGA-AR-A1AY | 45a65b22-252b-4f92-bd7b-16a76d26c165 | 08e07f8e-1ab8-4039-a26d-4a7e1fcb9d98 | 3,308 |
| BRCA | TCGA-AR-A24Z | 1251c29c-8d85-4187-80dd-c459542e3080 | 832184fd-56a4-488a-a361-8fee484a5391 | 2,009 |
| BRCA | TCGA-AR-A256 | 83a38860-cf50-4344-8dbf-97d4cda96864 | 3c6b93f3-c90a-4e23-9fdb-f7d1d3e89a82 | 8,670 |
| BRCA | TCGA-AR-A2LK | 452eb1a3-2221-409b-a97e-f3266c5737cd | 2c451421-5276-4960-9791-c9d9c5a2b7ff | 1,909 |
| BRCA | TCGA-B6-A0I1 | 30cac8b3-f09f-4eeb-8dff-e77e8b9c13fd | ad8ec5b0-99a1-493d-bd8d-9bb358f87991 | 3,938 |
| BRCA | TCGA-B6-A0I2 | 1f31bdc1-69ef-49bf-8200-d73661b23c67 | 32d9eca6-d09e-458e-8564-2f50ca1eb655 | 2,330 |
| BRCA | TCGA-B6-A0I6 | aea89793-bb38-432a-ab2c-b5b9e7e93b1c | 92e425cc-f200-440b-a6ad-71e6feae486b | 6,853 |
| BRCA | TCGA-B6-A0IJ | b665cc0b-5952-419e-87a4-d97d3298f607 | b5a04a08-cf8c-4237-bb3b-d8fb7674c945 | 4,810 |
| BRCA | TCGA-B6-A0IQ | 7f45dc68-74e8-4953-8221-00c2ba51dee4 | 11ba6141-baa1-4403-beb6-4c5aa4dff23f | 1,699 |
| BRCA | TCGA-B6-A0RE | f7dd44eb-562f-4d27-8c45-5ec7fbf49448 | 67bd449b-7035-4a63-8e7e-e32f1f49e4f0 | 190 |
| BRCA | TCGA-B6-A0RG | b38ff19d-a769-4388-81e2-03263bba7985 | 5ad51bb7-5bcd-42bc-b3ad-64bb242f91b3 | 282 |
| BRCA | TCGA-B6-A0RI | 9283f68e-d261-4afd-b0eb-0a29fdda83ca | a498f419-7468-4a58-8a82-6692d63a46df | 174 |
| BRCA | TCGA-B6-A0RT | 0058348e-638c-426a-9fba-b26a3143c45b | 75db9c75-640f-4ad6-9d97-a0cb485b35d8 | 172 |
| BRCA | TCGA-B6-A0RU | 2efc5248-9be4-4e98-a400-f2ee744ac29d | a29cdc57-52ec-48f1-87d6-5f2a56480a48 | 1,995 |
| BRCA | TCGA-B6-A0WX | 1d0beca4-681e-4dbe-a6cf-f6c492c370b2 | 5df0fe6c-10bd-4c09-8766-283df19e6760 | 1,237 |
| BRCA | TCGA-B6-A0X4 | 1973be6c-c5a9-40d1-bdd2-a752dd33d6d1 | b1fb31e5-69fe-4970-b9ab-98311778aa34 | 78 |
| BRCA | TCGA-B6-A0X5 | b59ca3be-27cb-4b66-90cd-b0ee1c15b607 | 1859efd4-29c0-48f0-9b0a-533ab337913a | 2,828 |
| BRCA | TCGA-BH-A0AV | b2a1b1a3-a780-4be9-9a65-8937ba5df3b0 | d56f0f11-ae97-4969-a5f2-84945ca2c281 | 2,767 |
| BRCA | TCGA-BH-A0B3 | 82535082-739b-4a9b-8609-4724bbdd27ee | 29af0d77-d43d-4e85-9010-c496c956b250 | 1,442 |
| BRCA | TCGA-BH-A0B9 | 1a49cb56-547c-4a77-84e6-116bc406ba19 | 9822e8f0-c9dd-4681-a427-46ce76876761 | 1,775 |
| BRCA | TCGA-BH-A0BM | 46f039c4-a705-4933-bf85-0e96e80e5024 | dfd425ac-1f68-4ee1-b9cb-983f4bc32371 | 159 |
| BRCA | TCGA-BH-A0BW | d87bcaa2-19f0-4c3b-8f85-8f47ed056ffc | 190c051d-e6c9-4898-ae78-5946797b28ff | 4,316 |
| BRCA | TCGA-BH-A0DG | d1c980ac-6070-4a3b-8ff8-208b0f7897cf | f5b1fe9e-b024-4845-a47a-42964a009d73 | 579 |
| BRCA | TCGA-BH-A0DK | 71acb20b-f186-44d4-a92e-a2751f845fe3 | 70caaa1c-198e-4a38-9d77-d97bb89f7c65 | 124 |
| BRCA | TCGA-BH-A0DT | 4ac2ec89-0e27-40ab-949e-6cb716b4d9e1 | fafbdaae-4f36-4715-85b5-8cc9be96343c | 436 |
| BRCA | TCGA-BH-A0E0 | fc737335-21e7-4997-b267-04d3f91043f5 | ba938f20-4549-47f9-929c-3351cd03e1ce | 1,492 |
| BRCA | TCGA-BH-A0EA | b1ff5cfd-0f6e-4211-a58a-81b120d0eb6d | 9a8c10ef-1524-476d-aede-92a75cf6f28b | 448 |
| BRCA | TCGA-BH-A0GY | bcac7e1b-ddeb-43d6-ba44-db40336bf564 | 2e13622c-8cd4-44aa-a010-0de1e4dbcbee | 180 |
| BRCA | TCGA-BH-A0H0 | 11f5f965-f934-4815-a294-395459390c2b | 45b4fccd-bfa3-497e-a46a-b39a2b2d1263 | 1,879 |
| BRCA | TCGA-BH-A0H6 | 9c69c17b-33de-4bb8-be9f-7e0dcde9db0a | f8cb702c-5eff-49df-98b0-e1404066dc8f | 1,541 |
| BRCA | TCGA-BH-A0H7 | d5ab471a-88d1-4966-865a-f7fa29214fe4 | a267b823-747c-4258-b5e8-9cb33834ff6a | 183 |
| BRCA | TCGA-BH-A0HB | c9a434dc-8ce3-485f-88aa-a2373f6cdfac | 6c213158-4cff-4e3f-a2fe-79b1eb8349dc | 219 |
| BRCA | TCGA-BH-A0HK | c4f3a9e0-4b45-4e06-9320-171c27d9e28d | c9ee059e-f157-47b4-a76b-64e57c14f01d | 223 |
| BRCA | TCGA-BH-A0HX | 06f7db86-540c-401d-bbd9-0b88698c5f36 | f43d0ad4-4ad4-4388-833e-94faf1db1598 | 127 |
| BRCA | TCGA-BH-A0W5 | eeccc516-2d3d-4f41-a324-c90f92a6f944 | b768a52b-e629-46df-b042-fa815cb7ce57 | 92 |
| BRCA | TCGA-BH-A0WA | 1671f5e8-9bbd-41c1-803e-c518d5ba004f | 8ed08a9b-096c-47fc-85fe-61038535e131 | 4,178 |
| BRCA | TCGA-BH-A18R | 14062e7f-b11d-44a9-ae98-3383c517bf91 | 6d0781dd-0e6b-44b9-bdba-469778ae2a04 | 979 |
| BRCA | TCGA-BH-A18U | 8797ff7d-3b2a-4ac8-b8a3-375982e8413e | 66d1645c-a6d2-4a4f-9150-78bd1f576ae2 | 1,732 |
| BRCA | TCGA-BH-A1FC | 96f6b8e2-d4c6-44db-a8d4-3c7549e04038 | 689d4e64-e8da-4e5d-8d57-fce960e4285a | 2,746 |
| BRCA | TCGA-C8-A12L | c12fbad8-2c66-4967-9bfe-6837f6d1e5d4 | 1961adaf-b22b-4091-b508-99f711a4eaf1 | 3,947 |
| BRCA | TCGA-C8-A12Q | bb80680a-245c-421b-a3ef-198dc78bad1e | 53c73214-7b73-4c24-9cae-1c11fcb39573 | 3,195 |
| BRCA | TCGA-C8-A130 | e70e14f9-ea63-4038-8a0e-5526391ea4fe | 44978a5f-a9ce-40ac-8b2e-3ade35e32525 | 1,679 |
| BRCA | TCGA-D8-A27F | f4f2c07e-84b7-47d0-88fe-7032564a1ecc | 5e608249-2680-4ecf-ba66-89998d2d6a1d | 6,968 |
| BRCA | TCGA-D8-A27H | bdde252c-e43c-448b-8e22-1ff90c7c957b | 71bbc4fc-5b27-43d5-8475-9a1d1c744147 | 7,035 |
| BRCA | TCGA-E2-A109 | 33884ffd-f43a-4cb9-beb0-9c0a7d6f3f31 | bd77a131-48d3-4950-992a-ace47923f7ae | 5,001 |
| BRCA | TCGA-E2-A14P | 6291117d-64a7-4c35-94bb-8e06275c498e | a639fa8f-f8d6-4d5c-85bf-5a37c02258d4 | 2,550 |
| BRCA | TCGA-E2-A14X | 775348e1-91bf-4bf9-93e9-ff9f955b5a79 | 0a820852-4686-4bed-8995-35aba8b37368 | 779 |
| BRCA | TCGA-E2-A152 | fe904377-4375-437b-8774-c80f11c6e0ff | a5f27034-eded-4286-bf8f-965029909919 | 3,420 |
| BRCA | TCGA-E2-A156 | 3d9a2788-2bc3-4df9-bae7-fef37f6096b8 | d87cfc01-42b6-4dc0-9403-d2daee055209 | 1,631 |
| BRCA | TCGA-E2-A15E | 3b80e735-e034-4867-be5e-52500f523ab5 | 9ed4cd7a-c9d6-47d6-8258-def05e151530 | 984 |
| BRCA | TCGA-E2-A15H | f359078d-7930-4e46-a5d4-3978f5434622 | b2b05078-b11f-488e-afc8-2d70b062609e | 905 |
| BRCA | TCGA-E2-A15K | 6463e302-836b-4097-b6c5-acf33fe1a733 | 4968e751-3f12-4e99-8228-6a58cbf1bca4 | 2,746 |
| BRCA | TCGA-E2-A1LG | c24e6791-401c-4b37-bfe6-669f050ea42e | 6bd8c3ea-15b6-4d42-b4f1-bbfdacb145f8 | 4,100 |
| BRCA | TCGA-E2-A1LK | cdbdb1d9-4c6d-4a22-8867-42722b596ae9 | 32766841-2370-4aad-b93e-8d5e8b002229 | 2,774 |
| BRCA | TCGA-E2-A1LL | 85360168-d41d-4201-8416-2cb4b45fca0f | f2fe81a1-3e80-4b03-9699-eaa8a4652be5 | 2,769 |
| BRCA | TCGA-E9-A1NH | f57cf947-b314-4a64-a112-d09cc8045594 | 9ec85678-e1a8-44bd-9676-72b0d1b991d7 | 1,207 |
| BRCA | TCGA-EW-A1J5 | 17593e20-8cc4-4961-8ff5-871806aeed93 | ce79ca01-0c22-4e25-a5da-fd9438299f91 | 21,753 |
| BRCA | TCGA-EW-A1P8 | 3a2f0cf6-0405-4ef6-be7e-75054acf1343 | ab0a625b-6478-4d76-8b3e-b3e479e4242e | 2,109 |
| BRCA | TCGA-EW-A1PB | 2ba1984b-043e-4cc5-8bc4-10b6c2915f61 | 46b06dbe-5f5d-4733-8c98-f4a3fdf451c4 | 5,885 |
| BRCA | TCGA-EW-A1PC | 4ded0771-e685-45fb-99f1-4dc1212cc9fc | bbbfdc86-7d19-4c67-9a3f-c46da84f900d | 4,642 |
| BRCA | TCGA-EW-A1PH | 877bbc2d-444f-44cf-8e0f-3fd2744c3f91 | 9bb03dd1-4a31-4ff3-930f-2ccf230339ae | 2,646 |
| BRCA | TCGA-EW-A3U0 | 107557e5-5611-4f30-9544-e8c315cb063d | e49f86f3-22dd-40c7-8b86-73ec1d90694b | 2,935 |
| BRCA | TCGA-GI-A2C9 | 5f4a8fdf-00c0-4ab7-8323-d04f23a9e4a8 | 70c562ea-b1bc-41ac-aa70-e3865fb5f03e | 5,484 |
| BRCA | TCGA-GM-A3XL | 77d3443d-1bf9-4867-89a2-65cc8e10141b | 1c380c75-c405-42a9-8984-c87a63cc94a2 | 4,213 |
| BLCA | TCGA-BL-A0C8 | 19df03d2-75d2-408e-86e2-482ddeed9736 | ccc5396b-d9e0-4f96-85ec-4a894585996f | 467 |
| BLCA | TCGA-BL-A13I | b7cfe609-703a-4750-b9e7-5bda8e948cc4 | 18c5e150-2417-4ca7-88c8-d147dfb43853 | 1,240 |
| BLCA | TCGA-BL-A13J | 02da398d-72c4-4356-b121-920eea9c4013 | 3c2c919b-b44a-461c-9ebc-614bce0d53cc | 2,835 |
| BLCA | TCGA-BL-A3JM | 5cd881b7-caa0-43ad-9ba8-f9696fdf7403 | 33f45265-9a07-4291-ad56-6a47f9838213 | 193 |
| BLCA | TCGA-BT-A0S7 | 55a63ec5-a803-44ee-905f-c7bfff9162d6 | 2414eb9d-53e3-403c-87e5-dcb81ae001db | 1,790 |
| BLCA | TCGA-BT-A0YX | 71886c7b-a73b-4c00-9555-5fd61b5e1347 | a3d8075a-0e5e-4177-ba61-0f8cf6367170 | 1,708 |
| BLCA | TCGA-BT-A20N | c7de2a6f-d24f-4be6-98af-307070063116 | 3a98735f-f5b6-4273-ac15-6d6f17971bcf | 2,143 |
| BLCA | TCGA-BT-A20O | c345a912-f3c8-4622-9793-88cf795fc964 | d9764a1d-1e8f-47de-8916-e254751cd477 | 676 |
| BLCA | TCGA-BT-A20P | d714be1b-c30c-4f13-8a00-4bf93507bb1c | 84bd8301-5869-4245-a3cc-3781508f00c5 | 4,038 |
| BLCA | TCGA-BT-A20Q | 9d2dc2d3-8995-42b8-9bee-010c8bce06e0 | c4e57539-5d0e-43ce-87df-73ae5c8e8738 | 5,773 |
| BLCA | TCGA-BT-A20R | e997f623-927f-4dae-8fe9-590eef9a9f59 | 81815e13-7a11-4252-9fea-8006d97d9a9b | 535 |
| BLCA | TCGA-BT-A20T | 7d40d139-a746-424d-9f90-d5d3da5e98cb | 1f33179c-e661-4003-9bb0-d0f222368c02 | 20,177 |
| BLCA | TCGA-BT-A20U | 9947c8cb-fa3d-4cc2-942e-b013f7a47faa | 62a2b30e-c3e7-4831-8cda-b72f3d30157a | 2,409 |
| BLCA | TCGA-BT-A20V | a193bc83-ec94-46bb-b7ee-60b8b75c8ad8 | cdb0bfa4-f9fb-4cd7-b19a-84f9afc80f69 | 548 |
| BLCA | TCGA-BT-A20W | a817f2b5-a06a-422e-869a-694baf6bb127 | f3f37b3b-3755-4102-b654-abea4d035ed1 | 841 |
| BLCA | TCGA-BT-A20X | 256c8b67-1fea-4666-b42c-21e8860946e3 | 85b68bda-849a-4597-8161-176a8ad06f09 | 599 |
| BLCA | TCGA-BT-A2LA | 587f05f3-8e6a-4a97-b59b-29371f2f8b74 | ee7f989e-a375-4956-a120-aab1394e350f | 134 |
| BLCA | TCGA-BT-A2LB | de67212f-594e-409c-bd19-3d40f013b89a | 832b34b6-3018-44f5-b83b-4b2b9f5f032f | 917 |
| BLCA | TCGA-BT-A2LD | 205bfe9f-2bc9-4236-930c-a862874ad8e0 | 81838e98-98aa-4bf3-94e0-1819bd2a4c6b | 94 |
| BLCA | TCGA-BT-A3PH | 9096dde9-2391-4c22-b816-2057b8adb483 | b3c44d04-1766-41e1-9f24-464ed682c9a3 | 18,156 |
| BLCA | TCGA-BT-A3PJ | d7aa3a92-c61a-478e-af73-14d0204cba5d | 717b2eb2-3d5e-4477-95da-1f30739ece0b | 3,070 |
| BLCA | TCGA-BT-A3PK | 637e8bd5-61ee-4d45-8cc5-e3b89950d020 | 17390bf6-09d7-4acf-bf79-b8cc3eded7e1 | 360 |
| BLCA | TCGA-C4-A0F0 | 38f56daa-7d69-4fe0-8895-10e22397ee2a | c5e7b6e1-16a2-46ee-85ed-49e5092c1e70 | 546 |
| BLCA | TCGA-C4-A0F6 | 2332be3a-c2a5-446a-ae60-b8f873fe1866 | b54ac4b7-3c94-4496-bf96-61b3ca3db0c5 | 2,860 |
| BLCA | TCGA-C4-A0F7 | f2782745-2bc7-47f9-8148-4ea61ecb7565 | f9c05a41-994c-402c-b582-8efe6bdf32bd | 7,199 |
| BLCA | TCGA-CF-A1HR | 65dee943-f41b-4f59-a5bb-2477053d1079 | 8f5d258a-5720-4244-a4e2-2288d64a7ef9 | 169 |
| BLCA | TCGA-CF-A1HS | ed90abf4-1d04-41ce-9de3-9dcf24c1e375 | 6774ba08-2ee0-4555-946b-3fcf2996801b | 254 |
| BLCA | TCGA-CF-A27C | 456bc333-4951-4b6e-845e-2af77b0a86d7 | ca3f55a3-b068-4298-8352-e30c4c9affad | 7,173 |
| BLCA | TCGA-CF-A3MF | 62e960fb-aaf6-4d7e-b15d-1b5136f6fa37 | 10a650d8-0b73-456b-a674-14fe44e158ec | 217 |
| BLCA | TCGA-CF-A3MG | 695ed631-423c-4381-ae8f-4a70fd6266b9 | 1385e047-6123-41fc-bd33-96122ff0e918 | 149 |
| BLCA | TCGA-CF-A3MH | f82c1606-fe81-428d-9889-58cafa1f29e4 | dd0cfd9f-503f-4f83-9fd9-acd2b40f14f0 | 223 |
| BLCA | TCGA-CF-A3MI | 6559c325-d368-4f95-a2c0-df5efa895e3d | b157e6ff-21c1-412a-953b-66770d827a6a | 1,182 |
| BLCA | TCGA-CU-A0YN | 89fcc222-6a37-406f-a641-b3d857104f9f | 4f24b2d4-a63d-40e7-82c7-bc6e188fbb4a | 280 |
| BLCA | TCGA-CU-A0YO | 98d555c2-93ae-4db5-9b19-43655d10609d | d9b681f1-ebb5-4f97-8585-796e6ae8fb6d | 295 |
| BLCA | TCGA-CU-A0YR | 197d19a7-07c1-4cd3-b41d-a8233c9d8a42 | 1efc9ebb-a445-4460-9053-367c5b4daaf5 | 209 |
| BLCA | TCGA-CU-A3KJ | 6b7a9ee7-ee89-4473-9170-7372313595f7 | a2af4db6-b93c-4e1f-9be7-28fdb75431a6 | 167 |
| BLCA | TCGA-CU-A3QU | c4c9ea17-be51-4e92-8cdc-0d5880def11c | 2cb07ca9-b658-4d19-ac30-c9da9c4550d8 | 311 |
| BLCA | TCGA-CU-A3YL | 8753d7f5-4d92-4781-8d19-d44710fa4f45 | 69971830-c186-43e1-a945-c5eb5bd2b1e3 | 237 |
| BLCA | TCGA-DK-A1A3 | 0e78dec3-7b8b-425e-9cc5-489bf825e0ea | dea57f3e-8079-4c79-9961-51b68eba2dcc | 1,749 |
| BLCA | TCGA-DK-A1A5 | 3aa26bad-446a-4391-a584-27b78fc638e8 | 43d7b3d5-6a0c-49f1-a63c-d49c8fdb8c86 | 645 |
| BLCA | TCGA-DK-A1A6 | 694693d8-dc04-4516-bc18-e8a9ccb24d28 | 182b13b1-b615-4fdf-8972-face22902f2c | 20,403 |
| BLCA | TCGA-DK-A1A7 | 4eade8e4-0984-473e-99f5-523fdc631190 | 77c6d0de-f590-405b-8d12-084a7ebbace2 | 6,598 |
| BLCA | TCGA-DK-A1AA | 006c1498-f327-4fff-8936-7d7cb06639f4 | c6b09f3f-3fe9-4d73-a1d9-c6ce0084ac1e | 2,037 |
| BLCA | TCGA-DK-A1AB | a5005e15-2ada-4bf4-9a61-0febc81a99ca | 9eaeaeec-3d40-4c27-8e58-b9aa313a628f | 839 |
| BLCA | TCGA-DK-A1AC | befc5d25-55df-4a22-b4e4-25508f3935c9 | 7b5dbb5a-bca2-4772-8f0f-af7cdca1713e | 3,317 |
| BLCA | TCGA-DK-A1AE | 2722f260-d3b4-4088-860d-7a5070921255 | bd5c0062-f4b4-40fd-abd7-657e17170eca | 1,677 |
| BLCA | TCGA-DK-A1AG | af228568-1064-4709-8bc8-435278c73ca0 | 61d6a720-87af-44e2-9d20-b63f90902707 | 7,544 |
| BLCA | TCGA-DK-A2HX | 17e4edb9-ac80-43de-a7c1-66edf1201ac0 | eb041cfc-02dc-491c-a584-db245f5d273a | 871 |
| BLCA | TCGA-DK-A2I1 | 724308ea-3fa9-4762-9800-445a044e94f2 | 7f2d5952-2a08-48c4-b4c9-5773f777866e | 249 |
| BLCA | TCGA-DK-A2I2 | 99517da2-f39e-4005-8efc-4487b92c0dcc | e257ae8b-2d82-4b7d-b7a2-976b611b16b2 | 319 |
| BLCA | TCGA-DK-A2I6 | 6567cd2d-a1e5-4226-9239-77c48c09c55e | 4456fd98-e15f-406d-a4be-d8e3ec851888 | 2,218 |
| BLCA | TCGA-DK-A3IK | 6c286798-955c-487f-b541-72f253866ada | 7044d689-15dd-4f2a-ab94-f0e69c9f157a | 214 |
| BLCA | TCGA-DK-A3IL | 092f2192-b2d3-48e8-be2f-f56c1febe40b | bacc62d8-d2ef-4268-8478-db3af3db5479 | 1,585 |
| BLCA | TCGA-DK-A3IM | 27ca0d22-3ef8-4c3c-814a-94dfb76c3289 | 1f1106b7-eff0-48a4-828a-0a42816b6736 | 510 |
| BLCA | TCGA-DK-A3IN | ff3d1235-2aed-4091-a8a2-72340b403e05 | 7f0ce9af-6471-4dd5-9040-7facfecf821b | 2,055 |
| BLCA | TCGA-DK-A3IQ | edc43d06-7118-46e7-894e-d40138b6d69c | 0cfcdefd-603d-4cc2-a651-88d3e5f5c56c | 410 |
| BLCA | TCGA-DK-A3IT | 276fd52d-8d51-4e03-85a5-a7f02ee03b71 | a0fae593-7715-4c62-865f-1a19df61e8c1 | 1,823 |
| BLCA | TCGA-DK-A3IU | 346b87a3-0c3a-455c-b3c1-0fb96acd0f2f | ef01cea3-48ce-4afb-a6e9-4a52d7c8a07b | 179 |
| BLCA | TCGA-DK-A3IV | 5bf5cb9d-7592-4967-8440-fe80dfe9c2a6 | 0e2f8cf4-b42d-4b2c-a2f1-68e364dd633d | 109 |
| BLCA | TCGA-DK-A3WX | 12365344-0a7a-4ef2-a651-2f1f3a15f6a1 | bed65641-3cb7-440c-a595-ed3f210f52a2 | 185 |
| BLCA | TCGA-DK-A3WY | ae1b3911-b342-4886-b75a-3e9be21dae41 | 0d07635d-ca76-4aca-986a-7167437f8520 | 214 |
| BLCA | TCGA-DK-A3X1 | 4c8de054-fd6d-4ebd-8a6e-b5061ef5913b | a711317a-7fc9-4637-968e-842d8b6afc22 | 404 |
| BLCA | TCGA-DK-A3X2 | a8b7a158-32d0-421e-82bd-edb6a44d69e0 | dc4b6741-a258-4b67-9da6-c236d90bddf7 | 326 |
| BLCA | TCGA-E5-A2PC | 496052c5-5ab2-4852-88e1-348116a86746 | d15f0d2c-94bd-40f7-8589-0087c2ec6339 | 2,831 |
| BLCA | TCGA-E7-A3X6 | 548e206a-eaec-49d3-a405-664cf76d0e16 | 2068d801-01a5-4304-a51d-95555d328e12 | 410 |
| BLCA | TCGA-E7-A3Y1 | aaeb4857-84bb-4a72-9e0d-81280de1ca61 | 57523af1-f9b2-4f88-abec-4cf325040361 | 272 |
| BLCA | TCGA-FD-A3B3 | 6b702a7f-ff83-4ff2-96a8-9da2f015ebcc | 675113cf-e818-448a-a767-827cd45c7827 | 447 |
| BLCA | TCGA-FD-A3B4 | 54b70ff2-6c53-438c-934a-97fe2837620e | c312905a-2708-40b3-9055-264aa217c722 | 573 |
| BLCA | TCGA-FD-A3B5 | d7af6a65-2c95-4a83-8e6f-756aef5309bd | 47bf4c5b-dc12-4d32-bdb4-e95b09f41768 | 1,414 |
| BLCA | TCGA-FD-A3B6 | 15aeae5e-1ef3-4574-a7af-025ac4d4be89 | e74f45ed-30c8-4516-9011-9146d0451714 | 1,299 |
| BLCA | TCGA-FD-A3B7 | 4a37fdf6-816c-46ca-b703-cf3d58b32149 | 846e6076-8a19-4673-95e6-f23e24f87b29 | 758 |
| BLCA | TCGA-FD-A3B8 | c0aadcb9-2d54-45c7-9585-575ed21ea52f | cc9d7008-bf77-41cc-b38b-bf5af9413291 | 638 |
| BLCA | TCGA-FD-A3N5 | 3519f8d9-0eab-45e8-b398-5ddc469243cd | 2ca8a15e-2f08-4dfa-9203-7f5d50080161 | 4,003 |
| BLCA | TCGA-FD-A3N6 | 2beb0463-75e7-4244-b88e-022e447445f2 | 65a2ebc7-7a90-4db7-9eb2-7123085b6ba0 | 232 |
| BLCA | TCGA-FD-A3NA | 8c4c1672-fea7-4c17-a6d0-d4a0f3c04558 | 6e999ca0-d8a6-4bc2-9630-4637a8193a3b | 557 |
| BLCA | TCGA-FD-A3SJ | 2635ae0a-878a-4237-ad5c-5f0092134ee3 | 42daaedd-31f4-4764-92e7-8eeccec3ee06 | 469 |
| BLCA | TCGA-FD-A3SL | 764ae567-a7b6-4d13-bef1-8cc55afb4524 | ed617be2-c160-4372-8c95-3b4037dfc801 | 360 |
| BLCA | TCGA-FD-A3SM | bacf3136-6605-4a66-a869-8d91e4003a0d | 63dc4922-4d08-42bf-bb9a-10a25d19c59b | 362 |
| BLCA | TCGA-FD-A3SN | 2281a267-dad6-42d6-9df2-8d37a696c507 | 6b3838e6-41ba-4745-bd39-cc0d6771fd98 | 968 |
| BLCA | TCGA-FD-A3SO | 6b296020-484a-4cc2-a114-6c8a34ed5712 | db8ba963-75f5-4fb2-a6bd-3da7bc62ab16 | 2,227 |
| BLCA | TCGA-FD-A3SP | 2da38f98-8b94-404e-99e7-7a8d58aa25c1 | 0fb6c4ad-36eb-4250-aff4-193b6fe1badc | 722 |
| BLCA | TCGA-FD-A3SQ | 104a1ca4-a3b6-42e6-b6ba-1b0916160573 | ccb5242d-6bc3-405b-829a-ed7ff3c509b0 | 1,092 |
| BLCA | TCGA-FD-A3SR | e8c95d25-27f4-4cba-b644-eb39019c6cc0 | ea09ef7b-e64d-4515-bd29-48e6bb5e9522 | 835 |
| BLCA | TCGA-FD-A3SS | 23aab0ed-a2ae-43db-9050-527db1729aa5 | 275f5d73-157d-4162-a8da-dec4f5f8c299 | 656 |
| BLCA | TCGA-FT-A3EE | cff49411-c565-469a-83c7-9a253f6b129d | 2398ce28-2b46-494f-a9fa-1bc987618a00 | 7,643 |
| BLCA | TCGA-G2-A2EC | 0d4b7559-34dc-4690-a8e9-144698537cdd | b2ae08c3-020a-401c-9a25-c937c153c403 | 1,222 |
| BLCA | TCGA-G2-A2EF | ac16538b-bd5d-496e-bccc-6f3374d226c6 | a641053d-d0ab-4204-a4c7-6fff5ed2ef4e | 84 |
| BLCA | TCGA-G2-A2EJ | 18abb4d7-25d0-477e-bda6-011263e96364 | 5ca74123-b721-4ec8-bade-785975c82891 | 1,919 |
| BLCA | TCGA-G2-A2EK | f98c8702-90e5-4e7a-ad12-d11a9af57bf5 | 2d89e6fb-9416-4502-9329-ae8eaa2caa9c | 568 |
| BLCA | TCGA-G2-A2EL | 15128dae-4500-4372-9676-d23d4c116527 | 9b81589b-68e2-4101-b77c-a356f255eb95 | 79 |
| BLCA | TCGA-G2-A2ES | 5922785f-2336-403c-9cb6-0843df3383b5 | f8583339-ea56-43ba-af11-e7b5945ecfb4 | 2,164 |
| BLCA | TCGA-G2-A3IB | f78443a3-8c1b-429a-93c3-fcd93f3c6d55 | a8fee664-cc6f-4175-9b23-497bb28d2dd6 | 1,177 |
| BLCA | TCGA-G2-A3IE | 171a2a97-43d5-41cc-a9ae-2f685c3ed80f | 4630216d-d5a0-4073-a48f-8326641af6de | 446 |
| BLCA | TCGA-G2-A3VY | fc9767fe-ddfb-4540-b151-0a80300c0a1d | 5135b458-af9a-472c-a3b2-0456e678130e | 1,955 |
| BLCA | TCGA-GC-A3BM | 1901f2a9-837a-482c-8073-472f3c4d9ba1 | 1701d500-8ca9-4b1b-87a0-fccb8808328e | 282 |
| BLCA | TCGA-GC-A3I6 | ec0a1f1d-c6e7-4b4a-a296-02087e0b6586 | 792b4366-c23e-4272-8b38-4ac0acc9ddf8 | 1,471 |
| BLCA | TCGA-GC-A3OO | 8c665767-edb1-4860-b26a-4171237962fa | 288ab04f-61f7-4278-8ac5-e7a12d87a6ae | 135 |
| BLCA | TCGA-GC-A3RD | 27573bf6-b72d-47e6-93f7-38d2887d8b32 | b81dceea-a960-49fc-9cfc-00d61570a8b0 | 290 |
| BLCA | TCGA-GC-A3WC | 5d982317-5753-41de-85ef-30ee85c56ba9 | 33924502-cf1d-4609-9e24-f717bcd9c439 | 561 |
| BLCA | TCGA-GD-A2C5 | 27e88ee3-dcb4-43c4-9653-c55aa1851cb1 | d56808f7-e6d9-4234-b65c-7ff6bad653f8 | 5,612 |
| BLCA | TCGA-GD-A3OP | e01f55a8-4fbe-4f9a-8617-ce0e6900f177 | cec19b51-29ce-4b48-81a6-7f61054cc053 | 453 |
| BLCA | TCGA-GD-A3OQ | a07d0456-032d-409c-af8d-c4d6349ef4db | b206c126-8a3d-4051-89bb-086c3e66ed39 | 164 |
| BLCA | TCGA-GD-A3OS | 2a801113-8404-4c17-9d60-508761ec1659 | 70ba803f-783d-4116-9e3b-0798393f59ac | 811 |
| BLCA | TCGA-GV-A3JV | 8f626567-33e0-4b6a-974f-6b41b341c60e | a8f1ba67-bcef-45d1-b508-d4ce580d4362 | 307 |
| BLCA | TCGA-GV-A3JW | 586abfdd-f50a-4834-8143-3bda5638ea60 | c147acc0-a739-4bac-a729-755e35fb2783 | 526 |
| BLCA | TCGA-GV-A3JX | 317cc3f0-7a98-402b-bb1f-81a0437fbaf8 | 4e9101b0-24aa-443e-aabd-2a8d28d2f118 | 179 |
| BLCA | TCGA-GV-A3JZ | d4d5d34c-9665-4291-a3d6-14bc9b48f2fa | 4b54f30a-eae2-443c-8c0c-f198b72091d1 | 234 |
| BLCA | TCGA-GV-A3QF | bcac19c7-1e8c-40ba-a577-39fce12db063 | c19d7e07-cd68-4880-a727-0e2717a37461 | 1,260 |
| BLCA | TCGA-GV-A3QG | 2102b9a0-6afe-47a9-89ae-9778452a6dcd | ee01ec50-47b1-43fa-96a1-b63ae8f92127 | 240 |
| BLCA | TCGA-GV-A3QH | 5672aee8-638c-45c6-af4f-0ccc5e2a1b45 | f84af011-24a9-4fbb-b801-5d6062f0b896 | 342 |
| BLCA | TCGA-H4-A2HQ | 46adb3c9-ac82-4284-b0d1-f95b10435695 | feb80610-46c5-490f-a029-9c6b401aeb25 | 1,977 |
| BLCA | TCGA-HQ-A2OE | 21c45d2f-f8bc-45dc-85d2-790ea5506bc4 | 2ff1fc67-2977-45d1-baca-e4098f7bca8d | 1,288 |
| BLCA | TCGA-K4-A3WS | 891c215b-20ad-4eb3-9e53-87910b1d06f4 | c3da67a4-50f4-4eb4-999c-b861247e86ee | 637 |
| BLCA | TCGA-K4-A3WV | ca7a6b06-a6fa-4314-80ef-725b81781598 | 3dc4b7a1-75cf-4f52-b921-7d6dcb7d1046 | 66 |
| PRAD | TCGA-CH-5741 | ae88c859-e075-4e3a-ba86-e6018e71fa98 | b82ce0ca-281f-43c5-a79e-de66036ebf94 | 288 |
| PRAD | TCGA-CH-5743 | 1c58f516-3239-499a-bbcd-f5a8d619d796 | 9e05f936-10ec-46a8-96d8-efc5c8510e51 | 194 |
| PRAD | TCGA-CH-5744 | b48809ce-93ac-45d2-b127-bf6ee8bd6870 | 7e3d3253-ca6c-4dbf-97ea-4d9126f9acad | 176 |
| PRAD | TCGA-CH-5745 | c1d27dca-0355-41e5-a53b-0af94a125c6a | 9ea9f797-df7f-4211-af54-30bb1c25941b | 259 |
| PRAD | TCGA-CH-5746 | c4373feb-ceca-4bb3-9db2-6f9e94d29605 | f96d7355-1ea6-48c4-8399-14a9bef45847 | 280 |
| PRAD | TCGA-CH-5748 | 02aa8440-93b8-43d7-a58b-6c4421e05321 | c1f919a6-32b6-4134-a118-61ce369b31d2 | 316 |
| PRAD | TCGA-CH-5750 | d0338227-feec-4f2e-bcfa-89c723bb89b5 | 54cfd889-407a-47bf-97d6-e938fb0f0f0c | 366 |
| PRAD | TCGA-CH-5751 | c152f07f-eb7f-464d-ba51-ce1b55777952 | 31eb5a98-80e8-470f-9e31-765c00c7b655 | 280 |
| PRAD | TCGA-CH-5752 | d1bae4e7-eb79-4ede-abb5-19a328e4ad0c | 78b4b6a6-b8b1-4499-8d0d-6f183072b505 | 779 |
| PRAD | TCGA-CH-5753 | 971890ca-4817-4b12-b817-51caf2239fa3 | 7244700c-38c6-43d7-8edd-8ffe49b8b04c | 530 |
| PRAD | TCGA-CH-5754 | 9b3dfe47-a8c3-4a73-8aa3-66af9d5001a2 | eff5ccea-5f74-449e-a9e6-4eb5a83e71c7 | 966 |
| PRAD | TCGA-CH-5761 | 4d316299-7958-45e8-93ab-2150e05d3210 | ce8211c1-e866-447d-9c68-847d7d6d07f8 | 679 |
| PRAD | TCGA-CH-5762 | 70b297c6-c1f6-4651-93c3-17888cf0b118 | 823dc639-947b-44e0-839b-882ab9437fac | 383 |
| PRAD | TCGA-CH-5763 | ba68ecb8-6421-467d-b680-850c82a0b95d | 51855ca2-0a23-4f4a-8e63-5b34fcb33c98 | 282 |
| PRAD | TCGA-CH-5764 | 036937fb-20a8-45df-a234-f7396aefb2d1 | 2a2319bf-1c6c-4a8d-84e0-a9b256cb0903 | 374 |
| PRAD | TCGA-CH-5765 | 265ff01e-199b-4908-acaa-62aa8986967a | b2118b85-15e8-4468-ac1e-ae5862adf28d | 445 |
| PRAD | TCGA-CH-5766 | f778efd0-d2ed-48bc-ab57-e2bd12130169 | 3e827e6b-5861-47a6-9904-89cef9f94a0f | 381 |
| PRAD | TCGA-CH-5767 | 4ec6dbff-5194-4938-949c-d5ae0ec6a882 | 1cbd34c5-4fa8-46ee-b97f-ccbd7b816165 | 140 |
| PRAD | TCGA-CH-5768 | 98aa2eb7-52a4-424c-aa01-a574d0215ee6 | 94474228-f50e-43be-ae1f-20104c2b61cc | 430 |
| PRAD | TCGA-CH-5769 | 2fe51267-827f-4ffb-8aa8-42352e2352d5 | f2ed1ffa-45fe-4497-9137-3a7270cc913b | 361 |
| PRAD | TCGA-CH-5771 | e5382b76-d2ee-4eab-b245-b0d58fda770e | 8db96075-ac39-478a-b26f-a624d6a83712 | 226 |
| PRAD | TCGA-CH-5772 | 97cf547a-bc4b-4b08-9f8e-c1986b18aa30 | 4101a708-70fa-42dd-8aae-c5e3d3b12dd5 | 406 |
| PRAD | TCGA-CH-5788 | fc337685-dcbd-4e7f-9b5b-369121193a7a | 4087c08f-e268-43c8-939a-54a67cf78246 | 2,636 |
| PRAD | TCGA-CH-5789 | fc8da645-2fde-4ef8-9ee7-8b8101580793 | 834487d2-4198-43c2-b567-02b05a6d52f9 | 188 |
| PRAD | TCGA-CH-5790 | 22e486a4-e20b-4211-b3fd-35f37d14e718 | d44cf3be-e75c-45a3-a47b-d7571c521c1b | 281 |
| PRAD | TCGA-CH-5791 | e2778c71-6feb-40df-8b1e-8d8f0e889beb | 747350b7-a5fa-4e6a-bf4b-72ea53989d5c | 187 |
| PRAD | TCGA-CH-5792 | 1c4a46f6-fc5f-44b1-b30a-5baaa806e78b | a3ab0225-b534-4b3f-8356-d82c84c70db2 | 158 |
| PRAD | TCGA-CH-5794 | 2cf518ee-af91-4cf0-a277-263170f71791 | 3c58e5e5-3f65-4e72-8ec0-9975b3385833 | 240 |
| PRAD | TCGA-EJ-5494 | 01d186ab-6289-40da-9715-4f766da74b10 | 7138c14a-2dab-4b19-b460-da77aa8d9819 | 128 |
| PRAD | TCGA-EJ-5495 | a2c14576-771c-460f-a71b-e25651274ca1 | c1976dc5-9458-46ad-a1a5-1db90aa4eb04 | 305 |
| PRAD | TCGA-EJ-5496 | 00039222-b24d-4c86-b6ac-ffc00b3b6ece | e93a13bd-c687-4275-9c7e-75b9b3c024b2 | 164 |
| PRAD | TCGA-EJ-5497 | 453a9559-644e-46b9-8be3-7e4511448e7e | e9e5af5e-fd0b-4048-9b62-94b0e5e932b8 | 100 |
| PRAD | TCGA-EJ-5498 | 1ea5110e-6493-415f-ae21-019a8d1e2a17 | ed7ab548-eb8d-46b7-ae35-1b081b1ed787 | 275 |
| PRAD | TCGA-EJ-5499 | 21f9da4c-83ce-44b4-b002-e18cd232ac6b | 3635d2b0-fde1-42b9-8831-c429d8b8e864 | 60 |
| PRAD | TCGA-EJ-5501 | 6e2cf9a5-b8c5-4f44-8800-c95ad84bc9a8 | 91be46fe-3b96-4d1b-a7f4-229e972c32d1 | 250 |
| PRAD | TCGA-EJ-5502 | 40f605db-50a6-4ed1-b652-61045959ba2e | ba38e3c4-9b7a-47f3-819a-86b4b443cdd3 | 188 |
| PRAD | TCGA-EJ-5503 | 4eba210f-0895-487c-9b48-db8a8c629cc4 | 6f2a0bc7-14c0-431b-ad1a-9ed9cdc21dc4 | 159 |
| PRAD | TCGA-EJ-5504 | ea018935-4ded-4def-a204-bda7035ba32c | fdf14dea-7dab-403e-9c85-8b16dcf88d2c | 293 |
| PRAD | TCGA-EJ-5505 | 156ad547-dfb8-48c9-88b9-6842c81fead1 | b49d48cf-faee-4790-83ab-3920e77326b5 | 174 |
| PRAD | TCGA-EJ-5506 | bd66d4a0-aa1a-4bf3-a764-cb7292bc89b1 | 299c9fa9-0cc5-4e97-80d8-e0b7c60bd861 | 279 |
| PRAD | TCGA-EJ-5507 | 0dbb7308-269c-4ad8-97c7-e7475ae08189 | f6d132bc-c74a-4472-8e91-6d86e10f785e | 210 |
| PRAD | TCGA-EJ-5508 | 69098ac9-e4d2-44cd-ae04-e179d90a749e | 0d2993b7-024f-4363-9ff1-9d3ce308eca0 | 105 |
| PRAD | TCGA-EJ-5509 | 6fd44a89-a20c-4ecc-bafe-54da9b58cd9d | 47e80516-5fba-4339-88e2-fbf8f7c524af | 282 |
| PRAD | TCGA-EJ-5510 | 95ce00b3-cfc9-4400-ab07-de043b39cccf | ded2ef5d-8342-4b98-be7e-ca384611f209 | 143 |
| PRAD | TCGA-EJ-5511 | dfa7170d-3cc1-43b7-9d24-7be107cf2896 | 22de66f2-6831-4ec8-8ee6-15de1605d466 | 387 |
| PRAD | TCGA-EJ-5512 | 1df8e29c-5be8-4dbf-abb8-f0ded9419a14 | e86e1bf3-6515-4cff-9811-7b19fe9b2d00 | 149 |
| PRAD | TCGA-EJ-5514 | 581bee6a-6abe-4b55-8dc4-63bb242df1cf | eb55f146-90ca-458a-a1ae-b7e8347c700c | 520 |
| PRAD | TCGA-EJ-5515 | 25dbfef2-277f-4974-a326-2747d9494a2d | d2477eab-8dfa-47b7-9212-c2d294698767 | 159 |
| PRAD | TCGA-EJ-5516 | 80ba9751-2e29-4949-a07f-6e3542df8770 | 62d826e0-8cb6-4568-9f74-02b23a1421df | 111 |
| PRAD | TCGA-EJ-5517 | add98c6f-07de-48db-b930-82504d0dd976 | 9b9eefbd-747b-48b2-9382-494a8c720a29 | 314 |
| PRAD | TCGA-EJ-5518 | 02ae3fcb-c1fe-400a-8172-0c62a8d4639c | 048d94f9-66cb-4acc-9246-166201f39dab | 318 |
| PRAD | TCGA-EJ-5519 | 75f448c7-b10b-4c15-ae49-4eb217c65391 | 79a51d91-ec23-4cd1-be9c-59457bbad94a | 178 |
| PRAD | TCGA-EJ-5521 | c9a43591-8317-4ff5-a2e9-33c98c6a6012 | a00f8885-cc89-43c1-bf1a-d15ddf152756 | 150 |
| PRAD | TCGA-EJ-5522 | ee0fc9f9-f7ac-4f93-a066-2751ffaa3959 | 63030ee9-29f8-4955-8347-16e12ab169ca | 172 |
| PRAD | TCGA-EJ-5524 | bc7b862d-c88b-41f2-9061-6c4895d6b5be | f7ef709a-7e6b-4b7e-9827-28d64d2cedb3 | 195 |
| PRAD | TCGA-EJ-5525 | 10ab041f-88f9-463c-873d-508073d1672b | 7a9972fa-ec25-4ba0-bfea-8a15187015dd | 348 |
| PRAD | TCGA-EJ-5526 | e57f50b5-06be-4be0-8b02-706bfb439f25 | 7101d4c4-2636-40d7-85dd-c3036f889b7b | 260 |
| PRAD | TCGA-EJ-5527 | 62c0aaa3-44c3-4b25-b1cb-8e6196d66189 | 38e4be78-cef4-4c22-9407-ba48b34614f5 | 232 |
| PRAD | TCGA-EJ-5530 | 83b4e2b0-670b-441b-9b17-fa5c1da740c5 | 9722e61b-009b-4167-927d-68a6e13d0ead | 298 |
| PRAD | TCGA-EJ-5531 | 0993d456-af78-4078-aa16-bd810177c278 | 4962332e-85c4-4f65-959e-90f2e6169742 | 204 |
| PRAD | TCGA-EJ-5532 | e7c833a1-c77a-4c65-b268-1db73a7028e3 | eb3898c4-6098-4e04-b474-26ff917324b6 | 291 |
| PRAD | TCGA-EJ-5542 | 45ba9a12-204e-4ab8-b544-61f2519b176f | 1e52051f-16ba-40b2-bbf3-8cf3fd858f90 | 272 |
| PRAD | TCGA-EJ-7125 | c476525e-5147-4afb-bed0-f9ad34226240 | 956fdfd4-adc9-438d-8935-bb8a0626167c | 273 |
| PRAD | TCGA-EJ-7317 | 7c1cef2b-4606-4c6c-ad6d-b49ca86b9f74 | 15df0c4b-18fc-47ff-a394-3f998afafcbc | 454 |
| PRAD | TCGA-EJ-7327 | fe462cbb-5d52-4a0e-8cb8-a7e6c0f39c7c | 27f26a58-df63-4203-aea9-f907fc527c8c | 403 |
| PRAD | TCGA-EJ-7328 | 09a1120f-21a6-48c2-a9e8-3b229a16fc7f | ad1bcbc2-4111-46be-aa25-6f8222612d52 | 319 |
| PRAD | TCGA-EJ-7331 | a85ad241-998c-4d85-a63b-903ddeb62201 | e78549aa-ba12-42c0-b788-639fb85774a8 | 262 |
| PRAD | TCGA-EJ-7781 | bf6c4719-7b88-4975-9591-8bd34c19e3cf | 710510b5-d614-4a34-984a-0cb67724df03 | 333 |
| PRAD | TCGA-EJ-7782 | f92312ba-ec6b-477e-a08c-12c8af6d1775 | 978aefd3-3013-45dc-a27f-b3b3a3352296 | 844 |
| PRAD | TCGA-EJ-7784 | a2c5ffab-f279-44fb-a025-079de410f1c4 | 9c78d3ff-5257-47e3-86d4-8c7bbdfc5130 | 398 |
| PRAD | TCGA-EJ-7785 | dfbfaf44-ade6-4c48-88fb-a3898d878853 | 138e80f6-23a0-459a-9177-6dd8c1ce4b69 | 263 |
| PRAD | TCGA-EJ-7791 | 60424977-6a58-4b53-9607-4cf61abb2c55 | 3a9a4363-1e8b-44af-ac8d-ca5532f1ed07 | 130 |
| PRAD | TCGA-G9-6329 | 1971b9d7-8a67-4447-835d-23525622c037 | b3789c8a-7d5b-4bac-9ad7-4d8c002c629c | 269 |
| PRAD | TCGA-G9-6332 | 91b58182-30c5-4b45-bf21-ee32128e4ad0 | 4e876235-3367-4541-8b5c-432145854a49 | 227 |
| PRAD | TCGA-G9-6333 | 0e4674b7-f8ed-422c-909a-b2bfdf58ecf1 | ead27538-b8e3-4f26-905e-764c0be3f524 | 257 |
| PRAD | TCGA-G9-6336 | 69760af4-102b-4920-92f4-1424ac8db32a | 1b1f382a-4dbf-432b-88ed-e94a3d9a55b3 | 186 |
| PRAD | TCGA-G9-6338 | 3c22fc65-7067-4d13-a148-5e953f3a7097 | 618841c7-8398-44d8-bf70-1275e50d01cd | 246 |
| PRAD | TCGA-G9-6342 | 2e8198a3-6874-4ea2-9d85-405122d87da4 | 28236670-c301-4b5e-a898-f0d264f00b8a | 204 |
| PRAD | TCGA-G9-6343 | e413f861-4cba-4bf6-a805-aff9ecf4eb3b | f331491f-dba0-4053-b733-d52da4ca549c | 238 |
| PRAD | TCGA-G9-6348 | 32ce4cfa-ee58-4c85-8e6b-9e7fd9bb6b8e | 4c26631b-a745-412f-890f-793dd4b04e14 | 128 |
| PRAD | TCGA-G9-6351 | ddf40c8c-cec5-4f4b-853a-1541566f51b5 | 8cd65f12-8354-414b-a45b-9cc78613df38 | 101 |
| PRAD | TCGA-G9-6353 | 86ec03a4-f08e-474e-b887-c5e06822b414 | e230f78f-00cc-4bd0-a15c-369e9ceb0adb | 271 |
| PRAD | TCGA-G9-6356 | 4c78b0a9-d4e9-4656-95bf-0306c7bb406e | c966ad1f-2ded-40de-950e-0b9ee42a5923 | 151 |
| PRAD | TCGA-G9-6361 | 4fc66c05-8bf3-4a56-8bfe-6d20c3195258 | ae16cf88-3acd-4765-9960-e47c371159d7 | 194 |
| PRAD | TCGA-G9-6362 | 04834dec-1ae4-4a26-956a-f51c69d6c324 | a86b9f92-8f31-484f-9706-b320c5f42cfd | 162 |
| PRAD | TCGA-G9-6363 | a64da28b-c092-4dfb-9503-06c15e566ac3 | 2a69deb3-9543-4506-b9b1-31593b6cb7d5 | 249 |
| PRAD | TCGA-G9-6364 | 01dfbaf5-2c53-4f89-97c0-f845932cfde5 | fe1be0ad-62e6-4e97-b330-8f32afa551a0 | 174 |
| PRAD | TCGA-G9-6365 | 9d7979c6-8197-4347-806d-4bb667d5cb2b | cc8dcea7-b44b-4a7d-a9ef-f98a23c37e9a | 139 |
| PRAD | TCGA-G9-6367 | d4203e0e-c16e-4638-a239-378ffed98124 | 0c245fbb-c69b-4134-9b5f-883401c73269 | 111 |
| PRAD | TCGA-G9-6370 | 7b80e96d-c20b-4080-9382-26a778397c15 | 843714e6-d7a1-47d1-b3e1-9bc71c2e21ca | 197 |
| PRAD | TCGA-G9-6371 | 46c365e2-d78e-45ed-8356-c9ad57629c34 | 9e568f08-d30c-467b-a69d-7e89d377c1a7 | 165 |
| PRAD | TCGA-G9-6373 | b8d1f15f-e927-4a6e-8154-620ccb08e6f5 | 245d306c-0874-4b09-84db-125f16769148 | 145 |
| PRAD | TCGA-G9-6377 | 126d92ca-d172-491d-9366-11555d88901b | 82468f85-9845-4b13-b10d-4118b20e9e48 | 226 |
| PRAD | TCGA-G9-6378 | cc75d8b4-ea9f-4b89-9c12-d370ba7a227f | 0d86bec5-370a-4a3d-a9e9-120faf1ed159 | 145 |
| PRAD | TCGA-G9-6384 | 75df59ac-7525-45fc-843a-5210bd036dca | 723bbf9b-83eb-429b-ae56-ab0fce6478b2 | 103 |
| PRAD | TCGA-G9-6385 | 53da3cf6-b29e-4b1f-9d51-c1f9648b6634 | 40e5463c-dc65-437e-a884-d99c34009f5a | 119 |
| PRAD | TCGA-G9-6494 | d55338b0-a49a-4c6e-ba7d-757df0a55dfe | b30a02c0-4047-4759-bcc8-ae1a36aec49d | 143 |
| PRAD | TCGA-G9-6496 | 9bf9f3ef-b79e-4a93-8df6-ab2b9a3f24aa | 6bf8c890-7563-4945-80e3-504f41b07661 | 100 |
| PRAD | TCGA-G9-6499 | db7d4277-9d52-4996-ab7f-5e6ddc01c234 | 03cd9985-d047-464a-946a-1992ac4c8c1c | 235 |
| PRAD | TCGA-G9-7522 | 07b2c012-7fce-49d8-a6b8-6d5eb4907508 | 6544c47e-8718-4c5e-8c3e-94577b766269 | 175 |
| PRAD | TCGA-HC-7075 | 8f4cf0ab-38b1-49e1-8c15-82f9d02b5f9f | 798928e0-e49d-420d-8f4a-d0345f5fe570 | 1,138 |
| PRAD | TCGA-HC-7077 | bea10bd4-1049-4162-bb70-0bf5073199c6 | dd0dd624-e056-45af-8e61-7badb7a7fe56 | 283 |
| PRAD | TCGA-HC-7079 | 75a8bf78-150a-4c1f-99fa-41db038cf5a5 | 105059d4-8eb5-4f7a-a63f-838e9c234d43 | 294 |
| PRAD | TCGA-HC-7080 | 4012de79-f9d8-4ee6-ab04-538482e6a66c | eec2b211-fe5e-4169-883d-c614c804bdfd | 322 |
| PRAD | TCGA-HC-7081 | f44b51c1-3e9f-49b0-817b-b31a58264c39 | 8696b1a7-d544-4806-95c3-83dff3aa5508 | 346 |
| PRAD | TCGA-HC-7209 | d228a749-988a-4fcb-9f15-e334295216fb | 2c1969af-caf3-449f-9e39-f2387807413f | 404 |
| PRAD | TCGA-HC-7210 | e6934494-d8e1-44c6-a343-30a6aa2e6cbc | 51025f8f-6dc7-4e50-b60b-bbb7bc5cb367 | 229 |
| PRAD | TCGA-HC-7211 | c6d8e1f6-600a-4387-a51a-8812fee4f285 | 05e04c32-8191-475a-9d50-3fa50aadb8d5 | 385 |
| PRAD | TCGA-HC-7212 | 4ff229a3-6637-4fdc-a3b6-079e28773187 | e9912ef2-dbc1-4b69-8688-122162432c0a | 272 |
| PRAD | TCGA-HC-7213 | 14482ee6-0af5-43ad-bc2b-c9574446f6b9 | 21be48c5-46c5-48a4-ad57-a4703154b4bf | 370 |
| PRAD | TCGA-HC-7230 | 40ad0094-93d2-45aa-a8b1-070359637c85 | 0f575782-1639-413a-a7d0-e25ba87f997f | 199 |
| PRAD | TCGA-HC-7231 | 4b0f4fcc-c7c6-44bc-95d3-e693b0fbc7c7 | 9ad2873c-4fe1-4a32-9c57-921c2e4c87d2 | 274 |
| PRAD | TCGA-HC-7232 | 26e5268d-e95b-44da-ae7c-dfb6fe50a127 | 650b3a4f-0196-40bf-88c0-7b01ad67b377 | 278 |
| PRAD | TCGA-HC-7233 | b16d1760-889f-44f7-a1c1-1dfa3f31efc3 | 5e685d94-c8fe-4fd0-945f-268e1a8d4d77 | 4,206 |
| PRAD | TCGA-HC-7737 | 855b7ac8-18b6-4ec4-9471-8d12a802bf46 | 0f09b06b-7747-47b8-be3c-4e592ca62c5d | 276 |
| PRAD | TCGA-HC-7740 | dd2df8bb-8035-4f08-b3d4-d9ee910fd0ea | d72a0569-8a57-429e-8b29-e42f85c3a252 | 169 |
| PRAD | TCGA-HC-7744 | 8589d977-d1ca-42ea-975f-c3e664d57eba | d577528b-ba13-4fce-94b9-4222459bab0c | 1,712 |
| PRAD | TCGA-HC-8258 | 696d5596-7630-4286-a1c4-8a3ea96ddf16 | 9a96b90c-ef2a-4dc8-a554-8c646275ad73 | 196 |
| PRAD | TCGA-HI-7168 | 97c2b4ae-c671-4cf5-8b44-5c36e7e8fb57 | ab5dab2c-9975-4682-921c-6e013cd929bb | 268 |
| PRAD | TCGA-HI-7169 | 4ec05d23-3f9f-473a-b292-63ddb0e7d37f | 070dd820-fa5a-42a2-8d3f-3a562df06f24 | 512 |
| PRAD | TCGA-HI-7170 | 4eefa9f0-575b-4039-912a-7c0e58f4e508 | 3469af72-4b83-4fe2-bc08-221168d68d84 | 209 |
| PRAD | TCGA-HI-7171 | 0dc8ef7a-58a8-4b25-ad57-4c31592151e2 | 966d1a07-f599-4c0f-86a7-d4b4616c7bc1 | 271 |
| STAD | TCGA-B7-5816 | b9711ffa-01b8-4434-ac32-445717298e6c | c42706d2-adb2-442e-a7d3-43fc52f02c43 | 256 |
| STAD | TCGA-B7-5818 | 9924779b-8261-4ea6-9925-28dd5a0aed8a | 144edb0d-1d38-420e-a1b5-ea974935bacf | 1,977 |
| STAD | TCGA-BR-4187 | 1da3a244-39ae-4b4d-baed-e4d9b312f777 | 6ba29a1c-c406-4572-bb2e-69bb70f46dfa | 351 |
| STAD | TCGA-BR-4191 | d71fa3ea-a7fb-40af-a7b9-64ceca39e135 | 78b2b7ff-d302-4923-bfa4-f8d169667d90 | 264 |
| STAD | TCGA-BR-4201 | 359db0d8-a78f-4d9d-9a3e-9f1a04472df0 | a6dc7c10-d45e-4f8a-a8fd-bd928d8f554c | 567 |
| STAD | TCGA-BR-4253 | 90590609-4f3e-48b4-98e5-963c1c97f057 | 6237a524-9f83-4bf5-8956-06e3e60438b1 | 1,538 |
| STAD | TCGA-BR-4255 | 5085f26e-d490-4a13-a172-4cac0f68a682 | e7981f49-1074-4ce6-820c-3630abd6e8d7 | 551 |
| STAD | TCGA-BR-4256 | 6e990147-2463-4d38-a785-6c76ae1dbc25 | 9e52ce3a-8344-444b-b624-ae3604caa8be | 130 |
| STAD | TCGA-BR-4257 | 5e38d8b4-97fb-407a-ab06-2f002456472d | 40fc7a6d-ebf3-490f-ab9d-19e641ea0dbb | 116 |
| STAD | TCGA-BR-4267 | cd48e485-3a3f-4eec-92e7-f1c52c0866f1 | 1ec7ecd8-69c7-4a5f-a296-4ebb9e591729 | 2,996 |
| STAD | TCGA-BR-4279 | 0fa8a7d6-6636-4135-82f6-21c9df279b16 | 1fc5b32a-8266-4f98-8444-a219ed9631a2 | 737 |
| STAD | TCGA-BR-4280 | 82a3e711-b833-4607-8007-dd29e00eea85 | d6f3f036-2ddf-4ddb-81e4-3a1c7a977bfd | 61,711 |
| STAD | TCGA-BR-4292 | a7dacb77-1a29-40e8-a758-c37b5bc38631 | 763db891-de7a-4a7f-8608-d70ab806e102 | 2,361 |
| STAD | TCGA-BR-4294 | 23c036ca-ac6a-45dc-9100-558b2249f3bc | 77aba0c4-8048-441c-89af-698eee7d38aa | 1,075 |
| STAD | TCGA-BR-4357 | 4bc9f55b-1edb-49c9-be1a-1d00a108b99b | 9c4e7319-08e6-46e0-a33b-68d2dc8a6926 | 694 |
| STAD | TCGA-BR-4363 | ab5af808-6239-48fa-ba17-b80aa3fc5e14 | 06da121c-66ee-44be-8a1c-58e736f82e55 | 106 |
| STAD | TCGA-BR-4366 | c4b9424f-d0da-48ae-8723-69cbccfdefc6 | 9179774b-7bf8-4aa0-886f-2917234201eb | 145 |
| STAD | TCGA-BR-4367 | 98785371-4af3-4d04-acc8-5c53c8076618 | ea05becd-5829-4f1f-8dd3-56dedaa95846 | 139 |
| STAD | TCGA-BR-4368 | 138fe28f-1055-4872-99b0-71ca92e3494a | 8d384344-e3de-49fd-b94d-7bf0f0c03920 | 191 |
| STAD | TCGA-BR-4369 | cbba5183-9ada-4363-8f98-bf406b64f6d6 | d5894343-a206-49f1-adce-53c336fec1ee | 155 |
| STAD | TCGA-BR-4370 | a1feb789-c851-4608-a5c1-382bb432c4c7 | 2d7a67d4-6bea-4ea5-a59c-edfc3adb7ed5 | 176 |
| STAD | TCGA-BR-6452 | cfd9aa7a-7f3d-4b70-9359-2c93f0238361 | b5d222fd-498a-40a1-880d-322d0a7a1323 | 132 |
| STAD | TCGA-BR-6454 | e344c34d-da8c-4145-8a02-b9ae248ca3a3 | e6845469-d068-4e77-bc48-b17e5ad0c4de | 77 |
| STAD | TCGA-BR-6455 | 719ee3a5-cfe9-48cb-9e48-e11e49bd217c | 63bba89b-b557-47c5-9c7e-894a64d517e2 | 205 |
| STAD | TCGA-BR-6456 | 9830e4a0-d8b1-4166-8375-49851288cf1f | 4fcd3184-8cb8-4097-8550-b9905d13adb4 | 233 |
| STAD | TCGA-BR-6457 | 695f7145-7735-48f7-98e8-99b86bb936f4 | c937c321-5e90-4ec8-925e-e1cd02e6d5e5 | 229 |
| STAD | TCGA-BR-6458 | 545acf44-8e16-46b9-b35a-c860c08b416b | 0fb3d589-66c9-49b8-9263-a04b09abcdd4 | 301 |
| STAD | TCGA-BR-6564 | 6ee5a220-be0b-4909-bd37-5a6d8c53177d | 8b88ec34-0f0a-4fe3-aa10-42a0ee7e46e5 | 4,177 |
| STAD | TCGA-BR-6565 | 8cfafa49-f561-4e59-98c5-b9a816534417 | 99084bc3-84ee-4ccc-9c54-1b5a74af410b | 243 |
| STAD | TCGA-BR-6566 | ae711b74-3be9-4959-addd-03427c07f35b | 929fba9b-a5db-4260-beee-93a5f1be80ac | 368 |
| STAD | TCGA-BR-6705 | cf5ffdf8-e424-4a51-862f-816cba0c92e0 | 7fbe76eb-2522-4271-89b9-61797e2e206d | 382 |
| STAD | TCGA-BR-6706 | 175293ed-2943-410e-8c01-a61c21354054 | f98df3e1-6bfd-4745-9c3b-412598e8d952 | 310 |
| STAD | TCGA-BR-6707 | e2babdbd-1849-48ed-8726-1e3788be6064 | a99c1ab3-7b20-46dc-b74c-1866a6aaaf62 | 491 |
| STAD | TCGA-BR-6710 | e0e0146e-bf19-42c8-801b-c5a6ed564c5b | a4f4bc0d-7d51-4c01-b0c7-98f206980452 | 235 |
| STAD | TCGA-BR-6801 | 48843aff-35a1-472b-a298-6489ad1b7de7 | ad1f6971-d904-467d-a98d-2bc17299540c | 579 |
| STAD | TCGA-BR-6802 | 73cec875-41a5-40bd-9d13-9e1ed4dd64d0 | 2c69e92d-3297-4add-a687-68ad89040c66 | 842 |
| STAD | TCGA-BR-6803 | af361f62-8dd0-4336-9fa9-41cdc800b18e | 8e0aa4a2-6371-4260-ac1c-249ff5c4f1f9 | 334 |
| STAD | TCGA-BR-6852 | bff2f642-516a-4072-95aa-f213a4555f79 | 9f843373-3bfd-4a53-ba12-0a753b47d24d | 631 |
| STAD | TCGA-BR-7722 | 2ca0a03f-9d9d-49da-98e7-b0ade46e5b18 | ebf8678a-8dcf-4977-8266-e030e89a984a | 2,957 |
| STAD | TCGA-BR-8373 | cbbdb4e0-d400-45ef-bb4f-790b2cc9cb87 | 2828f4a2-a0e8-4b53-9320-de0ab5598e09 | 7,566 |
| STAD | TCGA-BR-8381 | 8d5cdec5-779b-4ca8-a709-f2fbc527642b | 061589aa-8b3c-465e-a354-c6d9ba618e35 | 381 |
| STAD | TCGA-BR-8486 | 09ea179f-e2fc-479c-9b04-7c3ae8034874 | 4a10b9ff-71a3-46a4-8cca-d0b4f7dfec66 | 2,614 |
| STAD | TCGA-BR-8682 | 36f56254-f585-4353-89b4-00cb5cfa20eb | 7b22ed9c-2c20-44ef-9443-e6ad328825ba | 1,195 |
| STAD | TCGA-BR-A4J4 | 2cdabae0-c924-481e-ad7e-bbe3f72f5c79 | 183879a3-1761-4b9d-a0d2-fd2b2322599c | 2,108 |
| STAD | TCGA-CD-5798 | 5b2c6e2f-0b61-49e1-bef5-6d7408918d47 | 89241089-c1c3-413f-8996-a3f43497f231 | 254 |
| STAD | TCGA-CD-5800 | e62fc018-7ddb-4d26-b187-0781aa2ba772 | f7df3714-78fc-460f-8d69-6870f63c1f49 | 228 |
| STAD | TCGA-CD-5801 | 5196b0dd-1d41-47f4-84a8-999f211443f4 | 0127bc2c-34be-4051-8211-7ccdc8529b37 | 218 |
| STAD | TCGA-CD-5803 | 1002025d-03b9-46ab-988a-576f59957cac | c5245994-d084-4ea6-b869-e4f9a02b9ad8 | 290 |
| STAD | TCGA-CD-5813 | ccc0d204-3529-43aa-b108-62d12f6465bd | 7cbca34e-4195-452d-84af-5ae9596c63af | 852 |
| STAD | TCGA-CD-8529 | 65281f4c-9277-4d03-8566-75feb73c114d | 244c2dc3-6a22-4851-9e9d-ab106781e439 | 702 |
| STAD | TCGA-CG-4301 | 7d1cd13f-5ac5-457f-8e7b-89cabb2463c6 | 6015c990-2a38-459d-9d91-64c1561fac75 | 303 |
| STAD | TCGA-CG-4304 | 4fbdbad3-32cb-45de-990a-1337f32a25fa | 2a7989dd-2b9c-48c4-9b6a-ab9a5864b004 | 129 |
| STAD | TCGA-CG-4305 | dad3a237-d784-4731-bbef-c4d3dce49fa2 | a9425aae-9f82-4927-bef0-4b29926935d2 | 885 |
| STAD | TCGA-CG-4306 | 97a524b1-432a-4c20-adcf-5e45ee168834 | c0a0bf03-12bc-447d-9f12-b8881dee8f37 | 847 |
| STAD | TCGA-CG-4436 | 0b869c68-28bb-4675-ba6c-6d20a89a1a49 | 3eeefb8e-1bd7-4739-b4c3-b7695bedf7d3 | 236 |
| STAD | TCGA-CG-4437 | ff02c427-3a7b-4e29-9ddc-0b8034807ad6 | a6985d3e-05da-443e-9b5e-7098618981c5 | 245 |
| STAD | TCGA-CG-4440 | d72343ff-91f0-4705-a398-914c5d2344d9 | d6eefe03-313a-4546-8bd5-62d1cb79c1c5 | 99 |
| STAD | TCGA-CG-4441 | d3542e45-2b25-406d-976d-5d4c63e4839e | a450bc30-69de-4e0a-a561-2145c316b5f1 | 1,747 |
| STAD | TCGA-CG-4442 | 05a46a94-fa5c-4e76-b887-07ec0fa0af0f | c3e26eec-0792-4737-ad87-9d9a95fa438f | 27,436 |
| STAD | TCGA-CG-4443 | d15f98aa-3849-4230-aea5-46a959ece810 | da94bd95-a2f8-436a-8738-831e4e1ba8b7 | 5,420 |
| STAD | TCGA-CG-4444 | 37421315-54a7-4a59-9939-4706dcd66019 | 71fb65dc-8cfc-4004-a794-1a756c04d829 | 163 |
| STAD | TCGA-CG-4449 | c8b033d1-998b-4fee-88e7-33c14e8c3ffc | e078d104-2b36-461b-9e5c-af5b8b2fc89a | 63 |
| STAD | TCGA-CG-4460 | 1d341d0a-b2f5-4f35-b60d-7c87f2a7dc78 | 88366088-e644-4828-abfa-01a45fdb23d3 | 122 |
| STAD | TCGA-CG-4462 | 1bcd6b3d-eeca-4c1e-bf6f-52c0a6b151ca | c1ec8c57-0cf2-4bfe-b6d3-a7ed77f918c4 | 302 |
| STAD | TCGA-CG-4465 | cb9d270d-5328-46d7-b7ff-cb564fad9175 | fac76410-c449-448e-8f7c-c33a3e3cadc5 | 1,338 |
| STAD | TCGA-CG-4466 | a8cb52d3-64c6-4fca-a59c-b177968a7489 | af58bea8-9aeb-47c8-946e-6b197b06ea79 | 553 |
| STAD | TCGA-CG-4469 | ed99ca3f-0ae6-43ab-a171-dc2774cf52e2 | 8c7f32c0-34e4-4aea-8603-cce8acbc1b82 | 1,886 |
| STAD | TCGA-CG-4472 | f1d39d03-103f-46f0-8fbf-ef9e944d68ad | 0f6fc60a-9288-4cc4-9b91-35bb56cbad3f | 1,520 |
| STAD | TCGA-CG-4474 | 65c20199-b73c-42b2-b95a-3995b59e5d20 | af9433c7-f0e8-4b30-ba69-0ffec543d03a | 1,208 |
| STAD | TCGA-CG-4475 | 8d322c4d-bfdb-40cc-a222-87ff33a87718 | 8e28c3d4-9556-49b7-bde1-8f7fe76e98fa | 471 |
| STAD | TCGA-CG-4476 | 87a698be-8fb0-4ced-842b-fdc852b98aa8 | a6ccc4b9-9b2d-4960-900b-4e637c691eee | 461 |
| STAD | TCGA-CG-4477 | a2b4b066-6bb7-401c-9ff9-e88bee42884b | e9116b76-5b39-48ec-802b-70e4c7eec731 | 19 |
| STAD | TCGA-CG-5717 | 34ec4e75-2398-4904-adf0-34500892e1a5 | 6aa2ef88-18e7-4056-b37a-72a9acd0fed9 | 105 |
| STAD | TCGA-CG-5718 | 98032563-24e7-466c-9f90-1ba67975f2f9 | 30044c1d-1cc7-49e5-9bed-dbb5168be8f4 | 189 |
| STAD | TCGA-CG-5719 | e6237df3-92e8-4e8b-92a4-b09d3515f16f | 74d5e460-86bd-441b-9409-00b8f0248eb9 | 131 |
| STAD | TCGA-CG-5720 | 6cf7bbf9-63f5-40e8-b595-c9012a1825ba | 75fe7913-8432-492a-a33c-286c06164f35 | 235 |
| STAD | TCGA-CG-5721 | 450e30b5-bcb4-4169-a48f-bbf3db7feda2 | 259d4033-0c05-4f3b-ab2a-b0c7018bad33 | 1,767 |
| STAD | TCGA-CG-5722 | 05762233-5294-478d-b462-14b931579f63 | 53d1fe8c-b529-4f7f-a9d7-672085ce2fab | 238 |
| STAD | TCGA-CG-5723 | 9e55e0ac-b9dd-42dd-afe7-d6f93c1dcc88 | 6bd76290-af9b-4812-8b89-e9a8f8416fbc | 2,025 |
| STAD | TCGA-CG-5724 | f2f2a0e8-364c-4603-8030-6f0f803fe21c | 4289e6b1-841a-44f2-adf3-c4729a04518c | 2,371 |
| STAD | TCGA-CG-5725 | 61968fb0-fb5d-48fa-88eb-4cd76a6b4456 | 42216242-2d3d-4bb9-9b00-f0d50042790a | 418 |
| STAD | TCGA-CG-5726 | 87491384-bbcf-4127-8cf0-59e692b8e37b | 3c2e1753-936d-4794-bf43-9af5aa4b97ce | 2,135 |
| STAD | TCGA-CG-5728 | 557597f6-e3e0-4ac2-a626-902dc5b8709f | db35e103-ab69-44f7-b5c2-913ab5032554 | 1,067 |
| STAD | TCGA-CG-5730 | 61f3e14a-530d-43c8-a795-a6aef7d031e4 | 7f327ebb-598c-4874-93d0-5f525410a382 | 3,203 |
| STAD | TCGA-CG-5732 | 3bc07391-1ef5-4bbf-9614-e3f31b8448c9 | c4e25e33-1ac2-427d-adeb-7d6cfc0269a6 | 147 |
| STAD | TCGA-CG-5733 | 4b31f837-59fa-48a7-bfbc-475e1251bd07 | f72098d0-5e3e-49e7-a5e9-5bf9f620f2c0 | 2,166 |
| STAD | TCGA-CG-5734 | 4cac9679-5c5e-4b51-b7a0-0d82ab2c2237 | a90934d5-13a4-4673-80b4-25d0755eb14c | 238 |
| STAD | TCGA-D7-5577 | e417c8be-9c96-4e81-a2c9-ac3f23791347 | 9f188367-53c1-43c9-a6bb-c59e0546fb91 | 706 |
| STAD | TCGA-D7-5578 | 306aee2b-5fac-4825-952b-d838746530a5 | a07d68ed-d63d-42c7-80dc-29e0c8cf5367 | 1,052 |
| STAD | TCGA-D7-6518 | d4c1a136-c2f2-4cac-9b15-79669253b6e8 | 3158223f-89b3-425b-8b91-8b386fd4e604 | 228 |
| STAD | TCGA-D7-6519 | 66b2b99b-ac75-4068-9d05-34e6dbe9721d | 95e3ee3b-6ed5-4d0d-a7a9-64ae3139dab4 | 443 |
| STAD | TCGA-D7-6520 | 46b79d94-56c0-4b23-955d-2395a50ecaf9 | c35f64e9-94a7-4b34-aba4-01aea848176d | 200 |
| STAD | TCGA-D7-6521 | 72a895e9-b1b0-4b79-a5e6-6bae340e8798 | eb009690-3197-4b8c-9b74-0020480845fb | 320 |
| STAD | TCGA-D7-6522 | ac3cc7c3-e03a-4218-b297-d6c525338753 | b268b352-4bb7-4b44-8367-f621ab6d372c | 311 |
| STAD | TCGA-D7-6524 | 67f702e7-84ad-4d23-90e7-03040d417e4d | 53a3658a-e190-413c-904b-1f8bf1ddf4ee | 330 |
| STAD | TCGA-D7-6525 | 03097d75-d0ad-4e1e-8a9b-5157a768cc05 | f95ec729-b7a8-4b1f-9c2c-ef00b3a74a77 | 1,624 |
| STAD | TCGA-D7-6526 | 0a9e4a52-46b9-4c50-9dd3-9418ed5569e2 | 9470ad93-71f4-4d5d-a26b-8cec32d802dc | 743 |
| STAD | TCGA-D7-6527 | d1534b7a-a5b2-4083-9736-d1ad62a97ad2 | 0c16032f-bf10-4293-b1be-a0f6fe286885 | 936 |
| STAD | TCGA-D7-6528 | 478ea84f-df8f-4d1b-a7da-ab1ac7751e97 | d4ea96e7-6bf3-4ac4-88ad-e0199ddef1b7 | 628 |
| STAD | TCGA-D7-6815 | 6265f1bc-c42b-4073-b83d-4f13fbe9da6c | 48e964d1-7020-48a9-945c-f3409790f8ea | 5,336 |
| STAD | TCGA-D7-6817 | 9d6376df-5da8-4903-ac82-a2c69750107a | 3b400a10-191c-44f7-af35-4b9ea1cb192e | 328 |
| STAD | TCGA-D7-6818 | 5a217dfc-ada8-4dec-8f7c-06be0123b9ef | 327fce3d-289c-4d30-b812-d43fda314ac6 | 297 |
| STAD | TCGA-D7-6820 | 8b99bd62-c3d1-40c5-af7c-b1976df9622c | 99cd87aa-0deb-4ace-86d1-adfb8dbee226 | 60 |
| STAD | TCGA-D7-6822 | b8f8d06c-7bbf-4d0d-8f5a-e3012379df95 | 53456e48-e181-4f75-be35-28167c1fb50b | 7,512 |
| STAD | TCGA-D7-8570 | 4b984072-ec9c-4924-a922-f7c4f5c2eacc | 17649637-fcdf-4c26-add7-6820145afb75 | 1,032 |
| STAD | TCGA-D7-A4YX | 004a24e0-0e6d-4811-aac1-344ce1d54320 | 730adaf7-bfb4-437b-af0e-a94fb4379e88 | 3,255 |
| STAD | TCGA-F1-6177 | 347b5bd7-20b6-4635-ae86-d43256011296 | 1edb61f1-07b1-4e73-bffb-71fa3fd004cb | 102,759 |
| STAD | TCGA-F1-6874 | 734d68fe-de19-4aef-8031-8ba255233eba | 478d46a8-c228-4b0d-9d5b-c433514e64d3 | 778 |
| STAD | TCGA-F1-6875 | b6dcf3d5-80e4-40e1-a164-4697d1de7670 | 6dcad824-de10-458c-a0fe-32974ab948e1 | 10,135 |
| STAD | TCGA-FP-7998 | d124dc46-96b0-4101-b8a6-3e4ebf4862c9 | adb24a00-e8b6-40bc-8a22-3e8559a20903 | 385 |
| STAD | TCGA-HF-7136 | 09ada83c-8149-4ed4-8e85-b2c9be12c8d5 | c768ee2b-bcfc-442a-9697-6fd379b74814 | 7,769 |
| STAD | TCGA-HU-8608 | 58876087-e158-4469-af68-c91fe264ca8e | 7316c2ec-ff33-4f79-ab93-c88b253a0f5c | 752 |
| STAD | TCGA-HU-A4G6 | 01564622-cd3b-4f99-885f-ac2d8ca424c3 | 6a591fcf-0fe6-406e-b57a-76dc16219647 | 6,291 |
| STAD | TCGA-HU-A4H0 | 787fd694-c915-406f-a8a1-0ca998d01d93 | f597333e-382e-430c-b763-7d725833e28f | 2,298 |
| STAD | TCGA-IN-7806 | 65b3a7db-195a-47e8-804b-a1d4a5df79aa | e74ea4dc-0eb6-4d88-840f-17d1762d2c46 | 6,516 |
| CESC | TCGA-C5-A0TN | f860da6f-1cec-4913-aa35-b32670a1734f | f69c19a2-3142-43d4-b62b-2bfb91c76ac3 | 3,158 |
| CESC | TCGA-C5-A1BF | 913cddd8-a6ed-4824-8b2c-02f3160028d6 | 7a3dd7ed-42f8-4430-94ff-ce66abc93b74 | 3,760 |
| CESC | TCGA-C5-A1BJ | 0990b367-e2b6-4b2a-8ca1-dd9efab1866a | 5543916b-b5c3-40b9-9614-537c7ad6e235 | 889 |
| CESC | TCGA-C5-A1BN | df3a48da-646a-4834-969a-131455b7baa9 | 85c2015a-844d-46ce-9d10-7eb30a07fb3c | 6,049 |
| CESC | TCGA-C5-A1BQ | 58d97d84-99da-4b08-ae63-7e49f4b861b7 | 16a3e11d-b1b5-40a1-adc4-2985ddceef19 | 453 |
| CESC | TCGA-C5-A1M8 | bcb9d9ed-cd43-4eb0-a3e5-f667a80c20b7 | f9f50bfe-4bce-47cd-8371-5aa09e5b1977 | 5,668 |
| CESC | TCGA-C5-A1M9 | e9ec290e-cad8-4af0-8b69-cf804598a951 | 955cddcd-6ab3-40a9-bb44-8095d3af1446 | 3,555 |
| CESC | TCGA-C5-A1MI | e32cf6ef-190c-4456-b8a7-b40b9c9c8036 | 5283bba7-623e-476e-bd42-b147b6b6f394 | 2,747 |
| CESC | TCGA-C5-A2LS | 3c012d75-0d17-4b4d-834f-cd4767f6b023 | 2f77e8f5-e810-408a-aa50-caa8f614c77a | 472 |
| CESC | TCGA-C5-A2LT | f6c6e1bd-966a-4f96-b8ab-e8219607f960 | d4e12dd5-93a7-4afe-96c4-470639a67f6e | 1,673 |
| CESC | TCGA-C5-A2LV | 559f9675-671e-41a5-b073-7782ab8b5542 | 5d6ed525-a289-497a-a138-6e7570dc1f9e | 4,579 |
| CESC | TCGA-C5-A2LX | ef574f7b-2b69-4be8-83f3-cc02df579a0a | 2ae4bce7-ffc3-4bce-a766-63d284bcab75 | 633 |
| CESC | TCGA-C5-A2LY | f5457c81-3a5f-450c-b066-b1dedb472c0c | 03ebae4c-d724-4d32-9b91-943abd271d24 | 4,514 |
| CESC | TCGA-C5-A2LZ | 19f84215-7f5d-4cf8-a6f5-e135327ccd50 | a618c67d-3bd4-49aa-a5e3-8bd38024e5ae | 212 |
| CESC | TCGA-C5-A2M1 | 374c6310-126e-48e8-8b83-145f75714a4b | 78a6a70e-fe67-4b95-a5eb-6bff4842969c | 545 |
| CESC | TCGA-C5-A3HD | 475d850b-49e8-4f15-9f59-8d65bf9cf22d | a955b684-6a8f-4334-8362-a5d192f147be | 340 |
| CESC | TCGA-C5-A3HE | 1d64c1f3-438b-4576-b2bc-74c1a26a2aca | 127e65bb-ad88-449a-a8b7-a3b77a65f010 | 843 |
| CESC | TCGA-C5-A3HF | 4b4bea19-4db4-4996-8037-4129a4494419 | 19a65231-8bdc-43c7-81ed-e92f884b6c03 | 112 |
| CESC | TCGA-C5-A3HL | 84c6e022-f737-4fae-a2c7-812ef4f00adf | 9b5f1439-6b4f-4a02-857c-75dd4c6d298e | 151 |
| CESC | TCGA-DG-A2KJ | 4399a247-c12c-480e-bc92-cf902a8a0a69 | 88d67937-e5d1-4d8c-a4b3-05cf8f2ddfd2 | 3,934 |
| CESC | TCGA-DS-A0VL | 2c9bc14c-a6d3-4389-8360-024e9c04a5d4 | cd6ccfff-e6b2-4bbe-bcf7-290e0e3f80cf | 2,109 |
| CESC | TCGA-EA-A1QS | c7f62ea9-e05f-40ea-9e4c-1771bd722b74 | 59f4f825-bca8-4cb6-b942-b3af019dd252 | 270 |
| CESC | TCGA-EA-A3HQ | 33c80031-f923-4aa0-adfa-9e229f413f83 | 15d3baa9-7081-4e33-9ce7-8280513fd8c9 | 160 |
| CESC | TCGA-EA-A3HR | a2c5a51a-5933-4597-b207-2c761c54f7c2 | f3670983-2ab7-4d5f-ba0a-78f0cba773a0 | 103 |
| CESC | TCGA-EA-A3HS | 684e83c6-0894-4bfd-a2ee-6b1877c74f82 | 66d157df-3130-4d6f-b945-b2b660c92060 | 334 |
| CESC | TCGA-EA-A3HT | e0e70b08-26f3-48d5-80c3-4836e9faa569 | 381a7580-473d-404d-80c8-deb66057b4fb | 329 |
| CESC | TCGA-EA-A3HU | 4f5312e3-89a9-41d7-a296-8b90e22cb408 | 512fb31d-ffcf-410b-bb95-1e25ac434187 | 326 |
| CESC | TCGA-EA-A3QD | d97f385a-fc08-43a1-9481-35f9ffdfcc05 | c62c9fa4-7e51-4f44-a9e7-66773a034e59 | 506 |
| CESC | TCGA-EA-A3Y4 | 32cd0b72-100c-4e6c-81f1-6be30738cafb | 6e7276b2-90e0-400f-b317-ce52a58361da | 374 |
| CESC | TCGA-EK-A2PK | 7109053e-8be0-4965-a5eb-a9bbab5ee53f | fb7abe42-2f5a-4290-9765-595b6bc19069 | 1,265 |
| CESC | TCGA-EK-A2PL | ca898dd4-73fc-4fa3-8d2b-3d4f27be4736 | b844ef40-47ec-47f7-a517-bb3a171bfc7f | 582 |
| CESC | TCGA-EK-A2PM | 332b29bf-f948-4e2c-bf51-f768f819e9e9 | c41a041f-a7af-40aa-b516-42ca0cca7082 | 1,834 |
| CESC | TCGA-EK-A2R7 | b998e76d-dff1-4454-abce-bbe7b5a6c398 | 86ab2e15-4d96-4350-b5cb-99b156e7ea61 | 410 |
| CESC | TCGA-EK-A2R8 | d050a7c3-8fe5-472e-bc71-4dff85991e92 | 8498b73d-be83-4863-bad7-ddd22874d071 | 870 |
| CESC | TCGA-EK-A2R9 | d25f9f12-7f03-4bbe-befe-f9250955975c | e1a04d71-458b-4484-b2a7-5c5dc63dcba9 | 3,394 |
| CESC | TCGA-EK-A2RE | 87fe03fa-225c-4d35-81a3-8b401d985fae | e89116c7-baf6-41ea-a3b7-90fbdd692255 | 1,994 |
| CESC | TCGA-EK-A2RL | 5848efed-18c8-413b-8a2c-9a63deec56ee | 59e94488-d19c-44a4-9cb9-0a2d4d5836ab | 2,354 |
| CESC | TCGA-EK-A2RN | def278f8-93eb-4373-a525-98b2cf12af7a | 2fca04bd-9b4b-4ad4-b36c-b9e0f2a2fcac | 120 |
| CESC | TCGA-EK-A3GJ | dde88a3f-8602-42a1-9b43-037c49cb3d34 | 5b9221da-4fdc-484f-867a-280b02e25e7e | 263 |
| CESC | TCGA-EK-A3GM | 2ec77e09-5fb7-4247-9027-1db441eab2b8 | 5eb98be0-3349-4f64-acb4-1eebb375d826 | 231 |
| CESC | TCGA-EK-A3GN | b9e845ee-3e61-4ccf-b14e-f8a6b8481bd3 | 9b083a7a-df5d-4937-a89c-341a4622b771 | 110 |
| CESC | TCGA-EX-A1H5 | 9c7919ac-894d-4ecb-8c98-b99765ec4bbc | 23af8d74-ce98-421c-aa2c-f0cb33494eae | 3,597 |
| CESC | TCGA-EX-A3L1 | 73141d26-4c6a-46e1-ab5f-b431f2f35c23 | bfcd19d2-9301-411b-856a-ee0acb688dff | 185 |
| CESC | TCGA-FU-A3EO | 31874bb0-fa47-43d4-8135-1be05e8163eb | 6577de22-c954-4658-98b4-6dad43c6b910 | 133 |
| CESC | TCGA-FU-A3HY | 30d23ada-985e-43b1-9cc5-9e3931905b07 | 311eb518-2d97-4352-a019-82bbc266a540 | 522 |
| CESC | TCGA-FU-A3HZ | 40af9367-bd77-4c64-98d6-d2e2c6e5c11a | d2ffd1d9-03a6-48ce-9dcc-4302224492ec | 696 |
| CESC | TCGA-FU-A3NI | e3d152ec-0ab5-4fb6-b2ca-ecdb79e43095 | 012f7e0c-b279-426a-9979-aa846fda7c90 | 354 |
| CESC | TCGA-FU-A3TQ | 87ede4e6-17e6-4287-8ee3-cde27de3aa7d | ea233e9d-f242-4136-9f61-e20710d8c141 | 380 |
| CESC | TCGA-FU-A3TX | 3692ce0d-daf2-4076-a86f-4d8e9ea25745 | 02fd7716-d363-4d47-81b8-16f0a83a944b | 438 |
| CESC | TCGA-FU-A3WB | 39124a1d-9323-40ae-aee5-6fb32cf4a565 | 51850ff3-c031-496e-9989-ba2d764676ca | 478 |
| CESC | TCGA-HG-A2PA | 2c605f98-0c77-45ff-bcea-2fcf6094ca2d | cd34db28-8cb3-4000-9b95-22210dde4e3f | 148 |
| CESC | TCGA-HM-A3JK | cc4a29a9-d389-4d17-b8c7-53fcc5731b22 | 7d7efc4f-e09e-4207-80d4-b6965a70903d | 259 |
| CESC | TCGA-IR-A3L7 | c50ab1fa-015d-4420-a67f-b023a5377b16 | 15c1faa9-4030-4527-9860-f1846bbb8e9e | 243 |
| CESC | TCGA-IR-A3LA | 96c944ec-9e55-4b27-aa91-b936b4d7cff5 | 5b0c885d-73d4-4af1-9c47-a5d59d6a0664 | 2,849 |
| CESC | TCGA-IR-A3LC | 8b84ea0a-0512-427e-8456-2facdcd978b9 | 52127837-efa1-4fef-8096-506ecc100e1b | 155 |
| CESC | TCGA-IR-A3LF | aba44461-5b2e-40f7-9e8e-262aee825e6c | d1073f97-3ce5-4d38-9ccc-813fee28fb63 | 368 |
| CESC | TCGA-IR-A3LI | 3a9d3f81-919f-44e3-a9cc-813b4dcaf027 | a2fb1563-5e71-4358-b452-ac8ecd9024d6 | 161 |
| CESC | TCGA-IR-A3LK | 1c6981be-5fe3-4898-ba63-e0cd0eecc5e5 | a4df291b-d7e9-452c-ab2b-52ff1db1051d | 429 |
| CESC | TCGA-IR-A3LL | 817ccf3b-ceaa-45cf-a938-67e531a9f57a | 4b74b0ed-9434-46b0-bf59-1a357318e495 | 186 |
| CESC | TCGA-JX-A3PZ | ecad03c2-29cd-48c4-8c37-7eade85970f7 | 1078fe30-c37b-494e-97ca-3359b7d8f630 | 97 |
| CESC | TCGA-JX-A3Q0 | 20d5be50-f8a2-406a-9415-bc65cdff0cad | fbcd3e16-2a1f-44e8-a46e-d05f1c5d52d1 | 1,103 |
| CESC | TCGA-JX-A3Q8 | e8f02d77-0077-4817-a34b-ab4ddd64bbd5 | 638ee124-3225-4587-944a-aff7c7759987 | 71 |
| OV | TCGA-04-1331 | d7ce51ee-6050-45ea-ad60-5209ad11625a | d7b9a611-8068-467c-a092-ce952a64effc | 7,909 |
| OV | TCGA-04-1347 | bbcc4096-ca91-49bb-b631-a4db1ca7ab8b | 545030a3-4183-4f19-820b-c9b91d6ab6cc | 4,953 |
| OV | TCGA-04-1514 | 07a37a5e-5fa5-415b-9814-363bf0a7bc48 | 0ac0cfa4-e0e9-4251-b4fa-7d81e87dc0c1 | 3,128 |
| OV | TCGA-04-1542 | b959ed10-2767-48bb-8031-f7a331c04f91 | 63e77be3-ddf8-466a-983f-89668f506815 | 4,676 |
| OV | TCGA-09-1666 | 6d2f46e9-b19b-431a-aa4e-c409eb1787dc | 5391ca9b-f632-4854-bb32-74517979563e | 4,212 |
| OV | TCGA-09-2045 | 51ef343e-2d9c-48c4-9a6f-6c4f0dcac50d | bb45bc7f-13ad-4447-bc68-fd56481f2946 | 4,438 |
| OV | TCGA-10-0934 | 91697160-9785-41b6-a697-589c82cd7017 | 789d93f9-93ef-41b8-97ac-639d85c95ffb | 3,023 |
| OV | TCGA-10-0937 | ba32f721-1816-4462-b3f7-f496ee3d8c84 | f22e0e1b-84d5-4710-9bbf-647a1f02499e | 4,548 |
| OV | TCGA-10-0938 | 6df8b7aa-5a5e-4a7c-bd82-de0c3fff4dda | 4da78c52-e98e-47d7-86ac-b9a26b492ed7 | 4,162 |
| OV | TCGA-13-0725 | f5664f55-2c18-495b-8653-d54bdd67b6d5 | b45a893b-0bf5-41d7-a0f5-79d4995950e4 | 2,944 |
| OV | TCGA-13-0727 | e58e09a0-fb52-490e-baf9-19502ff4232d | 53740d76-9b94-4e51-8980-c22f626c8d0e | 2,664 |
| OV | TCGA-13-0890 | d6c010a0-4b16-4942-963e-c8f13dbb679e | 0c89033c-66bc-491a-b285-5528f579bcd6 | 2,301 |
| OV | TCGA-13-0906 | aee22c2b-6ab3-4982-b343-b10b89c4b98b | a700cfa3-bfbc-488c-8e17-1cbe431267b6 | 8,363 |
| OV | TCGA-13-1411 | 86bafdd5-8170-46b8-a813-f7fa65336119 | 1336f459-a60d-439b-84de-d5e072a99063 | 969 |
| OV | TCGA-13-1477 | 20a1de59-1aa6-4045-a699-63a91e441931 | bc631924-38a7-4613-83f4-0a467fa94b45 | 1,307 |
| OV | TCGA-13-1487 | 3ce95963-7650-4fe7-9cbc-554ee9d56aa7 | 608f88c8-b235-4686-a27f-8917130ace91 | 3,748 |
| OV | TCGA-23-1110 | 1a385700-67e4-4e15-95ac-102406f8fe18 | 8483e431-b465-4a82-9fae-5c7d7cfbfddc | 8,937 |
| OV | TCGA-23-1118 | be8b86e7-e6da-45ca-a286-471e3ef37950 | 411c7429-f7ea-439a-8768-7a2d1b1d7860 | 4,471 |
| OV | TCGA-24-0982 | 742e3313-2df4-4757-af69-eb1b20503155 | 87fdbcaf-d814-492c-be5e-e35cb03f12f9 | 1,845 |
| OV | TCGA-24-1103 | eaeb6b7d-1e14-46ed-95d2-75c5bbee93af | 5b4fc80b-0926-4222-b2be-42cbfd68b35f | 5,613 |
| OV | TCGA-24-1419 | 915b863a-f427-4e28-8809-4e9f244f9f4d | 0db83578-1dcc-40c4-8934-76d2201eea69 | 4,346 |
| OV | TCGA-24-1544 | d293d826-e945-45a9-8e15-cc375fcf4063 | 64788c72-67a2-4602-91be-abf114e49bd2 | 3,950 |
| OV | TCGA-24-1548 | fdbe2c95-c130-4b34-b97f-39aa05a1f35e | 0a967210-59cb-4e43-9d66-c3d5b0a049a4 | 4,886 |
| OV | TCGA-24-1552 | 5549fe8c-2ced-4cb8-826b-b212f6ba24d5 | bd42ac3a-fbfc-4ba1-a233-4d0d48544f4a | 2,974 |
| OV | TCGA-24-1557 | 07bba28c-8d1a-4228-bd9e-1b27066ba518 | c40714fc-62ff-4b73-a45f-b7d160a98b30 | 5,342 |
| OV | TCGA-24-1558 | ff5c5c7d-21a4-4a46-b611-efb7750a4316 | 40db651c-2e1d-469d-8e29-a8f1ef202e04 | 3,038 |
| OV | TCGA-24-1562 | b0fd1893-f409-4444-b014-3667fdc22720 | 7ea8ecc1-0905-408c-a2b7-3d442b28a95e | 2,719 |
| OV | TCGA-24-2024 | 03e36495-9e80-442c-bc07-456ee80a47ca | a181d053-2e80-4998-bcd6-b49bc17f8a77 | 8,555 |
| OV | TCGA-24-2290 | f1ffb755-85ec-4449-b604-8e9cd194d527 | 21beb4a5-39c2-40b8-9675-288c44901592 | 5,405 |
| OV | TCGA-25-1319 | e5546364-fc34-41ba-b682-67579744998e | dbb6d3a4-35c0-402b-bbe0-a0d42c1f6b53 | 3,950 |
| OV | TCGA-25-1632 | 541a0820-9b52-430b-9fca-5eaaeb06eb4a | 106ba200-8717-449b-9661-bdec1871a250 | 4,776 |
| OV | TCGA-25-1634 | ed70494d-da6d-45b7-ad0f-59cd3afa7d46 | 707be4bb-8cd1-45b4-961a-f6cb9e120736 | 3,224 |
| OV | TCGA-25-2391 | 4c0ed657-169e-4ee1-b8b9-4544a43ed2ee | fbeaf042-3331-49be-9336-ac194218ed9a | 5,272 |
| OV | TCGA-25-2400 | 8cee9d85-d13a-49d0-a53e-c44c00ed4e36 | 8985a7c0-08ee-4d1d-92fc-eedf204add84 | 3,045 |
| OV | TCGA-36-1570 | 7bd2e9c4-4e02-4a02-83d2-c1da3a047b3a | 970cda70-6d4c-4427-a50d-bb1a5ee7019b | 5,430 |
| OV | TCGA-36-1571 | 9c491361-7721-46fe-ae77-f268796b5b9a | df2be9c4-08ff-400f-98fe-4c742b4c9f59 | 1,489 |
| OV | TCGA-36-1574 | e3ba53fc-a70c-4bd4-8dc4-60969af737fa | ccbb810b-2b8b-4c12-8bd7-58ca5fd56b62 | 4,631 |
| OV | TCGA-61-2000 | 19381aa5-278b-49eb-87b8-c53fac7ee2e4 | 5a752b74-9068-4290-96f5-28cef6542078 | 3,361 |
| HNSC | TCGA-BA-4074 | da78e5e3-ea94-4ac2-8b21-69809cabb0c7 | 669ebf5f-7222-410b-b76e-2551f11e014f | 387 |
| HNSC | TCGA-BA-4075 | 85bcc36c-7376-4108-ac13-c495a75d3e00 | 5481b0b3-1652-4735-9746-fbf7bfb454a9 | 113 |
| HNSC | TCGA-BA-4076 | 62857418-5247-4ae1-8064-d900ef856428 | 57de9078-5a6a-4185-846e-9c81add66025 | 1,198 |
| HNSC | TCGA-BA-4077 | 4f243210-88ea-4d9f-88fd-faf45b45d38e | 82ff50a2-3252-4ce5-a0ad-979cce2ec4cb | 5,144 |
| HNSC | TCGA-BA-4078 | cf3b040a-b4be-41ce-a172-1a295ebd79b2 | 406d17b0-5421-49f4-a66e-227f2076da6f | 1,486 |
| HNSC | TCGA-BA-5153 | 3bd56918-8f2b-4486-86ea-e479c2cab413 | 246bc4c7-cd66-439e-b78e-15222d1efba7 | 171 |
| HNSC | TCGA-BA-5555 | cd7725ec-bc8e-4c93-aba5-19dbdeb78b0d | cb082daa-fb68-4836-bd36-abeed40b581c | 723 |
| HNSC | TCGA-BA-5556 | e855f092-899d-469a-981c-5094b07c31ac | a2799f35-a768-4cac-a6e3-80c77052b7f9 | 598 |
| HNSC | TCGA-BA-5557 | bb531fb4-064f-47dd-8231-bd6131866968 | 11ddaf7c-cee4-47b6-a167-60967da1bd66 | 387 |
| HNSC | TCGA-BA-5558 | 4116c10d-23f4-49f2-bd5a-9b9466f08bfb | da16b0c3-9934-4caa-b4ca-5e4c5d9a4494 | 606 |
| HNSC | TCGA-BA-5559 | a6749339-8b87-4219-84dd-8045857cbe79 | e4331399-eb06-4ee0-a375-64ffa929d8e7 | 1,043 |
| HNSC | TCGA-BA-6869 | e35c380b-a1ee-47e3-b382-139ed8640d98 | 609c293a-3273-47e0-b4d1-9fd98e1163db | 63,274 |
| HNSC | TCGA-BA-6873 | 4c500ced-c371-4cc2-a3ed-72eecede6612 | 462b2014-7710-44e6-a8da-32e584676737 | 1,961 |
| HNSC | TCGA-BA-A4IH | dc2ac994-c455-4e1f-9319-2865614853ba | e4a8ada8-a509-41eb-87a1-c2bc06960125 | 2,983 |
| HNSC | TCGA-BB-4223 | e8bc6546-004a-4702-9289-b5c90582298e | a8c97047-ef56-4439-be7b-19e789bde2bf | 518 |
| HNSC | TCGA-BB-4224 | 44a7ef3f-57df-49e2-80b9-1f0dca51067f | 5fadfb17-7c88-4357-a519-b6c61ea4dc49 | 382 |
| HNSC | TCGA-CN-4725 | 7ddc2537-f164-4a40-befe-6e0a3deed186 | bf165379-dbd3-45a0-b736-21fb5757d11a | 253 |
| HNSC | TCGA-CN-4727 | f26c4909-b285-4ca0-ac40-fe65ad4a6a1a | 03edf3dd-9b2f-4ef6-8846-5a0154eacc1e | 710 |
| HNSC | TCGA-CN-4729 | 8b48a937-18eb-4e54-95d1-680ce9284b60 | a038764a-307f-4a6f-8fd0-5ad2d2e2c05e | 336 |
| HNSC | TCGA-CN-4730 | 56942e06-fb67-4bf4-993e-16cb67f52239 | 975d6473-5677-45fd-8930-d94f3addfc30 | 533 |
| HNSC | TCGA-CN-4731 | 50e66965-d2d5-49dd-a113-8b1751d6f7b7 | bb50f6f4-9a32-49a6-a81f-dca3c16699cd | 542 |
| HNSC | TCGA-CN-4733 | c1887d1c-970a-4e8d-80df-a16844daef9f | 42c2db46-32f6-476f-8c0a-eef25c7b64db | 1,251 |
| HNSC | TCGA-CN-4734 | 4b3be461-622d-492e-8465-1d0d1167701c | dc55020c-4480-41b1-ad6f-0295abea2b6c | 409 |
| HNSC | TCGA-CN-4735 | b8913a67-9dd4-41f9-8888-c998df0a0a23 | 430a1e6e-314d-47f4-82fe-d5976cdc409d | 105 |
| HNSC | TCGA-CN-4736 | 45c5cea2-56a4-43af-9c53-19302867f2d9 | c96f69ef-daa9-46cf-b34f-d7c328956edd | 330 |
| HNSC | TCGA-CN-4737 | db7d0f46-b4eb-4cad-8324-08d0a5781624 | fdf0a7f2-3007-49fa-b8f8-88f75e9f9396 | 234 |
| HNSC | TCGA-CN-4738 | 142fc981-c09b-4d94-96d3-b0d48b4d59f2 | 6420a19a-13dc-4a15-9792-d6c02672c4c8 | 448 |
| HNSC | TCGA-CN-4739 | 796943be-2aa7-419f-801f-22c78459cc72 | 1f5e9479-345d-4dc7-a9c5-24dbf2e32405 | 1,071 |
| HNSC | TCGA-CN-4740 | c1580763-a5d9-4753-b80c-f8d5498a7494 | d36d8c79-6394-457e-a92f-c582f98097a0 | 309 |
| HNSC | TCGA-CN-4741 | cd19ee2f-be9d-446a-9045-e7bfcafb49aa | ada75f19-f08c-4bbe-a617-4f070b55e658 | 323 |
| HNSC | TCGA-CN-4742 | e655fe0a-a859-4967-b69c-4c8d36a8a7e0 | 2f928254-a183-425a-b4d0-4a9317c432fd | 472 |
| HNSC | TCGA-CN-5355 | 083daa5f-0e79-4402-94b2-48a892a5a4f1 | b41ec820-7463-4bef-84a1-09d5c40fc499 | 407 |
| HNSC | TCGA-CN-5356 | c8e59618-fd16-4e8e-abe2-e3d5d674390c | 057502ad-201c-4e2a-967a-1ed9dbf9245b | 803 |
| HNSC | TCGA-CN-5360 | 56414402-7420-42b7-a93d-3d1f88308155 | e5e36209-eb45-4a7f-b70a-547a36daace6 | 847 |
| HNSC | TCGA-CN-5361 | 490cddf8-816a-45f5-acbf-3fd898eb3479 | 34ee605a-e704-413f-91ff-81dbaef22fb7 | 386 |
| HNSC | TCGA-CN-5363 | 624b1e50-b5bd-4a3f-98fd-4514bff3244d | 97348237-93c4-44d0-9bb6-3c2ee7ac10a1 | 719 |
| HNSC | TCGA-CN-5364 | c408c80c-a712-46f0-8945-22511233d2bf | 2dc9d158-11f1-4d04-a6e6-8075121def2e | 566 |
| HNSC | TCGA-CN-5365 | 84352351-1bc3-4c62-93be-29c899ff9811 | 65cf1dc0-039b-4fc7-85e5-efe031070b06 | 17 |
| HNSC | TCGA-CN-5366 | 0a7fb0ab-3302-4187-a567-58bc19979dc5 | 8d6d4c85-46e4-449c-9bc1-f75a4af110a6 | 515 |
| HNSC | TCGA-CN-5367 | 9ab4b634-5191-4a97-b9df-f75528b90bae | b73d2b70-fbf3-440b-bf21-7a3ea5ebcdbd | 332 |
| HNSC | TCGA-CN-5369 | bd85b141-d240-412f-9b74-379c4e3797d2 | 054b3120-d8ab-44a6-93c2-a8a5ae1f7855 | 349 |
| HNSC | TCGA-CN-5373 | 25d55985-95b6-4558-b42e-5c32684d89b0 | 166276f6-82c3-4536-bce5-6e520be8554c | 420 |
| HNSC | TCGA-CN-5374 | 3713f8c9-23d7-490a-bec2-99cb7ab55eac | ed899124-7dec-45d3-a9f2-79402efa65bb | 2,266 |
| HNSC | TCGA-CN-6011 | 33680362-5ef1-477f-9dbc-64dc2ed0d5a8 | 76964677-277a-41bf-8cd5-1d52cbc10e60 | 23,266 |
| HNSC | TCGA-CN-6013 | 2ca5ef3f-76f6-4db9-85e7-d32249cd642f | 6b051d97-36f0-4a18-80f9-60c27c73c081 | 410 |
| HNSC | TCGA-CN-6017 | e0dae8a5-f74f-4e25-98c2-238e28c54863 | 553eeabf-a561-46bd-8f08-ad29986c1766 | 381 |
| HNSC | TCGA-CN-6018 | ce9ca233-92c4-44b8-b97f-b56f1f73cb5a | c73af9fc-fb38-4307-955a-8e275bdab29f | 367 |
| HNSC | TCGA-CN-6989 | 190e57a0-694b-46ca-a944-0bf0e9d732c2 | a59dd8f5-b54d-4460-ac1e-f3f5831ec920 | 2,372 |
| HNSC | TCGA-CN-6994 | 165171a0-a11e-4c82-b718-897ca9741478 | 48292910-3d60-4328-adc3-59a101b81ce8 | 5,179 |
| HNSC | TCGA-CN-6996 | 533cea76-ea79-4b37-8e72-5626bc894990 | ca0120d2-68ba-4605-a647-eb86ed714611 | 870 |
| HNSC | TCGA-CN-6998 | d281a462-58dd-4303-9419-617fec3fcf35 | 3bcec620-bcec-4025-afe3-25731e0db2a4 | 1,033 |
| HNSC | TCGA-CQ-5325 | a7440672-d797-4ad6-979a-c0835f0c2768 | 3fe557f8-8ca8-4e87-b083-a50b0c942824 | 222 |
| HNSC | TCGA-CQ-5327 | 3482a15d-acdc-4fcf-86a5-b015e57faefd | f13990a7-88ec-44b6-b54c-82cd223ab0e2 | 201 |
| HNSC | TCGA-CQ-5329 | b54a3eac-9c58-453d-9db1-395601ae82ea | 9f0c3cb3-500d-46d1-8abc-1b4492354050 | 232 |
| HNSC | TCGA-CQ-5330 | a37f15ec-e0df-4bcd-8275-88312d0a1566 | 583ffff7-e647-470a-ac1d-3d6bab206e8a | 228 |
| HNSC | TCGA-CQ-6219 | e1a5a3ec-64d9-4348-8104-4f08021e914e | 24f2b999-b137-4cd9-8d46-948ef995485d | 687 |
| HNSC | TCGA-CQ-6220 | 34bb201c-76f4-432d-b2d3-a46d882b205d | 7a4c76f2-7fac-4e3d-8651-0c74feb12abb | 134 |
| HNSC | TCGA-CQ-6221 | 16de682b-970e-4857-821f-5a32d6b74f71 | 74dfb1a3-cdd5-4360-923f-2c18cd7e6044 | 911 |
| HNSC | TCGA-CQ-6223 | d12d744d-5a58-4e30-bcb7-b8c033a8255d | e392e96d-65fa-489f-829d-9a222f0ff650 | 1,206 |
| HNSC | TCGA-CQ-6224 | e8b32a2c-944e-4b3e-a302-ffe9e9f8a3ea | 532aa1c3-e7a0-4117-9718-2accf1ab0fd5 | 1,028 |
| HNSC | TCGA-CQ-6225 | a96392ab-a0d3-4c22-a2e2-08b58bb3e61b | 82431af8-972a-4db7-b424-bac44c375bc2 | 12,718 |
| HNSC | TCGA-CQ-6228 | c8e6d97c-8cc3-45f6-99a2-d20b8bc3b2d6 | 5d6324c4-f5bf-4a4f-bc7b-c12e4c489709 | 9,077 |
| HNSC | TCGA-CQ-6229 | c3f7b43b-25d2-4df3-a85c-231a6708bf8c | 16ed6dc6-b3ef-4721-a79d-db228afd20c2 | 758 |
| HNSC | TCGA-CQ-7065 | a7ab0507-8b7a-45aa-9e07-21bfc50fddd0 | 2b184042-0329-4d3e-866c-14da9baa50d2 | 787 |
| HNSC | TCGA-CQ-7068 | 77b40e7d-41a7-4e5d-a3e9-ce1e2d077c98 | ab245773-e618-4653-9bbd-066158ce2131 | 701 |
| HNSC | TCGA-CR-5249 | cef48339-6b24-4da9-b25d-55a64c71d0dc | 6a1956ba-8f1c-4504-86e4-1395788b83ff | 790 |
| HNSC | TCGA-CR-5250 | 486d7653-48d6-47cc-b04b-362d1b0e485b | ff59a531-3bb9-4afc-ad7c-dfd839bcfa0e | 2,566 |
| HNSC | TCGA-CR-6470 | 1a71c849-3daa-48f9-b309-4c340a01def2 | 08705672-13bb-497c-8d31-11c561bfe1a4 | 945 |
| HNSC | TCGA-CR-6472 | 8bf4b442-c0d0-4794-b87d-990df38dfce7 | d2f9239e-7f1c-4f81-9d85-3690f7dd3412 | 33,998 |
| HNSC | TCGA-CR-6480 | cf0a07e0-8d0a-4c16-8d48-7efb9bf38f8a | 745de1a1-46c6-45d4-b70a-c06d2a9cef3e | 5,232 |
| HNSC | TCGA-CR-6482 | 47cd6984-3399-4fc7-a843-09a97b33bf4b | 313a6b3c-974e-4fd2-8f12-6cddbb546254 | 662 |
| HNSC | TCGA-CR-6487 | 5efcbe11-0d48-4b66-abce-d78199fe8fd8 | 8874033e-39fc-438d-96cb-baabcb41a1f7 | 1,596 |
| HNSC | TCGA-CR-6491 | 85a87fce-1144-4d78-8495-597c211a5312 | 1ff34fd0-0329-461b-a74f-409c6590c6a7 | 10,359 |
| HNSC | TCGA-CR-7382 | ab5f1c0e-d07d-49b4-9fc2-b74f3c4928a6 | eac78b10-617e-41af-8cf9-dd55919bc5c7 | 393 |
| HNSC | TCGA-CR-7385 | 0d321573-5651-4eb3-9f8c-a11f802e5e92 | c5913c28-c0b8-4239-a121-017446bde4c3 | 681 |
| HNSC | TCGA-CR-7391 | ef4db578-e4b8-4047-b761-df26ecf7a41e | 1013b4f5-cc60-41aa-9ed4-32c66ca645b8 | 148 |
| HNSC | TCGA-CR-7404 | f8742c2f-2e32-467e-9347-f0dd0959da6b | ee383040-a4f3-4618-8286-715b2d1fa381 | 1,586 |
| HNSC | TCGA-CV-5431 | 0cba1ad9-9899-40b0-9fa7-dc39030c04e7 | d7c39448-b5a1-49bb-8ff7-249509a8ffaf | 2,164 |
| HNSC | TCGA-CV-5432 | 6d564719-34a3-4b79-bf57-a5c8f73d7436 | 39f4a08d-69ba-4e67-b8a9-4983f17e14f6 | 31,844 |
| HNSC | TCGA-CV-5439 | e10c02de-cabd-4311-baab-3bb90e13717f | f81e667a-9c30-4093-8e59-c8fa7f8b772c | 711 |
| HNSC | TCGA-CV-5442 | 53f0ce53-95fe-4d6c-930d-f8427c0c46ec | 40685f5a-d0e0-4f57-be78-25032bb1a9f2 | 15,935 |
| HNSC | TCGA-CV-5443 | 9e76f187-2863-42f5-9540-da82b815d6c0 | f744f137-822a-4f3c-960b-aa2f4ab9ebb8 | 2,323 |
| HNSC | TCGA-CV-5966 | a774b0c6-39f6-4601-9c2a-5a81369b9f87 | 41d152b5-0e3b-4db4-a2b0-0f2a5276d6e8 | 369 |
| HNSC | TCGA-CV-5970 | e7fcc1c5-e66d-4024-adb8-45def5252f1a | 509b8904-dfc6-4184-9e6c-1248a6280d9e | 996 |
| HNSC | TCGA-CV-5971 | 03c22e39-15a9-4b8a-8db1-cf41d618b7fc | 1ceeef7f-0aec-40ec-b515-3b369cde02e2 | 735 |
| HNSC | TCGA-CV-5973 | 91b47661-f43d-4fb8-a616-e40ed72329d3 | f0e1f7a9-618e-4dfa-91db-019ab3386912 | 3,635 |
| HNSC | TCGA-CV-5976 | ef7745ae-17be-41bc-a3a5-c24eeda89075 | f5c4e8eb-c5ca-4eb2-b92d-a5a054c4a0c6 | 696 |
| HNSC | TCGA-CV-5977 | ccdab134-2f40-4e7a-8b7c-ac1269321d3e | e6770560-36ba-42ca-bdc1-8a715a864c51 | 1,426 |
| HNSC | TCGA-CV-6003 | b314312e-2a3f-45dd-a813-870344654395 | ee070c27-f11a-4e98-9b45-ccc7d38c8241 | 1,003 |
| HNSC | TCGA-CV-6433 | 0fc542ae-619e-45f5-9496-dcf6f31f4c32 | 3d294c7d-40cc-48ed-a436-c0a0b44ef8b9 | 3,511 |
| HNSC | TCGA-CV-6933 | 76d94513-c228-445e-91f7-474a75422b69 | 8d4274cf-c829-4e6a-abf1-e3380c63ab8f | 1,530 |
| HNSC | TCGA-CV-6938 | a328ffc7-c5cd-4726-a75a-e625d33859a8 | f15c68cb-d7b4-4f77-9154-f6beb21ff1f1 | 30 |
| HNSC | TCGA-CV-6939 | 28dc7c63-6d80-4648-a356-07e8f846bc80 | c0371496-b9f6-4925-bbcf-e8a5de80d00d | 283 |
| HNSC | TCGA-CV-6945 | 50a2b9f7-d572-47c4-bf4b-3d6255512094 | 881094bd-54eb-4839-a62c-4abb208130cb | 1,357 |
| HNSC | TCGA-CV-6948 | 87c6cf32-e113-471b-86b3-cca360f47616 | 44d87ab1-c19b-4411-8774-2f888600c978 | 3,539 |
| HNSC | TCGA-CV-6951 | ec458668-da73-4d3e-8ac6-08a0e974b879 | a087903d-fb41-4d87-9500-6b62de815139 | 2,298 |
| HNSC | TCGA-CV-6952 | 7145e1a2-e609-45d9-a111-efe59656286d | 48cd5e34-4d2c-4cb0-802c-1b4c2e7a44e1 | 23 |
| HNSC | TCGA-CV-6954 | 115469a6-3c3e-4a3b-adf9-ed831266829c | fa689d79-bedd-452e-935f-21bd4866b9d0 | 51 |
| HNSC | TCGA-CV-6956 | f9762350-aee5-49d2-a5de-165317ff2bd3 | c03352c8-542b-4ea8-b982-ff61e0723b16 | 12,669 |
| HNSC | TCGA-CV-6959 | ff9553cc-07a7-497f-be70-1ad5b8e09130 | e12b73cf-efab-4371-8449-776e46f7949c | 566 |
| HNSC | TCGA-CV-6960 | 5fdfb781-64fd-48dd-9d26-3de7bfdf5908 | 48a5d368-878b-4e4a-9232-501b2c2dfbfe | 1,215 |
| HNSC | TCGA-CV-6961 | 4b9344a0-a7c6-453a-b184-87e94c97920a | f4233ca2-6eae-497e-8d71-234f85182d06 | 13,965 |
| HNSC | TCGA-CV-7090 | 30e23a3f-f6eb-45e4-9d14-bddafc4fab6d | ba2b472d-b185-4c74-8e0f-8ff0e82db9de | 1,199 |
| HNSC | TCGA-CV-7091 | 8a0170b3-9cda-4426-be4b-9e488f6244c2 | fd03b019-209c-4744-9f18-0fdd68024ed6 | 2,882 |
| HNSC | TCGA-CV-7095 | d944910b-a3cf-45e9-84be-5ea0e6cd6f03 | 362b9d7a-e5e5-45ad-b441-a3e549093fc5 | 1,131 |
| HNSC | TCGA-CV-7100 | 4e9d6eec-9af8-4de3-b449-1d56dd8ba30f | 1a0d4aab-84fc-453c-a92a-e4af5474a74a | 1,608 |
| HNSC | TCGA-CV-7178 | f76f830b-a361-4cf5-a60f-69d4d574723f | 428da9ec-1409-4479-843a-6d2ecff5f7b8 | 1,532 |
| HNSC | TCGA-CV-7180 | b66083b7-e0ae-4f6e-ab6a-76caf55a13bc | 7a1bc885-10c7-42e5-9024-c742bde8c9cc | 691 |
| HNSC | TCGA-CV-7183 | f449ea84-f7cf-4a26-bc43-707ac56f0038 | f7f69891-d84b-4137-92ba-59f738bbc299 | 31 |
| HNSC | TCGA-CV-7238 | 3eb627a6-bbcf-4699-ad23-e52a26cd4453 | 70f8bf18-8050-4dad-9029-c90972a9bfb4 | 718 |
| HNSC | TCGA-CV-7247 | 0b4ae8f3-7acd-4d37-a39f-cb4fc2b537b4 | eda6d221-38ba-4187-8a10-2c0217d09e31 | 573 |
| HNSC | TCGA-CV-7255 | 7f307e2b-3164-4bb0-aa04-90c538b8bb46 | e24cfdc8-ec05-41ac-aa61-3339c1769152 | 2,566 |
| HNSC | TCGA-CV-7263 | d8d79518-dd53-49b5-850c-3af45043e30c | 64a6f3f5-8b60-439f-86f8-603118060103 | 1,136 |
| HNSC | TCGA-CV-7407 | e5709830-4141-4354-94f0-94f576fe84c9 | 324e0ca4-45b2-4348-bcc6-969674ada4bc | 280 |
| HNSC | TCGA-CV-7411 | 7db9b308-952d-4c12-98c4-9de1f4b9fbd2 | 9ad035d2-035c-4447-a744-24409cfeb23e | 564 |
| HNSC | TCGA-CV-7414 | d2264558-e25d-4169-9e22-8f3c23b50ad9 | 2f7acc14-13c2-4d7a-8ba8-6f6e9d6e12b6 | 884 |
| HNSC | TCGA-CV-7416 | 1374c517-8c54-4a22-80e6-0f48eb1d4de6 | 6650adf8-2533-476f-a2ae-1471135763a3 | 5,776 |
| HNSC | TCGA-CV-7428 | 234197c5-8479-49c1-9f04-5e9171364a3d | ff322aea-a4fd-4377-81b7-e8cac37aa770 | 1,032 |
| HNSC | TCGA-CV-7429 | 4860e1b8-251e-4894-91f2-c9416a9b7f28 | 6998b65f-568a-4636-a6cb-07fb23e984a7 | 691 |
| HNSC | TCGA-CV-7432 | d1f8d81a-4553-4c26-9d59-534aa2475dca | 7698153d-ef10-4787-a1d6-4b0022b02236 | 16,001 |
| HNSC | TCGA-CV-7434 | c645c62d-79d3-478c-a9c4-16294daad71c | 61309435-9e3d-4fc8-989e-63773d630677 | 590 |
| HNSC | TCGA-CX-7085 | 2a118e3e-c8f9-43b4-b1d7-4659e37a96ce | 2187824f-bc90-441a-8180-5f1524daf954 | 164 |
| HNSC | TCGA-CX-7086 | 2c64cb1b-6ae4-4191-9b2c-e5c13e267e83 | 62f53214-e1e5-42fe-821b-a2a656206e1f | 11,656 |
| HNSC | TCGA-D6-6515 | 5a79073a-b6a0-41df-b26b-247649784d4d | 9162c3e1-cccb-4dbd-8584-1ed07c67a6ae | 668 |
| HNSC | TCGA-D6-6517 | e84f34a5-8df7-4824-92d7-4f53b4bace58 | 019b158f-ce54-4f1b-9db2-b82383657121 | 2,027 |
| HNSC | TCGA-D6-6823 | 250e5855-f3af-4c16-9efe-918be02bf754 | 60900e52-222a-400c-9981-8ea2dfba4553 | 608 |
| HNSC | TCGA-D6-6825 | e785062a-dcec-4a1d-aa67-6089980c9e35 | dac94d31-7c1f-4e4b-98af-a3a585b86fae | 368 |
| HNSC | TCGA-D6-6826 | 377f848a-3e59-4eee-bd9d-87fdc1a898bb | cb6e32f1-cfa4-4387-a53f-e7a562ac45d9 | 440 |
| HNSC | TCGA-D6-6827 | 849f41d7-0557-40ce-bca0-8a54c496fef2 | aef80235-e93d-436c-8a32-888344249391 | 468 |
| HNSC | TCGA-DQ-5624 | 72ea153a-e312-4f8e-8fa6-bd705df2b106 | 72bf54c2-0056-4578-8448-34fb30ed8b3a | 2,596 |
| HNSC | TCGA-DQ-5625 | 33b84364-a81e-48fd-89d6-552802c7ff18 | faef45e3-26d8-4053-a1ff-b62774815392 | 1,205 |
| HNSC | TCGA-DQ-5629 | 3fb0505f-e847-4658-93f1-0af7d8617e66 | a61bf907-f580-4de5-8210-8c01ede3d802 | 37,946 |
| HNSC | TCGA-DQ-5630 | 865b23ff-32b6-41bf-8316-cde4e80d7873 | 1c95b304-0beb-462d-bb72-fe1946dfbc5f | 934 |
| HNSC | TCGA-DQ-5631 | 9aae2925-69ec-44f9-bc1d-bfefd16ce242 | f614c1a4-6710-4035-bf2b-23679a3c3fd0 | 996 |
| HNSC | TCGA-H7-7774 | ab347312-e135-4dd6-9be3-fa5fdebbcd68 | 4b8c5d66-16c6-43b0-8204-3a12b6a6394b | 750 |
| HNSC | TCGA-HD-7753 | 393b4af8-b44d-4c3f-b5ca-f7032b687a16 | ddb75587-4d2e-4c11-bd83-f7a0accb8120 | 7,898 |
| HNSC | TCGA-IQ-7632 | 9f8feb9c-6c20-4997-afcf-24177e9a7829 | dd7b51dc-f3ec-450a-91b7-909ceb81e85e | 717 |
| LAML | TCGA-AB-2963 | bede1739-c266-4e53-b85b-b8c54790c7ae | 19dbf21d-36b6-45f2-98d8-29add148f771 | 1,214 |
| LAML | TCGA-AB-2964 | 1eda9412-6d2e-4dc7-8513-b54b7bfa14a9 | e631649e-21df-440a-85ae-a99a41de267b | 4,116 |
| LAML | TCGA-AB-2965 | 95ca4aa2-cfaa-401e-aeb9-17abba0d8f70 | 1748acb6-a5e7-430f-b6a1-4600479c5eee | 1,194 |
| LAML | TCGA-AB-2966 | 4301b67a-475e-4317-97a5-694805d11c3e | a4c40a4e-68b0-4057-a22a-61796302a625 | 10,178 |
| LAML | TCGA-AB-2967 | 468b2662-9228-4abc-a54e-b3f919009c73 | 72716dcb-583b-4c55-87af-a6d8b4ae9244 | 1,578 |
| LAML | TCGA-AB-2969 | d991ead7-f6ea-4bb1-aee6-4bc4ee04c507 | 621bb448-1878-421c-84a0-6d52ee837fcd | 2,034 |
| LAML | TCGA-AB-2970 | 64bc5114-6e38-4dc7-b54b-47dae02ba515 | 9732df1c-1297-4584-ab33-65ef514d5452 | 887 |
| LAML | TCGA-AB-2971 | f780fa08-5d0d-4c0b-92f3-7052b52807b2 | c85319e4-b2e2-4551-b55a-550cbb8679f5 | 1,184 |
| LAML | TCGA-AB-2972 | 2b0ac0aa-7f5b-4f7c-8e77-36e6b2c7ceb0 | 864f579d-951b-4a07-bf55-de72e9a84188 | 1,820 |
| LAML | TCGA-AB-2973 | 12cbb28f-634c-4a40-aa29-c68f5562d67c | e3ee1732-fa04-45e6-83d5-8cba7f3094fa | 637 |
| LAML | TCGA-AB-2975 | d504ac62-ddde-4e0b-b070-88b97480837e | a121ea24-8e5c-49a5-8bc0-4110f8a6efc5 | 454 |
| LAML | TCGA-AB-2976 | df134dc1-7033-4c17-8ed5-eda72828667b | 550deec0-7595-4da3-87fe-088ebb3e4791 | 1,623 |
| LAML | TCGA-AB-2977 | 2120160a-fc45-46c9-a64f-bb9967ec366a | 2123339d-a200-450f-a7ee-80ce07c9b809 | 2,404 |
| LAML | TCGA-AB-2978 | 90ed4aa6-62ac-4580-937c-3438ee02ff46 | a1db1f04-cd5d-434a-9b0a-3e06016cc569 | 1,532 |
| LAML | TCGA-AB-2979 | e491dbf1-cc30-4493-bd32-4f6c09f6d468 | 9cfe41a4-e968-444e-874c-95185e9b1be7 | 925 |
| LAML | TCGA-AB-2980 | ec05a71f-c3c8-475c-8a2d-a858dfd2a291 | 03fcf98d-69fc-4abf-a465-8c834383c1f8 | 1,211 |
| LAML | TCGA-AB-2981 | 2f164e15-1d90-4d0b-bf80-f15a9a3f119f | 028aa18d-e3c2-4357-9582-bf225ff59aa5 | 950 |
| LAML | TCGA-AB-2982 | 04c340c1-a92e-42f7-9127-ca462ef38d86 | af79274a-6f4b-45af-b3ae-02f3780dcfc0 | 1,133 |
| LAML | TCGA-AB-2983 | 059a4661-5fc8-4e14-9e9d-208ab8e28688 | 9cfc4eb7-3c47-4ac3-bea9-869e65cdd0f0 | 1,898 |
| LAML | TCGA-AB-2984 | f24533b7-61fb-4028-b18c-c923e9e45b57 | c039cd89-dab1-4867-bb10-2a5957d4a71b | 1,829 |
| LAML | TCGA-AB-2985 | 17d2f07e-10c4-460c-a21c-43c5bf149927 | 3dc6b3cb-d03f-463b-bad9-c14d148ba6df | 595 |
| LAML | TCGA-AB-2986 | 611299d4-8cc7-4a90-8ee5-0ebb04f3b5e3 | 0899070c-b0c0-4d7a-8c0d-877f32a72142 | 521 |
| LAML | TCGA-AB-2987 | 5efd36b6-4b88-4c0d-9ab0-078daf12d67f | f137b969-aa9b-4f6a-ba66-2c9b21c41562 | 786 |
| LAML | TCGA-AB-2988 | 7e3f8199-9ea9-43e6-b392-b26841eaae42 | 4beb8dec-0fd9-437e-ae5d-50f8e5cfb7c2 | 1,725 |
| LAML | TCGA-AB-2990 | 3d27c41f-ca77-42a7-8b56-d5f6fcdc52b9 | 091d42b4-4a69-4f84-bb1a-ebf5a75ad283 | 954 |
| LAML | TCGA-AB-2991 | 49f91ddc-95d8-4158-9c18-1d3b432289f8 | 080ebb87-5b7a-4842-be66-17a04dfb9ca7 | 1,111 |
| LAML | TCGA-AB-2992 | 1ff3ab37-b951-4cde-996f-2fb38724fba8 | 25f893a0-7c82-4111-bc7a-43819e613133 | 6,909 |
| LAML | TCGA-AB-2994 | cdc58906-e1f2-4eb6-bf55-3450c8613244 | 66f2cb92-7986-4225-9703-acb61f0c1bc0 | 712 |
| LAML | TCGA-AB-2995 | fffb1142-1235-425f-92ed-bb291a36d971 | d6e0d24e-dbc6-46de-86f5-d07186dc3dad | 581 |
| LAML | TCGA-AB-2996 | ac3e4260-9459-4295-8d4f-ec11b112caf6 | 1fa7cc53-2c51-42a9-adae-e1bfc94bea98 | 1,537 |
| LAML | TCGA-AB-2998 | 85602a35-6c04-4309-8b76-c0ababd8040f | d251898a-fb20-4a77-bcc6-4b0f0a3c588e | 1,482 |
| LAML | TCGA-AB-2999 | e80221de-1dab-4ff7-b8f6-9631f09eabed | 20f5e9aa-90cf-4edb-8dbd-efd0ea4e0a91 | 2,647 |
| LAML | TCGA-AB-3001 | 810acc51-00f3-42f9-a53d-bac39e3e668b | 88361040-7c46-4509-84d2-9b9f711a3d36 | 2,108 |
| LAML | TCGA-AB-3002 | 0379d38a-fc2c-4a6e-a324-da5ec8552776 | cd0f3104-6cba-4878-ade4-6b2596c38496 | 2,215 |
| LAML | TCGA-AB-3005 | 62eeae43-d36d-4f1a-92a8-8a62f53bfd71 | e5b40042-4030-473d-bd3a-f78f8155447a | 1,411 |
| LAML | TCGA-AB-3006 | 3f83b9a4-47d8-4c54-a44f-4c4515936eb5 | 133e5abf-67bb-4e4c-be46-780705a4ca5f | 1,702 |
| LAML | TCGA-AB-3007 | e1a51d5e-e946-4e89-96a3-26c7881fd5e5 | 6bdd14ea-2d9a-41d2-9823-4fbce3e774d0 | 1,571 |
| LAML | TCGA-AB-3008 | 19f482df-3ecd-45bf-81e0-dd19cf432ada | 72e1624a-d8fd-4f9f-93df-b723298329b2 | 1,019 |
| LAML | TCGA-AB-3009 | db39d6c7-345c-462f-ae6f-fba90d4eefe5 | 38684811-17f7-497e-af49-ba81330afa07 | 3,980 |
| LAML | TCGA-AB-3011 | beceffbc-24e5-4c31-853f-66b75d98889b | 94f9b703-99a4-4113-9ef3-dc03f468025e | 748 |
| LAML | TCGA-AB-3012 | ba4506ad-207a-4732-b1fd-89e62a16aa6b | b9a0ff19-c21c-43ca-b213-3c263b98deb7 | 1,626 |
| LGG | TCGA-CS-4938 | 8b9dc3cd-2c99-405c-8306-3b388ad5bdf6 | d987c181-1947-486c-957f-c5bd782a1eba | 61 |
| LGG | TCGA-CS-4941 | 7df56317-1eca-4fb0-af19-36a4bfc5df00 | 76207c4c-20a5-4a76-aa64-df7b90201a78 | 431 |
| LGG | TCGA-CS-4944 | c266d33e-c72f-4059-b287-2f1bbd6fd48b | 4454284c-2c7d-48f0-8ddb-ee4896355ab4 | 281 |
| LGG | TCGA-CS-5396 | 3d26b7bb-4cc9-4d9b-8c42-6fbd2ea9b590 | 02bd97c1-a586-4f04-b6e2-16ca05a0ef21 | 220 |
| LGG | TCGA-CS-6290 | dd2c0db0-5e68-43b9-a591-4efa9a782af1 | 5e2f3e2f-546e-4dde-b8c7-ab81ce38c575 | 286 |
| LGG | TCGA-CS-6665 | 9d827820-042a-412f-ae10-1f97190c446d | 63dafea0-a011-4868-ae45-ad0d1524303a | 1,999 |
| LGG | TCGA-CS-6668 | 698a92bc-1d75-4e94-a533-d41f6a195e9b | 51192faa-2766-43bd-83da-486a2f32e8c0 | 2,271 |
| LGG | TCGA-CS-6669 | 7069c071-1fef-4c6f-b741-c67ee56619bb | ea003ef8-eae8-4e85-9599-556b0369c165 | 220 |
| LGG | TCGA-DB-5273 | 844bcbca-fe48-4dee-be69-4cef2ac59e27 | fe0c497b-caa5-4108-8564-a1507ba99ed4 | 149 |
| LGG | TCGA-DB-5276 | 02206442-a052-4c44-a4b8-1467493df2eb | c584cffe-856d-48a8-b1e6-95db5cb5f3de | 246 |
| LGG | TCGA-DB-5278 | 6780f08b-966e-4836-9cd8-46b87be05578 | fa6a196d-09f3-47ff-a614-66b54fa20d3c | 391 |
| LGG | TCGA-DB-5279 | 1c15ff7e-3cbc-41b1-b814-1cf03f1f5a27 | 767a0a89-778e-4312-bd3b-123ddca4d6e2 | 164 |
| LGG | TCGA-DB-5280 | 47363648-94bb-4221-a905-4832396ef90e | c2968c6a-5afe-480e-a036-e3522ae763ec | 81 |
| LGG | TCGA-DH-5143 | 9e2bca97-d504-4863-b306-8f559a2cc8a1 | 14764abf-abd2-4bc1-a8ee-79b07761338f | 385 |
| LGG | TCGA-DH-A669 | bc4e3b89-3a13-471f-8769-7d711cfbad8e | 958ecf79-2dcd-480b-a65f-d208b59fbf57 | 3,320 |
| LGG | TCGA-DU-5847 | a76674c6-34c4-4c06-b7e4-eb602b3bc5fa | d565f96e-c548-46a5-8dd2-341da735c3d8 | 319 |
| LGG | TCGA-DU-5849 | d4f0a4e9-de7c-498f-89e7-aad71fb01915 | ce60dc42-d2f9-4eda-b08c-8e71133127b7 | 218 |
| LGG | TCGA-DU-5852 | 9dd03595-2a2f-4fc1-9cd1-52a490696f01 | 6d9e6678-64e1-4fa4-b57d-338f163a62a1 | 215 |
| LGG | TCGA-DU-5854 | f499956c-7655-4be7-8a53-44403d30ea04 | ef916508-4f9f-45d1-b65b-8d69a7895db8 | 279 |
| LGG | TCGA-DU-5870 | 074b6b1f-4846-42c9-a6dd-ea94aabfb270 | 6f6fd391-7fbf-4f78-b27d-7f29183238a3 | 1,550 |
| LGG | TCGA-DU-5872 | 83a061e8-88d1-4f4e-8568-8eda2807ad1a | 31f28101-11ef-411d-baa5-15d1a1618d2d | 1,047 |
| LGG | TCGA-DU-5874 | b0268c63-93dc-4e0b-b29b-217fd6028c92 | 8fae1d21-7d63-4949-a484-770f1751d03f | 3,222 |
| LGG | TCGA-DU-6392 | 8f4eff17-77e1-4e7d-9fc7-83ae4db50b3a | 743888e9-97f5-42de-bb8b-acf4ade0af5e | 2,280 |
| LGG | TCGA-DU-6397 | d184ed85-cb21-43cc-95f6-0619f9283b29 | 623cc431-e17c-4b40-97ed-b5178540519c | 1,340 |
| LGG | TCGA-DU-6399 | bcd63f96-cd75-4e38-826d-5304bd0d18db | f4761372-b118-4406-97da-d7c39196df5e | 209 |
| LGG | TCGA-DU-6401 | f0537710-8458-4890-8a52-3f4457eb308f | 30e7af75-7d8e-4aa5-b01e-1149dff334ac | 1,192 |
| LGG | TCGA-DU-6402 | d556f16a-ad20-42eb-8e92-57114d46e174 | 3cd68c5a-a87d-41a1-97aa-29c156e57988 | 225 |
| LGG | TCGA-DU-6403 | d17b6f47-d1c9-4916-8722-8b6908f8552a | 5d588402-a79e-45a9-ba69-9bd3eba3aa6c | 188 |
| LGG | TCGA-DU-6404 | e8c73506-ba8e-44dc-87cd-4305bdb6b7f9 | 83019f18-87de-42c9-b3af-30c0cd151fe0 | 857 |
| LGG | TCGA-DU-6405 | 2d1d4000-347f-44e4-8ff3-b61fea3daeb3 | 107b4f56-260b-41d1-85fc-a6746e87ca0e | 236 |
| LGG | TCGA-DU-6407 | 5c3c83f3-c65b-4c23-99d4-64183122fe2c | cfed9604-e0dd-47be-8aa0-026854f244e2 | 1,603 |
| LGG | TCGA-DU-6408 | f441e949-5e27-4235-a0db-39c77aacdb5c | 8157eb3b-a140-425c-af13-e727b719c9ec | 223 |
| LGG | TCGA-DU-6542 | 4b6cb752-5ed6-42e5-aed5-c8bdc6b95945 | dda7afb5-9e07-42bd-b2e0-7499a263ebb5 | 14 |
| LGG | TCGA-DU-7006 | a423d9b3-2389-45d5-b834-d5cf57e5e647 | 7d7ca01f-5584-4e12-a544-5ffb6d2cdc3f | 156 |
| LGG | TCGA-DU-7007 | 84edb90d-0a24-46dd-bed3-126c5fdcda42 | df044351-b6ec-40c5-befc-9a76d30cd84c | 194 |
| LGG | TCGA-DU-7009 | 446ab57b-076e-4f55-b617-050e5f14bfa5 | a5ee0d34-649a-4a5e-8d63-b9d2dc8196e2 | 1,258 |
| LGG | TCGA-DU-7010 | 61859b0d-b4a7-4c67-bf02-21422618f2f2 | 550d6316-2067-4f67-ba64-a8687f3320b1 | 237 |
| LGG | TCGA-DU-7018 | 2736e751-ccc3-4721-b516-238f987ad26f | 7ee7f69e-1fcd-495d-bbfa-f262e711b4a6 | 148 |
| LGG | TCGA-DU-7290 | 827baf09-e242-460a-8d97-75059c6c3a98 | 3a7ec773-52e2-4eed-bd70-2dc1f059a2ec | 153 |
| LGG | TCGA-DU-7292 | 06fe9cf9-6572-4026-9990-7ec39cd21278 | 0ac4dae2-d60a-4869-a3a6-c2ecd1c7640f | 138 |
| LGG | TCGA-DU-7294 | 2df0ab9f-8d32-40e9-883d-f5afad653272 | fb816713-22f8-4650-a51e-d1dc31150849 | 149 |
| LGG | TCGA-DU-7301 | a788486c-7bba-436f-a31c-a5d294155cc6 | 8d315f21-d285-46ed-b707-ff48d1be543d | 2,431 |
| LGG | TCGA-DU-7304 | 7f2ad00f-02aa-4094-9662-9e8092d240e1 | 3aef62ad-f4ed-44ff-aaa9-13bad668bb24 | 2,172 |
| LGG | TCGA-E1-5305 | 0f660d53-1dae-499d-8a76-f591e1110acb | 69689ca9-e4c2-440e-9e41-961f4b548891 | 86 |
| LGG | TCGA-E1-5307 | 80d2e037-eb0e-4514-bdd4-caa1fc77176d | 39a2e360-0a4c-4285-ace9-946e589970f4 | 95 |
| LGG | TCGA-E1-5318 | 0fc8cc9a-c3ea-48d6-8f7d-18ab9b69ffb1 | a572d1ce-26c6-4788-9feb-6f9000207266 | 1,885 |
| LGG | TCGA-E1-5319 | 6b14040f-a3ce-4fe5-a5f9-f311cd1ec11a | d36d48eb-6148-40c8-be7d-706f53852d54 | 1,110 |
| LGG | TCGA-EZ-7264 | e0ea4522-35f1-40c7-92d2-5d1e7ec0cb63 | b74ca950-7df4-42c9-ba4c-a7c1599fe56f | 1,696 |
| LGG | TCGA-FG-5963 | 11f814ba-95e7-4824-ab33-1e14947dcbd5 | 58b865f4-5c4e-4736-b667-a649145bb764 | 154 |
| LGG | TCGA-FG-6688 | 3b45431a-1066-4467-bc8a-f1aa3342c5f0 | ed5c4e17-1960-4f9e-9152-02db5a6f6ec9 | 95 |
| LGG | TCGA-FG-6689 | 9e07a7ab-1033-43c4-a4f4-49ba35319fea | 08b74256-79b2-4fe1-81c5-28cc122988ca | 136 |
| LGG | TCGA-FG-6690 | e33fa79f-3d33-4f0b-81d9-705ab9c8a19c | a570edf3-b9eb-4c0f-b0d3-cb958f070979 | 145 |
| LGG | TCGA-FG-6691 | b404ac67-1c7f-4b01-8038-7432d3d6e489 | 87f5566b-671c-48b4-8170-bef9aa55eb6b | 86 |
| LGG | TCGA-FG-6692 | 66eb8fac-6ead-47b2-be40-effbb245f499 | 08012a5a-0f7a-4f6e-92cf-b1878218e33c | 91 |
| LGG | TCGA-FG-7636 | 336bd48f-3c20-4a2f-a2ef-38bb94e30e11 | 3936d9e1-9fc2-4de3-9c2d-6bb41eecfcf5 | 33 |
| LGG | TCGA-FG-7643 | ad52b277-d592-4ece-a72f-4d48f9394453 | 74141a93-8537-46c9-b176-28adb5fb5f94 | 365 |
| LGG | TCGA-FG-8182 | de01f2de-931f-4b79-9abc-966eaee7ca0a | cf62b63d-2102-4de2-9286-867afe868a79 | 602 |
| LGG | TCGA-FG-A4MT | c0b259dd-e27a-46af-a639-f80a0e1e7fff | 896a3c30-42ec-4bb4-9b26-cb4ce573d73a | 2,163 |
| LGG | TCGA-FN-7833 | 91f0d100-c921-4319-9d82-98819189b525 | 8a4a2069-1e0c-4d49-a1eb-77f38954eed2 | 65 |
| LGG | TCGA-HT-7468 | e1855a2f-1904-41ee-ba33-a27e1c202d99 | dd94727e-cce1-407e-99ce-89064229b3e8 | 204 |
| LGG | TCGA-HT-7472 | 518f9b23-97da-447f-90e5-a03c5ae09642 | 480868bd-a33a-4c42-855e-c6bfafe22ecb | 132 |
| LGG | TCGA-HT-7473 | 0c3613f1-9a5e-42d1-a1a9-884cf56894f4 | 2b1e29cb-e83c-4e25-a643-1ded3bbcccb6 | 222 |
| LGG | TCGA-HT-7475 | 13942ed5-f37e-4807-b133-369aab923f17 | 688c23ec-5c4c-438c-9553-132c7c3da53b | 351 |
| LGG | TCGA-HT-7476 | 8eda6747-01ab-4378-b3c9-6f638de37412 | 79fd9f03-72a7-4dca-8389-9855dd6660ae | 171 |
| LGG | TCGA-HT-7481 | fbc2ca18-8dd6-41c3-9ec3-0189f96877ce | 7b00539d-b389-4e45-b200-b742550d68d3 | 141 |
| LGG | TCGA-HT-7601 | 5faa261d-a169-4ab5-a52b-de3b50c46299 | 94873a5a-9d98-4c3a-8ea2-528afa271e07 | 78 |
| LGG | TCGA-HT-7602 | fdc7bab5-c807-4791-be95-6a588e86a4b8 | 73cb1269-4aff-4f51-a2ac-32f3211af789 | 320 |
| LGG | TCGA-HT-7604 | b3d0b563-b12d-4872-9a0c-f795ca13842f | b43b5074-d9bd-4d06-b367-bebab284ebfe | 96 |
| LGG | TCGA-HT-7689 | 55583ba3-88d1-4c7c-8bf7-bfe2699d74da | 4e07a7fd-8425-484d-8ef7-d48bd50ff286 | 1,328 |
| LGG | TCGA-HT-7695 | 0e08feb2-0a06-458b-a3ea-797020f9a68f | bd971aa9-0dc2-4b9f-8d94-b08b15ca0339 | 749 |
| LGG | TCGA-HT-8104 | 85f4a2b5-246f-41da-9777-57d2ceaf1b70 | 29612660-50bd-442c-9c73-a18e3a340ee8 | 3,993 |
| LGG | TCGA-HT-A5R7 | c239bee2-d22d-4390-be44-ed4be0395f57 | 80186405-a532-4ebd-9122-8575deff3ec0 | 651 |
| LGG | TCGA-HT-A61B | 95493209-d334-4fbe-aeef-683bf27cdf09 | 303ebcff-0840-40c9-bba1-2f343a9883a3 | 1,677 |
| LGG | TCGA-HW-7486 | 77a6b9b7-4a25-4784-95e5-98346720dcef | 78f274f8-ffb8-467e-8f6b-7d6fd1852127 | 376 |
| LGG | TCGA-HW-7487 | efb882b9-fe40-4205-a704-5757740ee898 | 141d09e0-7b88-4ec6-acd5-3e8588633fd3 | 1,485 |
| LGG | TCGA-HW-7489 | e81e0ec5-b546-421b-a1ee-5adf17fd07c0 | bcccd3c8-2eee-44ca-bb10-16e4dc8672d5 | 190 |
| LGG | TCGA-HW-7495 | 10d85d7f-6926-453a-bfef-d1b9d129d9d1 | 938bedfd-8385-4a31-a739-3d8bb17c8a1d | 216 |
| LGG | TCGA-IK-7675 | 19191a31-9d4e-43a8-a88e-aff919fd0f95 | 41e4eaa9-9947-44d6-8638-6ed9593c9670 | 5,258 |
| LGG | TCGA-TM-A7CF | 41c3ad23-569a-452e-8577-85a783c2a020 | 96ad4e5b-aeec-4421-9d5e-120287a57086 | 1,592 |
| LGG | TCGA-TQ-A7RK | 4b48a880-f86b-4e8b-9d21-65c14ba90284 | 2e36e341-bb8b-468c-bec6-a3cac541abf0 | 2,041 |
| LGG | TCGA-TQ-A7RV | de819dc3-dbde-4a92-b859-86e1e2bcd3c4 | 95a7f273-e22c-4af9-b245-ca5d839080d2 | 1,019 |
| SKCM | TCGA-D3-A1Q1 | 6ea97d1a-9ee3-4afb-9ce1-ba48e4f7599b | ef92a606-f1f0-4fce-b864-ac45d19029b6 | 3,675 |
| SKCM | TCGA-D3-A1Q5 | fde49f52-05cf-488c-8ac4-45a538a6b006 | 55b34f20-71ee-4c4d-abba-4d2024a125c3 | 1,291 |
| SKCM | TCGA-D3-A1Q7 | f9cdbc4b-24de-4377-ac12-2d532ee5fa4f | e3e0036b-8e9b-4e50-87ac-5b77b6295bd6 | 540 |
| SKCM | TCGA-D3-A1Q8 | 0f0ce208-c694-4ce1-9864-0eef3d792bb5 | 9944e3cf-cf47-4a95-bb4f-b82804de4674 | 235 |
| SKCM | TCGA-D3-A2JC | a7d7b7f7-01dd-4c3f-9406-513601821293 | cfb3a5aa-61a3-4974-b482-8860495c3f56 | 554 |
| SKCM | TCGA-D3-A2JD | 65653c49-be73-44c2-b13c-3aa7f55d5748 | 28a73c15-1674-4322-94fc-ecf6ca914d51 | 666 |
| SKCM | TCGA-D3-A2JP | ed56a64c-9f71-4163-bdda-c3d36448943e | 4b9843ae-869e-4a07-ba84-7e300b9c03cc | 203 |
| SKCM | TCGA-D3-A3C3 | fcc6bf30-9ef1-43ac-84e6-68661b4266d8 | c29f3065-3e54-48d6-bbed-7aac0fbd2f92 | 280 |
| SKCM | TCGA-D3-A3CC | fd606b6a-a303-4b04-be1d-f12779b19969 | 59c97e2f-e76c-4017-a55f-31efc29ca6c1 | 130 |
| SKCM | TCGA-D9-A148 | 5581a0f7-47cd-4369-995b-8c74d2e3d62e | a0f07bb7-ce52-49fe-83aa-97c4fe2af227 | 246 |
| SKCM | TCGA-D9-A1JW | dd38c2b9-2249-41f4-81df-ff133c8dc39d | 81b083e1-deb3-4935-b274-bded2a4d47fb | 114 |
| SKCM | TCGA-DA-A1HW | 2ebf409b-961e-4948-ba57-3e2cbf64f8e5 | 19d8b19f-ad7f-4a2e-8a4b-e1cac0212f82 | 4,842 |
| SKCM | TCGA-DA-A1HY | 4067ff38-29aa-4cbb-98e3-14283e4a270b | cd19c1ed-65ae-412e-8d09-572a87a53517 | 3,559 |
| SKCM | TCGA-DA-A1I0 | 66bdb1db-88a0-4989-9f84-06291d75ac65 | 2bc72654-62f3-4a1b-bb0e-f9fcf6b38eb9 | 291 |
| SKCM | TCGA-DA-A1IC | bdd7c2f0-1c47-4c40-bde5-745c17b0824a | 615d39b9-8a2a-4712-87f9-40c1911e199c | 8,533 |
| SKCM | TCGA-DA-A3F3 | 7899dc05-8772-49e2-895c-202ffb731c6f | 01f1d9e1-0bec-49f2-8dec-55f3f6c3b9f6 | 622 |
| SKCM | TCGA-DA-A3F5 | a8b342c2-9bdd-4e3f-a1b0-0f1796f5f603 | ea9e834a-de7e-4891-b51f-6b0b0e4c362d | 401 |
| SKCM | TCGA-EE-A29B | 0d511b2e-752b-43a4-b27f-81a205cac15e | 628ddf51-45ee-4b67-891e-132909689e76 | 180 |
| SKCM | TCGA-EE-A29D | a8d2aa91-891f-4915-ab15-e620b21c436c | 1b436706-8b80-4fb0-bf1c-1375bfc4d0f1 | 233 |
| SKCM | TCGA-EE-A29H | 21070c2b-7503-4c5c-9a66-5e6564c37d01 | 66ec2170-64e3-497d-aa60-e0f40b3875a6 | 124 |
| SKCM | TCGA-EE-A29N | 1a8bfce8-1606-45e1-8da4-fbad852ec59c | a5f48fa3-7af5-4273-be19-0d3562ccb440 | 94 |
| SKCM | TCGA-EE-A29P | 54534e99-c9b7-483e-aa8e-c9fd9fa41058 | 74fe7aee-fe79-4d9a-bb11-86f865d6a0b2 | 133 |
| SKCM | TCGA-EE-A29Q | ac015b9b-1772-42b1-a39c-c771b610492d | 3a35cc5e-4313-46ab-857d-f22cddd7625f | 147 |
| SKCM | TCGA-EE-A29S | b6eef3eb-6a3c-4010-a637-48540f303a1c | acaa3027-4446-467e-bc84-eaf1b02d992c | 138 |
| SKCM | TCGA-EE-A29T | b9efa6bb-7ce3-4b05-9ecb-6d56bedf6e7b | 83148501-88fd-4908-bad2-144b603a893c | 911 |
| SKCM | TCGA-EE-A2A1 | d2b1ca54-da4a-4eba-8e02-a1cc07f55639 | d20d23e7-eda7-44ca-8c01-958309009ef8 | 625 |
| SKCM | TCGA-EE-A2GB | 2dd72796-7cac-418d-a092-523f88e1dd64 | 310b2515-640a-417e-9a35-bee64f90ac1c | 673 |
| SKCM | TCGA-EE-A2GC | 0c42714c-45d4-4591-bb02-c4f3116a72ea | 5949ea6f-17c0-4445-98ef-38a1e483e726 | 618 |
| SKCM | TCGA-EE-A2GS | effe1a0a-bcd4-4a16-9d4b-62396920b6fb | 49f9daa4-da67-4d32-9183-82c94e8344b4 | 288 |
| SKCM | TCGA-EE-A2M7 | 7212211c-0722-4a90-a452-49224f4e3e43 | 42bab265-08ca-4684-a949-bfbfe42eedc1 | 355 |
| SKCM | TCGA-EE-A2MC | f0ce1c44-2f33-40d9-842a-55269b546ed4 | a6801df2-863b-49c3-8b02-52bbb3536e41 | 358 |
| SKCM | TCGA-EE-A2MD | d129b319-ccf2-4715-9abd-3890ff13945f | 036982a1-44c4-415d-bccb-fc3d2738a8f4 | 276 |
| SKCM | TCGA-EE-A2MG | e2184793-28bf-4bf1-b685-2958c8c0b8a3 | 0bf941ea-fb16-4935-9af6-a05d078fdf99 | 414 |
| SKCM | TCGA-EE-A2MJ | b16f81f2-a92c-4e0d-8f13-16041c8c6cf5 | ef89481c-65ce-4d5e-8f55-3544d9b1be29 | 237 |
| SKCM | TCGA-EE-A2ML | 2336ae9f-4b2d-45d1-853f-e0ed0b651e71 | 2eb86a50-3721-4e6d-9fac-46ce15804128 | 333 |
| SKCM | TCGA-EE-A2MN | 73490ea3-b79d-4ba3-a8ef-250ff13e4c21 | 4dd5f91c-db26-4412-82e2-69e29de55207 | 175 |
| SKCM | TCGA-EE-A2MQ | 55e99af5-5846-42c7-a272-5ff5cf525357 | 379ff829-20a8-487f-a935-55656f3311b8 | 96 |
| SKCM | TCGA-EE-A2MS | 4cc98a8c-1e89-4a0a-a8c5-4a23e7880ac4 | ff803ed6-50bf-4221-a401-0cc77d8299a6 | 120 |
| SKCM | TCGA-ER-A197 | 63175366-aa4e-4f4a-99c8-5d6985a153a6 | 256ef618-0c68-4c1c-b754-2fb29870cf2c | 66 |
| SKCM | TCGA-ER-A19A | d62acfb8-3c88-47ac-b523-665ad3712081 | 9bcb5eb0-7146-42d1-bb51-a661b93142ac | 686 |
| SKCM | TCGA-ER-A19E | d3874936-da5d-479f-8c3c-1f3d9306dde2 | ddd1871b-91a1-4f2f-acd5-a69ecf18c07e | 1,770 |
| SKCM | TCGA-ER-A19Q | d8a90220-2acb-4d71-9feb-6febee1f8745 | 34899f85-566b-41ea-a927-a0707757d97f | 287 |
| SKCM | TCGA-ER-A1A1 | a0dd4823-2335-43ba-8e32-a91e08f80afd | a929309c-6fce-4ba9-ab55-aca676e8c78d | 211 |
| SKCM | TCGA-FS-A1YX | 6126185a-3723-4e2e-8299-9b572802970a | a2b998a3-6ed9-4244-b31a-5c256868a3d8 | 876 |
| SKCM | TCGA-FS-A1YY | 4c946768-7b87-4e7f-97eb-e35ca0ffdd80 | 80ab908d-d526-4748-b549-a33d238a0ddd | 175 |
| SKCM | TCGA-FS-A1Z3 | c18e2956-90d4-4a37-8841-e3b090936c2c | b12e8334-b794-42ce-97be-51daf06e6962 | 400 |
| SKCM | TCGA-FS-A1Z7 | 3e98b38f-dcad-4b32-9291-c9cd6b59c84b | affbe8f2-deb0-45e1-a584-9c4d3b4f981b | 682 |
| SKCM | TCGA-FS-A1ZE | 7e15422f-a494-44e1-a671-0b2ec886c3e2 | c24d750d-c28d-461b-b1b0-31a50556c535 | 188 |
| SKCM | TCGA-FS-A1ZF | fc766dcd-902b-4505-bf3f-e747d286eec1 | 33a85bcf-963a-4790-bd1f-1c7d28f1db24 | 344 |
| SKCM | TCGA-FS-A1ZG | 0cac5d8a-2184-4f80-9dc3-2ad5a8d9f0de | 0121e582-d1b1-4c5c-a9be-8ef4511b3148 | 511 |
| SKCM | TCGA-FS-A1ZH | a2479881-3488-41b7-be08-204b2027d50f | b643b41e-59b2-4796-99cb-3c40e021942f | 269 |
| SKCM | TCGA-FS-A1ZK | 04965cc3-824b-486b-9698-336c746b5e30 | 99af9f9f-fd5f-41ce-8394-6d9210c013a8 | 2,273 |
| SKCM | TCGA-FS-A1ZM | 3f7ff4ec-1e07-4f3c-a649-4819d0db712c | 802e023c-a4de-478e-8d5a-10bd9d163dec | 632 |
| SKCM | TCGA-FS-A1ZR | 3dc9427b-527d-465c-bfd9-7593de78af96 | 474ef1b0-6baa-490c-a7db-37e3d083bc3b | 139 |
| SKCM | TCGA-FS-A1ZT | 0a573a63-c701-4756-9f53-242b32237d44 | 174a6efc-f288-454f-97ae-e05d238bb3d9 | 380 |
| SKCM | TCGA-FS-A1ZW | 5e95e70e-4afd-41fb-87d9-eaf4198223e9 | 7453f952-2bcf-495f-a111-5064baf2d40a | 309 |
| SKCM | TCGA-FS-A1ZY | aa1f701f-1d30-46f6-a60c-0d519147d690 | 38a9b4ea-ffdd-466d-a579-e92170d3b596 | 304 |
| SKCM | TCGA-FS-A1ZZ | a063d15d-a203-4c9c-ba64-60b38c6876fa | 23022644-31a1-40a0-b7eb-97e14bec2d3f | 276 |
| SKCM | TCGA-GN-A262 | c7719432-272b-496c-9ca1-743d2a8c08ef | a59783ac-70a4-44f6-92ac-c5d6fdb3a8ee | 1,706 |
| SKCM | TCGA-GN-A264 | c2a3d9c8-b29b-4246-be90-70b59cdcc922 | ee630afd-acae-4b74-8ebc-ed5e124fd230 | 633 |
| SKCM | TCGA-GN-A265 | 95729723-84b0-4360-acd5-b7f42668ae18 | 8ad44bb1-82fb-42de-8306-68b2e86c747f | 200 |
| SKCM | TCGA-GN-A266 | faac710c-c974-41c3-ba2d-102952992342 | 59df09f9-be81-41ce-bb63-1a6f3ce58efc | 309 |
| SKCM | TCGA-HR-A2OH | e5833102-da03-41b2-8042-99e9d2c36758 | 2a69954c-008f-4a79-a1eb-0e0307cac08a | 309 |
| LUAD | TCGA-05-4249 | 94e6a5cd-7ba5-4f3e-bf83-7d07522963b4 | 61a4cbac-f493-484b-bbd9-4a2745d89f22 | 523 |
| LUAD | TCGA-05-4250 | b7bbb240-9e61-41e0-a669-1744d0bda12b | e2b88393-7111-4c16-8b98-344fb8aa701f | 15,264 |
| LUAD | TCGA-05-4382 | 42d76c95-55a7-4160-b22f-4338315e6146 | 12d07a36-94e0-4d6d-8625-86f383684a7d | 240 |
| LUAD | TCGA-05-4384 | 985192a3-1403-4393-895a-ca9a4d12268f | 3b48426f-446f-4943-a27f-fe59bd1c1820 | 805 |
| LUAD | TCGA-05-4389 | 0979871e-d324-4f7e-98da-f65ff17a74d7 | 8bba0a8e-5113-4344-9cdd-3abfa2e10d55 | 10,470 |
| LUAD | TCGA-05-4390 | 625e3715-ed9e-4946-8c92-f5a93e92e6d4 | ad4b348b-351c-4edf-b95d-e49dfe7c5de7 | 1,922 |
| LUAD | TCGA-05-4395 | dcb07672-8bde-427e-9d94-ac644e457e91 | 4e22fe2c-0d95-4a62-9b9b-af0b7b0839c1 | 63 |
| LUAD | TCGA-05-4396 | 61bb72a1-5c87-411d-af04-3e058c2beab2 | 09cacbbd-7331-443c-8ac0-78a0778344b5 | 34 |
| LUAD | TCGA-05-4397 | 48ff480a-fd08-4db2-8ba6-ce90769d4811 | 952667bd-0130-41d5-8e5e-d419d6ecc6f9 | 91,796 |
| LUAD | TCGA-05-4398 | fc9ee166-a6ea-4c5a-9978-b63277aec83e | c5182685-5c5b-42dc-9a85-4c0076ee389a | 1,941 |
| LUAD | TCGA-05-4402 | 603e16bf-9b05-49c9-a925-f409626c969a | 30d6ff50-187c-495a-8364-d0ae343f89c2 | 68 |
| LUAD | TCGA-05-4403 | 80e2118c-7ebf-42fc-8600-573afcbb3fe6 | 09d1673e-6daf-469e-ae53-19559464d5ed | 78 |
| LUAD | TCGA-05-4405 | 47fe3db4-7a54-4703-b3a4-7ad7def2469c | 5ba6a451-8249-4373-b08c-91c603ad51bc | 689 |
| LUAD | TCGA-05-4410 | bdcc8e9a-8d27-4167-ae6f-3c41b38c6c51 | 53af425f-3279-4968-bc2e-56cc1f8678b3 | 2,918 |
| LUAD | TCGA-05-4415 | ae778c61-5e62-4f34-9b62-f28d9acf21b1 | 57cfe098-8cf8-4aab-ba84-1d57f4617af9 | 1,832 |
| LUAD | TCGA-05-4417 | f4f3791e-3b72-4f33-986c-dca1cf3ada94 | e22b68ac-9772-4665-b367-38eee8691898 | 1,277 |
| LUAD | TCGA-05-4418 | 961e7ff2-7efc-46b6-9fe6-3f19366eeab8 | ba806a17-fb08-48cf-8ed8-069510bc4e50 | 40 |
| LUAD | TCGA-05-4420 | aa05f7cb-814c-4ddf-a639-50291fa19a95 | 803b9ff0-11e8-413f-8486-c9ad1ea350bc | 21,531 |
| LUAD | TCGA-05-4422 | 9fcb53c6-7d0c-4468-8a77-40e6ea3d43d8 | 735fb0c2-720d-444e-a61b-218fc42c9a1f | 665 |
| LUAD | TCGA-05-4424 | 1e635509-6178-417e-9452-bc77482ee8ed | d10fa246-b319-4797-ba50-e80f1686b364 | 356 |
| LUAD | TCGA-05-4425 | bf137f71-691f-42f6-a29c-e37a457bad18 | bfe20429-49d0-4a72-a8e2-5254fa77bbc1 | 780 |
| LUAD | TCGA-05-4426 | 63037227-adb0-4505-bd71-2527743ec736 | 7e65d1f5-4103-44eb-a39a-9944b9f86be9 | 238 |
| LUAD | TCGA-05-4427 | 10c44e9b-d94b-4f27-b275-2dab46a863c6 | 9d47bafd-adc4-4120-b715-fe1ecbf86b82 | 1,686 |
| LUAD | TCGA-05-4430 | e4ae6498-889f-4900-a73c-2e61a6e154d1 | 794fbc75-e590-4df0-9b98-9e113eef5c46 | 200 |
| LUAD | TCGA-05-4432 | 6110bc85-6f14-48d5-b166-2b3f42776573 | 10b90006-c305-434b-a1e6-0d2d780fe84f | 117 |
| LUAD | TCGA-05-4433 | d1b6064d-6736-4aad-8ad3-70ab64c92e3d | 5d83cd06-942c-4ea6-827b-4d1fdb9c4cee | 237 |
| LUAD | TCGA-05-4434 | f43f1ef0-b383-4a17-a3cc-25c353416799 | 9a0b4c83-372f-4226-889a-a732c74385ac | 188 |
| LUAD | TCGA-05-5429 | ce2afa67-8778-4902-aede-8362a8556f53 | f3d21928-332c-46d7-8a92-bafee2ed961c | 6,591 |
| LUAD | TCGA-05-5715 | c666800e-9d3e-4b96-8473-8b23fcd54503 | 93ca0971-dab1-4845-ad59-48fe54014068 | 1,839 |
| LUAD | TCGA-35-4122 | 3a18de12-d101-42f5-9eb4-f1726b519b4e | 17ab570d-b678-436b-be4b-61d8ca689578 | 3,181 |
| LUAD | TCGA-35-4123 | c620a587-dd69-4a1c-81c9-5ae742181fbc | 49076069-b689-4bbd-984c-7718df09a20e | 49 |
| LUAD | TCGA-35-5375 | 94c5194f-ea42-4739-b18a-4d1c8a4a9bbd | 4d89a22f-819e-43c7-889d-ae79cbbd0a31 | 48,478 |
| LUAD | TCGA-38-4628 | 2fb6e0f6-487d-4404-9958-6ca019a21395 | eb3c2eef-32ad-4303-bb2d-a79c9e8fe7d5 | 3,259 |
| LUAD | TCGA-38-4629 | ecb1f536-930a-4c20-a11a-c69c6740295f | 364e81ba-b440-470b-a9e4-8e3795d66795 | 328 |
| LUAD | TCGA-38-4630 | 1ece9db5-148a-413e-9dc4-bf65d88ecc5f | fc60deae-7143-44b2-82d5-4a904a986c32 | 21,616 |
| LUAD | TCGA-38-4631 | b1dd06d4-cd7d-4e62-8deb-12fed31731b5 | 5ae01470-ac1e-459f-9598-e44be31e416a | 5,613 |
| LUAD | TCGA-38-4632 | ee38fc59-2812-43a0-b6c0-ab684daab63d | 64c992dd-605a-4f70-b4a2-240b669c9e61 | 4,967 |
| LUAD | TCGA-38-6178 | f309d501-6d91-40ce-839c-bef9a434bae1 | a780b7c3-dbec-455d-b43e-7865f892c084 | 923 |
| LUAD | TCGA-38-7271 | 3d353009-70c5-4b0c-859e-8781ae224de4 | 2721100a-5893-4e96-b684-0e17eb4b1779 | 353 |
| LUAD | TCGA-38-A44F | 9103b22b-01cc-4dfd-945a-8f8bf2bbe312 | cfabe3c8-1dbc-4aab-b869-2b01606490eb | 394 |
| LUAD | TCGA-44-2655 | 6637465a-ebf9-481f-a7dd-9a0ba0f99757 | 25d361ca-11c2-4392-a688-79a000a8e01e | 608 |
| LUAD | TCGA-44-2656 | b0caa073-dc33-4f0a-aa7c-757dc1bf2718 | 0e4eaf03-9b2b-44d4-a50e-d2c306a45f8d | 35,019 |
| LUAD | TCGA-44-2659 | bcdd7201-d7f2-45f5-bbaf-e0ca63b987c4 | fb53288f-0723-4e89-a768-e236489cce5b | 28,184 |
| LUAD | TCGA-44-2661 | fbe0b7e0-45f4-4c61-9965-be52ae790251 | d921a1d0-fc6b-4140-adea-0274147e2026 | 34 |
| LUAD | TCGA-44-2665 | e8d44df6-c3ec-4eb3-b4aa-621dbc3c6cb6 | 7b1ce680-03d0-4279-871c-e96c91a60a8e | 692 |
| LUAD | TCGA-44-2666 | 7f2cc312-2955-4066-877c-b5bdb615fb49 | 1be0178e-fdc1-4ffe-809a-659dc052336a | 1,719 |
| LUAD | TCGA-44-2668 | 4b06cba9-58da-445f-8d45-46d13015a14f | be23b200-8888-4fbb-8809-bd6616a20b1d | 2,223 |
| LUAD | TCGA-44-3396 | bf55243b-c7c6-46fc-9d54-e1916b0d7ea2 | 03abf6dc-ed65-46a4-bca8-920d74f39139 | 2,718 |
| LUAD | TCGA-44-3398 | 60476396-b9c1-4093-82c0-5663fbd13049 | 2fae14bd-079f-464e-9fac-16ded87e8205 | 250 |
| LUAD | TCGA-44-3917 | 32392802-b62f-4779-8127-0a1cba3669ad | bdc2fd38-8b39-4e6e-84ae-dbfc8163fefb | 16,411 |
| LUAD | TCGA-44-3918 | de9eb245-c084-47d4-aa74-c451d0288296 | 297c5d46-e808-423b-9a00-4da7e03a6cc5 | 10,305 |
| LUAD | TCGA-44-3919 | 14d2cfc2-f4ac-4ab1-93d3-0e178305e26f | 0b6ce057-abf2-4149-a103-a243ae3d9354 | 6,962 |
| LUAD | TCGA-44-4112 | 7eec19b6-a5b4-4cf5-9470-08b7c88e2a16 | 9efbc36a-a54f-4ad7-ad07-1fc79001229d | 26,414 |
| LUAD | TCGA-44-5643 | e485193c-f3e0-44a0-8a09-3ffd4e90400e | a4f575fa-63a7-440d-974c-36caaf9a4c96 | 1,237 |
| LUAD | TCGA-44-5645 | ee5a021d-f489-4c9e-9b66-8a118be13fac | 046d2d90-c237-4a78-9661-79758711edfe | 436 |
| LUAD | TCGA-44-6144 | 4b5e64a9-e315-4b7c-949a-be7e2460b2df | adca72c6-ec26-44f7-83f6-27fcfee47296 | 458 |
| LUAD | TCGA-44-6145 | e000212a-50a0-4120-972d-772fbce61634 | b21d12d5-2762-46b6-bd9c-d2f3579c9779 | 1,999 |
| LUAD | TCGA-44-6146 | 8048300a-5c05-4e39-ba4a-b8b1921b419c | f7ee8faa-879c-4d5c-ac00-24452fb7fe3f | 175 |
| LUAD | TCGA-44-6147 | 2fbcfbf4-ee8c-44a1-bf31-a141b65af69f | ea7b7b64-8b4e-46fd-91d3-0a22729856f0 | 3,931 |
| LUAD | TCGA-44-6148 | e3f95d46-09ce-43a0-a2c7-581cb9fe8da9 | 7713e797-fc02-4096-8f27-9a4d1260df7d | 1,582 |
| LUAD | TCGA-44-6774 | 5ff8e7e5-3b91-49af-b384-9d870bd9e425 | b2c21986-7ec8-4686-89b3-45fc5f257b49 | 166 |
| LUAD | TCGA-44-6775 | c17b3c9b-e41f-48e2-9cde-facc8537afbc | 5628940b-6674-4130-b526-a3c2921984c9 | 366 |
| LUAD | TCGA-44-6776 | bb5d193c-cc97-466b-9b28-bdbf5cf255e8 | b7a6be76-449b-46fc-863f-1e1cb9498eae | 844 |
| LUAD | TCGA-44-6777 | 8f0fdfb5-6a7c-4e9a-9a4f-d32f9aca542d | e60ccd22-0cc2-4758-8098-0d5f331aceb1 | 1,633 |
| LUAD | TCGA-44-6778 | 8d0de456-4a8c-48ad-bfc5-fa6e66ba6e0d | 81a9ac95-7603-496d-a966-9fa6ebf33f49 | 16,449 |
| LUAD | TCGA-44-6779 | ab458a7b-afac-4c72-8db7-45994af6ceb8 | 85ba6328-d485-4bdb-8720-4dca67560048 | 406 |
| LUAD | TCGA-44-7660 | c83224de-2c8c-4f4a-a4b2-8384ec63e9d7 | c2df8a7a-b32b-4440-b773-0e4ef51b1f1e | 36,272 |
| LUAD | TCGA-44-7669 | d3907e52-d25b-4e1e-8c9d-1dfafdf808dc | 21226920-f452-4737-93a4-a1345c7571e4 | 41,587 |
| LUAD | TCGA-44-7670 | a1a876dd-e103-44a4-97e6-888f754cab18 | caa805d4-d58c-4205-a240-2178f0ae585d | 112,911 |
| LUAD | TCGA-44-8117 | 1e76dcec-7e26-4337-be45-0c6dd599ffcb | 4b101431-af13-4893-ba8e-125837680a47 | 72,996 |
| LUAD | TCGA-44-8120 | dd58b39c-ee83-4f82-b3ce-b2f1e0c911f0 | 9b2ee900-b749-4570-8a9d-00c14cf13d56 | 48,018 |
| LUAD | TCGA-49-4486 | 53a1d9af-aa10-4382-bfad-f13af3c6cab9 | 83ce2c4a-f55f-47bf-af39-3fe89232672b | 76 |
| LUAD | TCGA-49-4487 | d416671f-aea2-4667-ac81-d59839ae7ef8 | 72bae131-f61e-4fca-ad63-74ce48e1f32f | 1,864 |
| LUAD | TCGA-49-4488 | 26421230-2518-42aa-85ee-f9c83d47a6ba | 1a64c497-725c-488a-a6a9-c32abe246dea | 1,851 |
| LUAD | TCGA-49-4494 | 3cb48d09-95ac-452f-9c92-19d7f966aec0 | 56ee8c8c-9b38-4430-97b6-3272b9f84d68 | 66 |
| LUAD | TCGA-49-4501 | 2df919ad-215d-40a5-8e9d-e22862898cd7 | 2e94d307-de8c-4849-a870-8731a029c297 | 69 |
| LUAD | TCGA-49-4505 | af550be6-0986-458b-8206-9c7cf928c215 | edd13c78-4a11-4932-8158-20f922466310 | 90 |
| LUAD | TCGA-49-4506 | b07f7ea8-c74b-4559-97d3-987f1985f2e9 | 30c3d956-3fe6-4cf5-9829-00573c3a5813 | 198 |
| LUAD | TCGA-49-4507 | 557606c6-d245-45ba-b9f0-adc05c6a5f4f | 85ba42cf-c97b-4bd6-910f-d3e06fb69a79 | 263 |
| LUAD | TCGA-49-4510 | abe8920d-80b9-4fc8-a249-5bf0a4a06ac7 | b8bf16ce-9774-4ef9-a278-487ae6ec947e | 4,292 |
| LUAD | TCGA-49-4512 | bfba789d-2b84-4c34-89f9-7a38586facf6 | 425e9966-6cdb-4456-8501-ac186279082f | 9,707 |
| LUAD | TCGA-49-6742 | 2fdfd442-4aa3-49d0-82e8-7b21b059e7e3 | 55cf964d-5dc3-457c-a7c3-0ec44617951b | 1,125 |
| LUAD | TCGA-49-6743 | 8b2cdd00-4024-407a-94a7-dbfe5eaa6cd1 | b26d0042-8fd0-41ca-8236-a6509446e036 | 14,481 |
| LUAD | TCGA-49-6744 | 8290093a-0193-41fa-8e87-71960e0e640e | f185e615-65c6-4c02-aa8a-77c17e89e5e5 | 241 |
| LUAD | TCGA-49-6745 | b4eee7d7-a56c-4927-929e-ac66abe41408 | d9c97389-6652-4cde-82d7-7b8704ed1f0d | 283 |
| LUAD | TCGA-49-6767 | 759c1dd3-0f7d-4ba7-8c8e-c276faf0255b | a55991e3-e5fc-4ba4-b7e8-a98bfc625775 | 109 |
| LUAD | TCGA-50-5045 | e3bc7a3f-c296-412c-8974-8c21cf01dbfb | 425edc83-a2c5-4ada-be24-0e87cc319fbe | 169 |
| LUAD | TCGA-50-5049 | 16ea05eb-f830-4b36-83bb-989c8de99568 | 3a6b8979-927a-418d-971c-e268a67a00c1 | 584 |
| LUAD | TCGA-50-5051 | fd2a1c20-3ed6-4057-8fb5-4027ce0eec74 | 8d384520-9763-4a0c-ba5d-b99274779d43 | 213 |
| LUAD | TCGA-50-5055 | 97723d64-755b-4f00-b641-95ad083c397e | d1b8082f-27b0-4a78-990a-3de7ee79daa3 | 353 |
| LUAD | TCGA-50-5930 | 7ef1f666-79cc-4c94-90b9-49c32631ba72 | 8e682cfc-cf44-4c71-b08e-152f8539f402 | 5,037 |
| LUAD | TCGA-50-5931 | 718cd53d-80a6-4f5f-94d4-247b4ff5bd01 | 743fd57e-59cb-4ad1-986d-556950fbb2b9 | 47,137 |
| LUAD | TCGA-50-5932 | a1be116a-f392-49b3-97c6-b674ac8dee93 | ec0e95d7-b66b-415c-9d7e-c47759b2411f | 388 |
| LUAD | TCGA-50-5933 | ca97b42b-d723-4c0a-8f71-819eca1bb805 | 0cb84810-ecf1-4b3a-99d2-525d6d78e81a | 483 |
| LUAD | TCGA-50-5935 | f4ad5c7d-a36e-4d9a-9c36-444e4b443953 | 32fab0c6-8503-4996-a31a-4968a251eaa8 | 211 |
| LUAD | TCGA-50-5939 | 7a3d6187-a909-4946-83bf-0d6fc6a753ee | 67afa188-5949-4813-8056-994a819a72b5 | 1,101 |
| LUAD | TCGA-50-5941 | f7c353bf-7be8-42d0-88d0-e89032718c2f | 30af8e89-edaf-467e-a92e-6c6387c651dc | 685 |
| LUAD | TCGA-50-5944 | c76f3861-5117-4ed7-9ca6-eecc10e949f3 | 8879c93f-169f-4772-9556-86a9ed97e824 | 97 |
| LUAD | TCGA-50-6590 | 5dbd186c-2a64-4bbd-8889-3fdc26de58f4 | f63c14b0-0ace-42b3-85c7-75722e8c0f57 | 31,427 |
| LUAD | TCGA-50-6591 | 3d4858b4-9af4-47f2-84d2-6618c7a21581 | 3cf01f5f-1d8b-46a3-85da-9d9a16ff694d | 192 |
| LUAD | TCGA-50-6592 | 207da806-7a0b-406f-aaa6-b5bb17f7f28a | e1284c11-5577-48dd-b880-07c391fb1cc6 | 205 |
| LUAD | TCGA-50-6593 | f552d49e-eaf1-4d8f-bd11-0f2017c95ba3 | d6164f86-2486-463f-897c-fa4c2934fd35 | 1,235 |
| LUAD | TCGA-50-6594 | e9b1ca6e-b0f0-48d8-ba44-ebfcec8cd171 | 29591d1b-39c4-4087-a852-643e35cc55b0 | 943 |
| LUAD | TCGA-50-6597 | e7110ba5-6000-459a-8128-32b420e21194 | cf5182e3-08aa-45af-8c6f-2d6afdf01d08 | 500 |
| LUAD | TCGA-53-7624 | 016e6739-9184-4b6a-8c08-20aef965b333 | a3d8da32-e91b-4b15-be68-c8823758debc | 101,409 |
| LUAD | TCGA-53-7626 | f733ffe7-6b5b-4a67-8f6c-68c498f2daae | 8c4610f2-187b-44c8-a9d7-19f269d0de7e | 9,292 |
| LUAD | TCGA-55-1594 | 692f523c-9fa5-429d-ae27-3c0b2d1af53d | 74601a5c-3f0c-4a39-9025-4ee7f3cf35dc | 12,262 |
| LUAD | TCGA-55-1596 | bfb2ff32-a7a8-4df0-b8fd-ddcfec0fae22 | 2a257d09-8210-45fd-920a-efb7008724bd | 20,753 |
| LUAD | TCGA-55-5899 | 025bbc63-6a66-45e9-9a3f-d8c6db518c3d | 58e4bc69-e0ad-41de-ad55-c83ae97417d7 | 54,483 |
| LUAD | TCGA-55-6543 | 0715b83e-fcd8-41e9-88ac-cfeb5fabd3de | a66c52d0-6f85-4b19-aa52-97c7fd45d6f1 | 2,315 |
| LUAD | TCGA-55-6642 | 9b12800d-6911-4d3f-a25e-4d1b686b63aa | ae204af3-603b-4186-b84a-4634f70af495 | 2,947 |
| LUAD | TCGA-55-6712 | fb1682cf-0fca-4bf1-963d-67a3a177fe98 | 88cd05e2-cda3-4c5c-b350-f06a08f7be13 | 1,798 |
| LUAD | TCGA-55-6969 | 75739358-0308-44ea-a831-6173365dcfdb | 331d3e8a-6093-4a6d-bac1-a0d8631f36f5 | 8,722 |
| LUAD | TCGA-55-6972 | 30f0b521-90e2-41a2-9dcc-97b6702a46fc | 03a2b1be-957a-46f6-afb1-88c725c5b5e6 | 28,010 |
| LUAD | TCGA-55-6979 | 953fb4de-3be8-4d07-8ccd-c4d8ec07ded0 | 98c8ad65-c4a4-4835-a074-42fd1521491c | 1,148 |
| LUAD | TCGA-55-6982 | c6027d22-f05d-42e4-a467-de21b4606751 | 30195f83-7677-450a-83b9-06eeeda2a008 | 551 |
| LUAD | TCGA-55-6984 | 244af712-e56d-4419-bc80-cdbfc0262b92 | ca4b6c95-902e-453e-8a03-292a01304628 | 742 |
| LUAD | TCGA-55-6986 | ee88a53a-b82b-4bd8-b13b-da8f5da9e39c | f1b91760-f564-4921-a6ff-13403a3c0b19 | 298 |
| LUAD | TCGA-55-6987 | ef4a9dab-c7e4-4e4e-ac00-a8609da26448 | 8a0cf7ac-0e97-4f49-bda8-bd7edfc7f5cf | 930 |
| LUAD | TCGA-55-7281 | 19c6ac35-62fd-4db0-8641-22abc19d0b52 | 38a96bd4-b9ef-4b20-a45f-e1b0fcd37962 | 29,404 |
| LUAD | TCGA-55-7570 | 9cd3a11b-1814-49f7-9601-fd009349f582 | 4eb38fe0-2e12-411a-a69b-86e19db40ef2 | 33,706 |
| LUAD | TCGA-55-7574 | f667764c-3f48-45eb-87c3-a79aacce798d | e14622ff-c051-4f8f-9488-ee6622784fa8 | 683 |
| LUAD | TCGA-55-8085 | 1d193525-5a99-4ff6-9e84-781a2704866c | 1b163e21-5a45-48f5-91b2-861739443b82 | 17,218 |
| LUAD | TCGA-55-8091 | edf65852-49e7-4aee-9875-0a85a8eb2ae9 | 204e94a1-e687-4f3a-998f-c77501e7d455 | 1,292 |
| LUAD | TCGA-55-8208 | 291ff580-3d53-4f50-83af-b9e21884d878 | 94a2d171-3472-45f5-845e-28bba8ab0fc7 | 455 |
| LUAD | TCGA-55-8299 | 0900edf4-dd21-4cc0-a6cb-6de863318729 | cd560db1-53a9-4e91-88f1-abe2efc405a1 | 326 |
| LUAD | TCGA-55-8301 | f904fbfb-f7e2-49ec-a11f-dfdc3ab1f237 | 23d1b5c7-89d0-4139-a23c-9cd53e722ae4 | 4,178 |
| LUAD | TCGA-55-8507 | 381c4b92-cccf-43cb-ae37-bf8465f05d56 | 6a1860dc-7021-4d1e-b89f-856b957b0dcd | 88,500 |
| LUAD | TCGA-55-8510 | f95360b3-74f7-43be-8e06-90138c443199 | ee803e6c-83a7-4519-909b-ad7314416605 | 9,229 |
| LUAD | TCGA-55-8614 | 5361feb2-dc38-406b-ab12-de1eca7dc610 | 422b62d4-8d6e-479d-915e-a4edf5d38377 | 9,185 |
| LUAD | TCGA-55-8619 | 4ab403f0-6d28-4da1-8a8d-0ffa00cbb029 | cacde27d-3999-4a0e-9368-6282582dc6f1 | 401 |
| LUAD | TCGA-55-A48X | 5360e5c3-9296-469f-85e1-f54941ffe1c8 | ce205aa8-3f81-439c-9622-499f6c3ea7a2 | 1,136 |
| LUAD | TCGA-55-A48Y | 77f116c6-7659-4349-af5d-35d2a20264b3 | 330ad8c1-3728-4f0f-bc98-507a74cbb032 | 1,817 |
| LUAD | TCGA-55-A492 | 58ecf1ea-29eb-4b5a-92e6-7d78eb8c3c77 | 5aaa3fe6-f98d-4660-8184-f6ed134e60fd | 20,368 |
| LUAD | TCGA-55-A493 | ebc3aec6-f722-4515-91f0-8c3d79716f83 | 974a352d-e097-4756-abbc-fb4e0d9d39b4 | 998 |
| LUAD | TCGA-55-A4DF | f9a4942d-2386-4d4e-ba6c-bb017c13910b | 4a9f420e-c384-46b8-9b48-05fa6f6cbfba | 33,588 |
| LUAD | TCGA-62-8395 | f845ce2d-48e2-4f05-896c-9b4dd46eb138 | 33a90e78-cf44-4967-8606-27de08304d04 | 1,764 |
| LUAD | TCGA-62-8397 | f4382f43-e5a6-41b9-bea0-985c945a53e3 | f90725a9-f978-4106-b815-063779ca690e | 2,323 |
| LUAD | TCGA-62-8399 | 7711deb3-44cb-4f35-bb70-2df1911f32ac | 9801fdbe-87cf-49f6-bdde-4ed26e90f193 | 34,233 |
| LUAD | TCGA-62-A46O | 2404f57b-5282-4cd7-9826-149148d9ba14 | 6ffe9388-6a16-48cf-bb0f-284a5fa02c59 | 128,133 |
| LUAD | TCGA-62-A46P | ad09bc79-0009-4da7-b0d0-29c339dd657b | 3ef635ca-621e-4925-b690-547b671e1bfd | 33,961 |
| LUAD | TCGA-62-A470 | 2cd67c85-ed37-44b5-bc46-8bead5250fe0 | 06a88296-7433-4485-a7c7-8f40f16d3698 | 6,965 |
| LUAD | TCGA-64-1678 | 3abe422b-51c0-4fb1-932d-f9e07cdb2150 | e6876c57-9b2f-4df9-a630-09281125e6c0 | 36,180 |
| LUAD | TCGA-64-1680 | 6b9dbd25-b2c7-4ba6-8030-e98c1c8e9991 | 0e182f6d-70c5-46a2-a737-29990f3c472f | 5,832 |
| LUAD | TCGA-64-5774 | b5b4f9c8-7044-4c76-ae48-241c8c4dd953 | fee3375f-0641-4334-a876-7570f40bb0c0 | 1,223 |
| LUAD | TCGA-64-5775 | af9f3482-e78b-4fe4-ac01-64f69b84d78d | e35e2779-31f7-418f-8ed1-52c4af96d40c | 183 |
| LUAD | TCGA-64-5778 | bd5570f2-ed45-408a-a615-0a6808c31d84 | daa71055-8f7e-4087-850f-1669fc21e1dd | 204 |
| LUAD | TCGA-64-5779 | 0f2bcb97-fc17-44e9-8ae3-46e7140e78c8 | 03d5cde2-7328-454a-81fa-08bbf4770c5e | 163 |
| LUAD | TCGA-64-5781 | 0ff0be96-1698-468a-ac88-a3e6ffece079 | 233479be-ff10-4869-929b-83ada64ec434 | 28,626 |
| LUAD | TCGA-64-5815 | d64bfa4e-ae37-494a-aabe-6787ac58054a | 9ed58ced-f480-42df-be0a-fdc8a5c298f7 | 748 |
| LUAD | TCGA-67-3771 | 0ebe8891-f3c8-405a-ad50-93850699da24 | f8c0eb87-e277-443d-9ae0-6a8890763e83 | 59,535 |
| LUAD | TCGA-67-3772 | 71899006-7e86-49e1-9d51-c2bd90b4d74e | 45ff8274-fe76-4b92-9398-f2f9434295e7 | 424 |
| LUAD | TCGA-67-6215 | 502bd59b-5e8d-4170-9a24-b18fa50a5dc1 | 30aaa08e-3e7d-4c12-8452-ae8bfb552ef2 | 999 |
| LUAD | TCGA-67-6216 | ae4aaef1-8b47-476b-933a-ff69761aa0a9 | ca8f48ed-992b-4763-ad25-7d8f16528337 | 357 |
| LUAD | TCGA-67-6217 | cb2962fe-7f6e-4a68-871a-11d696bb5279 | 868f0059-3a66-4f4c-a285-0b547b522450 | 1,300 |
| LUAD | TCGA-69-7763 | 993ef879-1510-4459-ad29-c338fb9cb547 | 421d5924-3f89-47e0-ba9d-1ecef185647e | 3,159 |
| LUAD | TCGA-69-8255 | 1d322786-f630-488c-b9e4-a9cb904be6ba | 3999859c-adf9-4086-95de-9906d8fe6d8f | 22,286 |
| LUAD | TCGA-71-6725 | 9c1abbc4-5f6b-486c-a6a3-17a01c8783cb | 6ab86bce-d92f-4638-a2d7-4761bcc675d7 | 720 |
| LUAD | TCGA-73-4658 | 9ee6d96c-98e7-4829-94dc-f41e5f078854 | 154d1467-8cef-484b-b813-86a414c476d2 | 863 |
| LUAD | TCGA-73-4659 | 967b7c80-0ec2-4e5f-81fd-07bffeeeaa7f | f6587098-0071-4ecd-b925-ff4145189b87 | 10,204 |
| LUAD | TCGA-73-4662 | bb68d6e1-efca-4ccb-bbd6-45c4da207bb7 | 2018de7f-d9e8-4e7f-8459-9794814494b5 | 210 |
| LUAD | TCGA-73-4666 | 7af89597-2a6c-4030-acf6-4bd33963a3f3 | eb82e62e-f7d1-4c52-8b12-2d7dba1bd1b8 | 251 |
| LUAD | TCGA-73-4668 | c3c82aae-4b64-4f65-b4ed-048053aef739 | c21a88c1-b349-46d8-8008-d9bc9b8cf337 | 422 |
| LUAD | TCGA-73-4670 | 8e4449ab-641c-44ea-af17-a681182f7299 | fb70791d-abaa-4c78-8e45-77605f89d966 | 41,648 |
| LUAD | TCGA-73-4675 | 14b3ad66-64c9-4b39-a34e-5cb29788735a | 5c0c01fd-bde7-41d5-ab01-78909043c0fe | 132 |
| LUAD | TCGA-73-4676 | a2b68d59-a4d4-4793-9e6c-72ccaf7c242b | a85b7af6-36cb-40db-9085-3d24b08100ea | 487 |
| LUAD | TCGA-73-4677 | 5dd207d7-c666-46ae-8154-cb0c0cd6467f | 12b710fb-7df3-4422-baf3-33178cb5a9fc | 119 |
| LUAD | TCGA-73-7499 | 78a616cf-926a-41ac-9ca4-05f214046730 | 2184998c-4532-4650-b6b8-24a0bee8e6f1 | 2,558 |
| LUAD | TCGA-75-5122 | 611aeb0f-3dbe-41c6-8b8e-b4a7c827ac1d | b223fc0a-cce3-4c35-8a1f-fb5fceca402e | 336 |
| LUAD | TCGA-75-5125 | 043f5225-fe1f-4ea6-bf29-e657f8a6a9cf | 6ca16082-7c39-4191-949d-7b0103fdbf85 | 583 |
| LUAD | TCGA-75-5126 | 8c670948-9cf1-4db3-82d5-39ed824c61ff | c15f5681-638a-4471-a349-56138d68fe62 | 530 |
| LUAD | TCGA-75-5146 | 926ac8ca-ff8f-4822-bf62-819bf5525a5d | 22cccb1d-1901-4369-8c11-24895005a4df | 39 |
| LUAD | TCGA-75-5147 | 6ec725e2-6d00-4866-8473-fd1b02309066 | 31e5136c-8e36-4019-8f00-1cab1d4a9303 | 7,166 |
| LUAD | TCGA-75-6203 | 976f4757-1570-44bb-9a05-7680fc9d30b6 | 279e57e9-744f-4530-a856-cf989245b8ef | 5,775 |
| LUAD | TCGA-75-6205 | ccc090ad-afab-4e7d-b71b-dcd39a87e359 | 247f390f-cb78-4845-a99b-65b33f838887 | 301 |
| LUAD | TCGA-75-6206 | 2b0c2142-3073-4443-9998-a7e909add85d | 56ce7501-9487-4758-8a1c-0068f71b89b2 | 405 |
| LUAD | TCGA-75-6214 | c39dcd60-dc0c-48fd-a2d0-0dce1533aaa0 | 43091020-7e96-4cbd-88a1-caa66438e030 | 50,320 |
| LUAD | TCGA-75-7030 | 804a0fba-a82e-4501-9b0b-87c34318562d | d5a3ce22-1fc8-401a-ac7b-fa7bc6fad8f7 | 439 |
| LUAD | TCGA-75-7031 | 64a5c836-4b06-4ce8-be27-9972344a8ccd | 90e99a72-5f62-4ded-8706-9b79d323609e | 15,793 |
| LUAD | TCGA-78-7143 | d0505bbf-c663-484b-93b6-a129883be59b | 06f9a2a0-93e4-4d22-a956-91e3d6a212ab | 1,143 |
| LUAD | TCGA-78-7146 | 17e2078c-ad94-4ecc-ba36-9a8e9e4e237e | e3e01e40-c359-4dc8-90ae-a703361e2439 | 32,150 |
| LUAD | TCGA-78-7149 | a8178fd5-6f16-4b5a-ab14-eb29c93860ca | a3e7aab6-6231-43f2-b740-8cf15ffcf171 | 8,599 |
| LUAD | TCGA-78-7150 | 6252fdf2-c74e-45c9-bcbd-0f0f1d5dee07 | 93c7e79e-4185-47c8-b321-1a862c2b1d80 | 36,351 |
| LUAD | TCGA-78-7152 | d399e786-9714-48aa-b798-8f1e518d494a | aeb3b08d-b17e-4c5e-911a-116b78dace7a | 7,274 |
| LUAD | TCGA-78-7155 | 814b0b2e-98a9-473e-a974-0946534abc6b | ae3d686c-737c-4183-a591-fd9705bc9a4a | 225,023 |
| LUAD | TCGA-78-7156 | f02c1668-47cc-4ebd-a656-5c2a70f1ae9f | 93ab8b16-f10e-47da-af99-cc812181135b | 30,774 |
| LUAD | TCGA-78-7158 | a905fe2c-571c-4769-9c35-b107679cad76 | 492a4551-71af-44ec-bbaa-f7642fc42054 | 33,035 |
| LUAD | TCGA-78-7159 | ff4430d4-f099-4e46-b76c-4a5d64d624a3 | e7498e1f-c471-4133-bc3d-e968b12ab793 | 25,890 |
| LUAD | TCGA-78-7162 | 4ca61254-01c1-44fa-9af3-8a1030f4757e | 8a0625fb-895b-4c34-8566-f0cdffbafe39 | 451 |
| LUAD | TCGA-78-7535 | c7601721-da6e-403a-9b43-7971f6513153 | 1a2489fb-ddaf-4d68-aec6-3707b64b8d96 | 16,457 |
| LUAD | TCGA-78-7536 | cdabc2dd-9276-4568-b7c5-05a212bf89a4 | 91b44ad7-543b-4dcb-915a-f95b1de48a93 | 80,104 |
| LUAD | TCGA-78-8640 | a15a69ad-a07d-43a0-9f93-4d9f04c265c6 | 0d5a6726-1485-4e2a-b6e7-21ab521fb83c | 28,358 |
| LUAD | TCGA-86-6562 | 71d960e2-40e8-4084-bcc1-5ebbefc183af | 02776cc1-3edd-49d5-b77e-8a9ec2d68f5a | 455 |
| LUAD | TCGA-86-6851 | fd54e659-2680-428f-ab83-afd6049b2c08 | 16f6006e-9cbd-4f90-97f5-b8ebdec258c7 | 44,378 |
| LUAD | TCGA-86-7701 | d37a81d1-a2d3-4ea0-abbd-3a07d5235bf2 | bb61f945-2211-41aa-82a9-0b54ca19c64d | 13,949 |
| LUAD | TCGA-86-7711 | 70f74bd2-94d5-4961-a742-f74c7029cad2 | cb12ffd8-08a2-4eb1-b857-67c7f73ea8a2 | 4,625 |
| LUAD | TCGA-86-7954 | a441e0b1-6145-4336-b207-b7be9e4c5d0a | c2fd9584-1d46-4f35-befd-3917b9d466ed | 3,787 |
| LUAD | TCGA-86-7955 | 7f50e25c-7549-446b-8804-08f3b310189e | ffcdfd02-69ea-4515-9366-5df60b395050 | 48,171 |
| LUAD | TCGA-86-8358 | 04f1ef84-2a06-4987-9d58-8dc77ddda9ea | 4a15cc25-73b0-445d-9cdf-2859e406fb2a | 148,768 |
| LUAD | TCGA-86-8585 | ce4071b9-5bee-4249-b4df-476a213da0e2 | 90f4eb67-280c-47e3-9da7-bfbc1f47ab69 | 25,969 |
| LUAD | TCGA-86-8673 | 01482104-71be-4873-82b1-d8f5f448b7e6 | 8dfe3545-cda3-4e83-8f20-f70abcda64ee | 18,612 |
| LUAD | TCGA-91-6828 | 94e2b6af-1f25-4698-9414-f3d916d26aee | a572c04b-c826-4837-94e8-72ccdc833516 | 1,431 |
| LUAD | TCGA-91-6829 | be9bb40c-b149-4b91-b271-b8881bb90d6f | a1eb6a43-9ab8-4c52-a1e2-a18576cf662c | 5,804 |
| LUAD | TCGA-91-6830 | 30b407eb-ca65-44da-9abb-f19b6def6a24 | 0036465b-ebab-4fa7-8525-ff035dba64a8 | 1,187 |
| LUAD | TCGA-91-6831 | 319d6c8c-207c-477a-8254-30695b75f9e2 | 668cb7d1-0943-47c0-a4fa-a0b61974412a | 23,840 |
| LUAD | TCGA-91-6835 | 8c5e1a0f-e9a8-4ce4-b066-97b2040c5327 | 842df4e9-6924-416b-bf5a-773944b4ba5e | 588 |
| LUAD | TCGA-91-6836 | 68fe0529-8b54-4c01-83f0-a5dc5097e2cb | 76fb7328-b5a3-43dd-b29f-412366ad711a | 4,117 |
| LUAD | TCGA-91-6840 | 27c919e7-a635-4c6d-a12d-31d1ee814e12 | ecb2ce47-ebd7-406d-9b76-31a08bc2531c | 6,895 |
| LUAD | TCGA-91-6847 | 7d8d3882-a579-44c1-b63d-eabff64671d1 | 251ee127-d60e-475f-8ea0-8724fa15c60b | 3,797 |
| LUAD | TCGA-91-7771 | 11bfd55a-b383-4b4c-adcb-d14190daaf06 | a094287e-6191-4b90-9189-9b35f33ced36 | 2,931 |
| LUAD | TCGA-91-8499 | 3bce5ae7-fb88-4a1e-a341-524b5ee95806 | 771e9142-d2f4-465e-8db2-636b7693476a | 35,285 |
| LUAD | TCGA-91-A4BC | 87ea14b6-2c41-4b1d-8bf8-b69af94b770d | e31b71cc-3223-4f6b-84cb-03bc66f4bcd2 | 9,875 |
| LUAD | TCGA-95-7039 | fcf5d66e-7475-4c3b-9c09-ea1a193f2632 | 789d86cc-94e7-45d4-ad6d-3a0daf7a41ab | 85,489 |
| LUAD | TCGA-95-7948 | 36ac2ce8-f4f0-4852-b74f-c5480a15e89d | 2115aeab-d5fc-44db-bb2a-634303b11a85 | 4,627 |
| LUAD | TCGA-97-7552 | d02ef824-2329-472b-aee1-6cead6432e8e | 9d23eeba-26a2-4947-9389-f4d742638e5b | 418 |
| LUAD | TCGA-97-7937 | be7a9163-98f3-4b37-8b95-358e90132d12 | 9312d02b-cb1a-4d00-9785-d8d755eca508 | 47,512 |
| LUAD | TCGA-97-8171 | e34a02d6-fc08-42eb-af7c-d4b40644bcf3 | ee38fa28-8a23-4ff5-ab4b-bda6fc6a60ce | 4,414 |
| LUAD | TCGA-97-8172 | c3713af1-85a5-4759-9b1e-ddb7e0cb67ea | b07d3b34-9a8c-4493-b004-6acac672829a | 9,559 |
| LUAD | TCGA-97-8174 | c1c45c3d-643d-48b8-b632-c144d2546e8a | a1765d8b-a943-4b7b-be30-e1a835a2c7f3 | 5,570 |
| LUAD | TCGA-97-8179 | 5f309873-0ce2-4d26-b98a-55b9ed5d9e84 | cd428d4f-661a-4d0f-adfc-d505cced4619 | 23,085 |
| LUAD | TCGA-97-A4LX | 0b18d800-8bb4-4f28-beee-a803c1020618 | bb1f0866-c162-4cba-a373-9127c8eafb7b | 280 |
| LUAD | TCGA-97-A4M2 | 4ec0322a-4d73-4d5b-a3fe-2b3c4cdc50df | b0c76d0c-f8cc-456b-a5e4-7d94d2c9f310 | 294 |
| LUAD | TCGA-97-A4M3 | 39955466-8f73-47fc-a64c-c4e7f13879a0 | 25b6e74e-e74a-4364-914f-58f045e29b3f | 13,348 |
| LUAD | TCGA-J2-A4AD | a76bf156-3b54-40f2-966e-3587f1ffd920 | 35dca363-a088-424f-9115-2aa50cc8e68a | 8,012 |
| LUAD | TCGA-MP-A4TH | 11c52e9d-e4fd-4d36-9834-b51526e2d873 | e8e55023-201b-49f2-aee1-0cd4506b6b4e | 2,371 |
| LUAD | TCGA-MP-A5C7 | 44d208ce-ff14-4773-b84d-89809fe71ca7 | ca455e36-c424-4f10-a827-e2d73edd93fc | 6,764 |
| COAD | TCGA-A6-2677 | 08a8edf8-3d2b-47e5-bc25-df07a165a2c7 | 57243607-fde9-46db-817d-b6e60dfece20 | 7,966 |
| COAD | TCGA-A6-2680 | d43e0ed3-ab1d-486a-bbf5-91dabacf8220 | 918f63e2-8d07-4be2-aace-c4515f9dfd6b | 6,322 |
| COAD | TCGA-A6-2681 | eda754a1-3a13-47a5-8e0e-49ef1384646d | 39e367a8-940b-4e1b-aa38-cdb0fdfe9d26 | 6,749 |
| COAD | TCGA-A6-2683 | 06c41745-bee7-471d-9c51-8886c6c8dd9c | 98608f88-13ef-4b96-91d0-d6f053a8a8a3 | 10,160 |
| COAD | TCGA-A6-2684 | adc6dd32-0ea9-4757-8daa-2c44f5d75699 | d8856104-2d3c-40a2-8938-f7a6b239274b | 5,237 |
| COAD | TCGA-A6-3807 | b5897ed2-6c9b-4c6e-b131-41b62e60ecb9 | 1555806c-03b9-4392-b3ab-82a34f015b0d | 2,772 |
| COAD | TCGA-A6-3809 | 5b346028-8694-4d79-a7f3-28263f4b171e | a13d12c2-ae81-49f6-b1c4-96d81ca82bc5 | 45,707 |
| COAD | TCGA-A6-3810 | 54687a8d-c693-4ae5-8451-4f4c80adca53 | f725df5d-8fb1-4c7b-b4c6-828a349265d2 | 4,555 |
| COAD | TCGA-A6-5656 | 9eb2efa2-318e-4cde-a110-8308c3792e1f | 7bd174e3-bace-491d-beb0-c781f3078e54 | 6,116 |
| COAD | TCGA-A6-5659 | 175be571-4e11-475a-aec1-dbebad709b6a | 4f4bf64c-edf2-418b-a5e2-e42011b44fc0 | 13,669 |
| COAD | TCGA-A6-6141 | 6fd83764-dbd4-43ec-bab9-c2950c46cf59 | a191bf4a-e494-4741-963f-04125c2a7d25 | 65,155 |
| COAD | TCGA-A6-6650 | 76422a6b-87fc-44d3-8b6e-b14ce682162b | 7addbeb8-cc13-4220-aac2-6ce1df297d91 | 7,266 |
| COAD | TCGA-A6-6780 | 7f5e0f27-fbbc-428a-9e5c-3a902b531462 | 85b53165-7f3f-43be-9ebe-3b80b7224046 | 38,078 |
| COAD | TCGA-A6-6781 | bee0f5f3-5654-4349-b3dd-afe0b89cc488 | a27b491f-dfce-4f7b-a18c-8ada5309959c | 31,329 |
| COAD | TCGA-A6-A565 | 94ddbe0f-9753-4acc-abe8-2bfda86dd8b2 | 1a964232-376f-4359-a91e-ba7dd9d9c03a | 432 |
| COAD | TCGA-A6-A566 | dd970455-c925-43f6-b58d-24982c846bdd | 0edbadde-12aa-44e7-a08e-3b7504787818 | 1,063 |
| COAD | TCGA-A6-A567 | 062aa65c-624a-4bc6-97bc-6c35e7dc0ba3 | c11fb256-ad9d-4750-ac63-bce5056c770b | 2,550 |
| COAD | TCGA-A6-A56B | 319d9aec-3e76-4b48-b4a8-9ffcb0f3acc4 | ee77fab1-ee89-43e8-b46f-8f15ee0ea73c | 5,287 |
| COAD | TCGA-AA-3514 | 2c81df5f-0948-463a-a303-3beb5bd5205b | 878b531a-852c-4466-8dcd-2f1ec7299ef6 | 8,337 |
| COAD | TCGA-AA-3516 | f25d539d-8289-4400-8da7-ddce46918578 | c5af0ea6-d2e3-4410-88c0-19d66b59324d | 88,863 |
| COAD | TCGA-AA-3518 | 7d2638fc-9574-4f2a-baf2-495e6ba99e93 | 20e838c3-2c21-4875-90e0-626174a3c1db | 68,790 |
| COAD | TCGA-AA-3529 | 8d96c289-9e08-45f8-a5dc-8c089ca7eb93 | c3a90e62-0feb-4728-99f1-8b461bfa7760 | 6,418 |
| COAD | TCGA-AA-3534 | 6e533853-5822-49cf-95a8-d75c517521cc | 6005aa4b-25dd-4a9e-9837-cb87da406ec0 | 7,462 |
| COAD | TCGA-AA-3555 | 6843fb20-8f01-4121-9e2c-823e5ce9da98 | e46a72b9-a8a8-41dd-af2d-3dadc0faf711 | 175,596 |
| COAD | TCGA-AA-3664 | 1eb3d869-c9bd-4087-80cd-7fb426eb7997 | bb549fb1-8744-496c-bf19-13b063fd8edd | 12,934 |
| COAD | TCGA-AA-3666 | 0bc29a5e-10c6-4c45-8cb6-2284ab7be445 | 58cda208-4b02-4391-9361-09d9fb432ba9 | 12,281 |
| COAD | TCGA-AA-3685 | e8516b3d-fcff-4f10-94a4-993b059ac4aa | 5997dc8e-b2ff-409c-ae44-75c132609542 | 3,174 |
| COAD | TCGA-AA-3956 | edcd6fc2-ca83-409b-957a-ff0199cedf0d | 10d20517-4670-47bf-b211-1c3989c3a7e0 | 5,085 |
| COAD | TCGA-AA-3977 | 34a6b8b6-5a24-4e03-9f20-8b51e27033c5 | f511ad35-378f-4391-a60b-6df71d8d62ab | 397,177 |
| COAD | TCGA-AA-3994 | dadc0f63-410b-4eb6-b287-c05bbc274608 | 0bea6476-1972-414f-be09-4c4197784248 | 11,102 |
| COAD | TCGA-AA-A01R | dc5cfaa9-514c-4e4d-b882-21df489668aa | e3101288-fc25-4e13-b4de-7cf4122567d3 | 76,526 |
| COAD | TCGA-AA-A01S | fe7c227a-1329-4956-b9bf-08bbb6fe161d | 272c148a-7ae2-411c-bdcf-153d522ba479 | 5,803 |
| COAD | TCGA-AA-A01T | 0dd44906-041d-4af2-8f42-a1bed185e1c6 | bb48e041-be2a-4f2c-83ff-39500c7790bb | 5,432 |
| COAD | TCGA-AA-A01V | f9fde040-dea2-4d98-8a28-05cb13efa1c2 | a0120ca6-fc3e-4ff4-b926-6c38e3180674 | 14,019 |
| COAD | TCGA-AA-A01X | 4b96a6ce-adbd-45b8-9c53-ea1850f0731c | 3f501377-1ded-4810-b320-714cacc4f71e | 5,168 |
| COAD | TCGA-AA-A02O | e53db54a-bb1b-4c60-8214-aaa269abd153 | 8fc33742-2fc4-4f40-8319-f7aa7d19f507 | 14,943 |
| COAD | TCGA-AA-A02Y | 258e501e-d510-4eda-b9fb-9d4bf0226c16 | 50ef042d-a98b-44e3-b1d9-85c89deaa267 | 8,934 |
| COAD | TCGA-AA-A03F | d227eb87-8290-4223-b1d2-c241162935c9 | 6363126b-bc8c-4b4d-9d0f-efe9e48683f9 | 8,305 |
| COAD | TCGA-AD-6964 | 67abf892-45e8-4468-a1fd-8446401b3f55 | 0378ffd0-fa85-4cc3-8efb-c0b12364d74c | 22,715 |
| COAD | TCGA-AD-A5EJ | ce0bf9ea-efa0-4c54-8970-a703051d667e | 42598f0e-7e52-4cd0-ab8d-06a385849f82 | 94,960 |
| COAD | TCGA-AD-A5EK | e84aec8b-4304-43e0-96c7-d605fc64dd11 | 3f5d539e-383b-419e-8904-03f97fd7ffb0 | 8,529 |
| COAD | TCGA-AZ-4315 | 8b6a06d6-c15c-4491-996c-2a012440f8a2 | 9a6322eb-3fac-4d37-bbec-89355ddc9dbb | 334,488 |
| COAD | TCGA-AZ-6601 | ae0bedfc-1ef9-4999-8ff2-c0eb8e421ef3 | 4db18c79-d142-4670-82f6-5d52be5fb5c9 | 19,241 |
| COAD | TCGA-CA-6717 | da552d40-23e1-469c-9fe7-ff2904a35ebe | 6896d05c-37ef-4347-bb21-6d3651e63b21 | 94,648 |
| COAD | TCGA-CA-6718 | 3f12f1a7-45cc-4500-b7e1-8ac12465fbfe | c3a26964-f2a1-4130-9280-3f40b76dd363 | 283,093 |
| COAD | TCGA-D5-6540 | ea4fb044-493b-4e48-ac28-28ba39b8be21 | 1f5b0f69-5647-47cd-9e0f-48e2bc982525 | 100,564 |
| COAD | TCGA-NH-A50T | 2893d404-6a2e-482b-86f7-eb3ae051faf4 | 80c16860-8c0f-4bf8-9d4e-58680115459c | 6,315 |
| COAD | TCGA-NH-A50V | 80f9989d-1696-49b8-a893-f045bc8615ea | b51c024f-b89d-4306-9e04-9720e3301d0a | 9,312 |
| COAD | TCGA-QG-A5YV | fc951188-ba57-4cd7-a209-28726710bb92 | 3b0443a1-23bb-40f0-bad1-f1fbc593e673 | 449 |
| COAD | TCGA-QG-A5YW | 32a205bb-36f6-42a4-8bc1-42318eeccd0f | 809c20cb-03fe-4d13-9715-d6cafb4bd36c | 10,306 |
| COAD | TCGA-QG-A5YX | aba79b86-0583-4858-8c09-d179c8a3b77e | 1c42d042-124a-40c6-842a-08d4cdca842a | 7,798 |
| COAD | TCGA-QG-A5Z1 | cc89a29c-287c-4a64-87c1-40ea6c90f58a | 62c3e1f9-6214-4f67-82cc-424f37f29d79 | 6,861 |
| COAD | TCGA-QG-A5Z2 | 2113677f-10e1-4987-bde0-35d29025de2e | 85b2ff62-069c-484b-b1cd-6bcde3e9c7a6 | 26,690 |
