## Supplementary material for "A catalog of *cis*-regulatory mutations in 12 major cancer types": Table S4

| **Cell** | **Place_or_Treatment+Antibody** |
| --- | --- |
| Dnd41 | Broad+CTCF |
| Dnd41 | Broad+EZH2_(39875) |
| GM12878 | Broad+CTCF |
| GM12878 | Broad+EZH2_(39875) |
| H1-hESC | Broad+CHD1_(A301-218A) |
| H1-hESC | Broad+CTCF |
| H1-hESC | Broad+EZH2_(39875) |
| H1-hESC | Broad+JARID1A_(ab26049) |
| H1-hESC | Broad+RBBP5_(A300-109A) |
| HeLa-S3 | Broad+CTCF |
| HeLa-S3 | Broad+EZH2_(39875) |
| HeLa-S3 | Broad+Pol2(b) |
| HepG2 | Broad+CTCF |
| HepG2 | Broad+EZH2_(39875) |
| HMEC | Broad+CTCF |
| HMEC | Broad+EZH2_(39875) |
| HSMM | Broad+CTCF |
| HSMM | Broad+EZH2_(39875) |
| HSMMtube | Broad+CTCF |
| HSMMtube | Broad+EZH2_(39875) |
| HUVEC | Broad+CTCF |
| HUVEC | Broad+EZH2_(39875) |
| HUVEC | Broad+Pol2(b) |
| K562 | Broad+CHD1_(A301-218A) |
| K562 | Broad+CTCF |
| K562 | Broad+EZH2_(39875) |
| K562 | Broad+HDAC1_(SC-6298) |
| K562 | Broad+HDAC2_(A300-705A) |
| K562 | Broad+HDAC6_(A301-341A) |
| K562 | Broad+p300 |
| K562 | Broad+PHF8_(A301-772A) |
| K562 | Broad+PLU1 |
| K562 | Broad+Pol2(b) |
| K562 | Broad+RBBP5_(A300-109A) |
| K562 | Broad+SAP30_(39731) |
| NH-A | Broad+CTCF |
| NH-A | Broad+EZH2_(39875) |
| NHDF-Ad | Broad+CTCF |
| NHDF-Ad | Broad+EZH2_(39875) |
| NHEK | Broad+CTCF |
| NHEK | Broad+EZH2_(39875) |
| NHEK | Broad+Pol2(b) |
| NHLF | Broad+CTCF |
| NHLF | Broad+EZH2_(39875) |
| Osteobl | Broad+CTCF |
| A549 | EtOH_0.02pct+HudsonAlpha+ATF3 |
| A549 | EtOH_0.02pct+HudsonAlpha+BCL3 |
| A549 | DEX_100nM+HudsonAlpha+CREB1_(SC-240) |
| A549 | DEX_100nM+HudsonAlpha+CTCF_(SC-5916) |
| A549 | EtOH_0.02pct+HudsonAlpha+CTCF_(SC-5916) |
| A549 | EtOH_0.02pct+HudsonAlpha+ELF1_(SC-631) |
| A549 | EtOH_0.02pct+HudsonAlpha+ETS1 |
| A549 | EtOH_0.02pct+HudsonAlpha+FOSL2 |
| A549 | DEX_100nM+HudsonAlpha+FOXA1_(SC-101058) |
| A549 | EtOH_0.02pct+HudsonAlpha+GABP |
| A549 | DEX_500pM+HudsonAlpha+GR |
| A549 | DEX_50nM+HudsonAlpha+GR |
| A549 | DEX_5nM+HudsonAlpha+GR |
| A549 | DEX_100nM+HudsonAlpha+GR |
| A549 | EtOH_0.02pct+HudsonAlpha+NRSF |
| A549 | EtOH_0.02pct+HudsonAlpha+p300 |
| A549 | DEX_100nM+HudsonAlpha+Pol2 |
| A549 | EtOH_0.02pct+HudsonAlpha+Pol2 |
| A549 | EtOH_0.02pct+HudsonAlpha+Sin3Ak-20 |
| A549 | EtOH_0.02pct+HudsonAlpha+SIX5 |
| A549 | EtOH_0.02pct+HudsonAlpha+TAF1 |
| A549 | EtOH_0.02pct+HudsonAlpha+TCF12 |
| A549 | DEX_100nM+HudsonAlpha+USF-1 |
| A549 | EtOH_0.02pct+HudsonAlpha+V04+USF-1 |
| A549 | EtOH_0.02pct+HudsonAlpha+USF-1 |
| A549 | EtOH_0.02pct+HudsonAlpha+YY1_(SC-281) |
| A549 | EtOH_0.02pct+HudsonAlpha+ZBTB33 |
| ECC-1 | DMSO_0.02pct+HudsonAlpha+CTCF_(SC-5916) |
| ECC-1 | BPA_100nM+HudsonAlpha+ERalpha_a |
| ECC-1 | Estradiol_10nM+HudsonAlpha+ERalpha_a |
| ECC-1 | Genistein_100nM+HudsonAlpha+ERalpha_a |
| ECC-1 | DMSO_0.02pct+HudsonAlpha+FOXA1_(SC-6553) |
| ECC-1 | DEX_100nM+HudsonAlpha+GR |
| ECC-1 | DMSO_0.02pct+HudsonAlpha+Pol2 |
| GM12878 | HudsonAlpha+ATF2_(SC-81188) |
| GM12878 | HudsonAlpha+ATF3 |
| GM12878 | HudsonAlpha+BATF |
| GM12878 | HudsonAlpha+BCL11A |
| GM12878 | HudsonAlpha+BCL3 |
| GM12878 | HudsonAlpha+BCLAF1_(SC-101388) |
| GM12878 | HudsonAlpha+CEBPB_(SC-150) |
| GM12878 | HudsonAlpha+EBF1_(SC-137065) |
| GM12878 | HudsonAlpha+Egr-1 |
| GM12878 | HudsonAlpha+ELF1_(SC-631) |
| GM12878 | HudsonAlpha+ETS1 |
| GM12878 | HudsonAlpha+FOXM1_(SC-502) |
| GM12878 | HudsonAlpha+GABP |
| GM12878 | HudsonAlpha+IRF4_(SC-6059) |
| GM12878 | HudsonAlpha+MEF2A |
| GM12878 | HudsonAlpha+MEF2C_(SC-13268) |
| GM12878 | HudsonAlpha+MTA3_(SC-81325) |
| GM12878 | HudsonAlpha+NFATC1_(SC-17834) |
| GM12878 | HudsonAlpha+NFIC_(SC-81335) |
| GM12878 | HudsonAlpha+NRSF |
| GM12878 | HudsonAlpha+p300 |
| GM12878 | HudsonAlpha+PAX5-C20 |
| GM12878 | HudsonAlpha+PAX5-N19 |
| GM12878 | HudsonAlpha+Pbx3 |
| GM12878 | HudsonAlpha+PML_(SC-71910) |
| GM12878 | HudsonAlpha+Pol2-4H8 |
| GM12878 | HudsonAlpha+Pol2 |
| GM12878 | HudsonAlpha+POU2F2 |
| GM12878 | HudsonAlpha+PU.1 |
| GM12878 | HudsonAlpha+Rad21 |
| GM12878 | HudsonAlpha+RUNX3_(SC-101553) |
| GM12878 | HudsonAlpha+RXRA |
| GM12878 | HudsonAlpha+SIX5 |
| GM12878 | HudsonAlpha+SP1 |
| GM12878 | HudsonAlpha+SRF |
| GM12878 | HudsonAlpha+STAT5A_(SC-74442) |
| GM12878 | HudsonAlpha+TAF1 |
| GM12878 | HudsonAlpha+TCF12 |
| GM12878 | HudsonAlpha+TCF3_(SC-349) |
| GM12878 | HudsonAlpha+USF-1 |
| GM12878 | HudsonAlpha+YY1_(SC-281) |
| GM12878 | HudsonAlpha+ZBTB33 |
| GM12878 | HudsonAlpha+ZEB1_(SC-25388) |
| GM12891 | HudsonAlpha+PAX5-C20 |
| GM12891 | HudsonAlpha+Pol2-4H8 |
| GM12891 | HudsonAlpha+Pol2 |
| GM12891 | HudsonAlpha+POU2F2 |
| GM12891 | HudsonAlpha+PU.1 |
| GM12891 | HudsonAlpha+TAF1 |
| GM12891 | HudsonAlpha+YY1_(SC-281) |
| GM12892 | HudsonAlpha+PAX5-C20 |
| GM12892 | HudsonAlpha+Pol2-4H8 |
| GM12892 | HudsonAlpha+Pol2 |
| GM12892 | HudsonAlpha+TAF1 |
| GM12892 | HudsonAlpha+YY1 |
| H1-hESC | HudsonAlpha+ATF2_(SC-81188) |
| H1-hESC | HudsonAlpha+ATF3 |
| H1-hESC | HudsonAlpha+BCL11A |
| H1-hESC | HudsonAlpha+CTCF_(SC-5916) |
| H1-hESC | HudsonAlpha+Egr-1 |
| H1-hESC | HudsonAlpha+FOSL1_(SC-183) |
| H1-hESC | HudsonAlpha+GABP |
| H1-hESC | HudsonAlpha+HDAC2_(SC-6296) |
| H1-hESC | HudsonAlpha+JunD |
| H1-hESC | HudsonAlpha+NANOG_(SC-33759) |
| H1-hESC | HudsonAlpha+NRSF |
| H1-hESC | HudsonAlpha+p300 |
| H1-hESC | HudsonAlpha+Pol2-4H8 |
| H1-hESC | HudsonAlpha+Pol2 |
| H1-hESC | HudsonAlpha+POU5F1_(SC-9081) |
| H1-hESC | HudsonAlpha+Rad21 |
| H1-hESC | HudsonAlpha+RXRA |
| H1-hESC | HudsonAlpha+Sin3Ak-20 |
| H1-hESC | HudsonAlpha+SIX5 |
| H1-hESC | HudsonAlpha+SP1 |
| H1-hESC | HudsonAlpha+SP2_(SC-643) |
| H1-hESC | HudsonAlpha+SP4_(V-20) |
| H1-hESC | HudsonAlpha+SRF |
| H1-hESC | HudsonAlpha+TAF1 |
| H1-hESC | HudsonAlpha+TAF7_(SC-101167) |
| H1-hESC | HudsonAlpha+TCF12 |
| H1-hESC | HudsonAlpha+TEAD4_(SC-101184) |
| H1-hESC | HudsonAlpha+USF-1 |
| H1-hESC | HudsonAlpha+YY1_(SC-281) |
| HCT-116 | HudsonAlpha+Pol2-4H8 |
| HCT-116 | HudsonAlpha+YY1_(SC-281) |
| HCT-116 | HudsonAlpha+ZBTB33 |
| HeLa-S3 | HudsonAlpha+GABP |
| HeLa-S3 | HudsonAlpha+NRSF |
| HeLa-S3 | HudsonAlpha+Pol2 |
| HeLa-S3 | HudsonAlpha+TAF1 |
| HepG2 | HudsonAlpha+ATF3 |
| HepG2 | HudsonAlpha+BHLHE40 |
| HepG2 | HudsonAlpha+CEBPB_(SC-150) |
| HepG2 | HudsonAlpha+CEBPD_(SC-636) |
| HepG2 | HudsonAlpha+CTCF_(SC-5916) |
| HepG2 | HudsonAlpha+ELF1_(SC-631) |
| HepG2 | HudsonAlpha+FOSL2 |
| HepG2 | HudsonAlpha+FOXA1_(SC-101058) |
| HepG2 | HudsonAlpha+FOXA1_(SC-6553) |
| HepG2 | HudsonAlpha+FOXA2_(SC-6554) |
| HepG2 | HudsonAlpha+GABP |
| HepG2 | HudsonAlpha+HDAC2_(SC-6296) |
| HepG2 | HudsonAlpha+HNF4A_(SC-8987) |
| HepG2 | HudsonAlpha+HNF4G_(SC-6558) |
| HepG2 | HudsonAlpha+JunD |
| HepG2 | HudsonAlpha+MBD4_(SC-271530) |
| HepG2 | HudsonAlpha+MYBL2_(SC-81192) |
| HepG2 | HudsonAlpha+NFIC_(SC-81335) |
| HepG2 | HudsonAlpha+NRSF |
| HepG2 | HudsonAlpha+V04+NRSF |
| HepG2 | HudsonAlpha+p300 |
| HepG2 | HudsonAlpha+Pol2-4H8 |
| HepG2 | HudsonAlpha+Pol2 |
| HepG2 | HudsonAlpha+Rad21 |
| HepG2 | HudsonAlpha+RXRA |
| HepG2 | HudsonAlpha+Sin3Ak-20 |
| HepG2 | HudsonAlpha+SP1 |
| HepG2 | HudsonAlpha+SP2_(SC-643) |
| HepG2 | HudsonAlpha+SRF |
| HepG2 | HudsonAlpha+TAF1 |
| HepG2 | HudsonAlpha+TCF12 |
| HepG2 | HudsonAlpha+TEAD4_(SC-101184) |
| HepG2 | HudsonAlpha+USF-1 |
| HepG2 | HudsonAlpha+YY1_(SC-281) |
| HepG2 | HudsonAlpha+ZBTB33 |
| HepG2 | HudsonAlpha+ZBTB7A_(SC-34508) |
| HUVEC | HudsonAlpha+Pol2-4H8 |
| HUVEC | HudsonAlpha+Pol2 |
| K562 | HudsonAlpha+ATF3 |
| K562 | HudsonAlpha+BCL3 |
| K562 | HudsonAlpha+BCLAF1_(SC-101388) |
| K562 | HudsonAlpha+CBX3_(SC-101004) |
| K562 | HudsonAlpha+CEBPB_(SC-150) |
| K562 | HudsonAlpha+CTCF_(SC-5916) |
| K562 | HudsonAlpha+CTCFL_(SC-98982) |
| K562 | HudsonAlpha+E2F6 |
| K562 | HudsonAlpha+Egr-1 |
| K562 | HudsonAlpha+ELF1_(SC-631) |
| K562 | HudsonAlpha+ETS1 |
| K562 | HudsonAlpha+FOSL1_(SC-183) |
| K562 | HudsonAlpha+GABP |
| K562 | HudsonAlpha+GATA2_(SC-267) |
| K562 | HudsonAlpha+HDAC2_(SC-6296) |
| K562 | HudsonAlpha+Max |
| K562 | HudsonAlpha+MEF2A |
| K562 | HudsonAlpha+NR2F2_(SC-271940) |
| K562 | HudsonAlpha+NRSF |
| K562 | HudsonAlpha+PML_(SC-71910) |
| K562 | HudsonAlpha+Pol2-4H8 |
| K562 | HudsonAlpha+Pol2 |
| K562 | HudsonAlpha+PU.1 |
| K562 | HudsonAlpha+Rad21 |
| K562 | HudsonAlpha+Sin3Ak-20 |
| K562 | HudsonAlpha+SIX5 |
| K562 | HudsonAlpha+SP1 |
| K562 | HudsonAlpha+SP2_(SC-643) |
| K562 | HudsonAlpha+SRF |
| K562 | HudsonAlpha+STAT5A_(SC-74442) |
| K562 | HudsonAlpha+TAF1 |
| K562 | HudsonAlpha+TAF7_(SC-101167) |
| K562 | HudsonAlpha+TEAD4_(SC-101184) |
| K562 | HudsonAlpha+THAP1_(SC-98174) |
| K562 | HudsonAlpha+TRIM28_(SC-81411) |
| K562 | HudsonAlpha+USF-1 |
| K562 | HudsonAlpha+YY1_(SC-281) |
| K562 | HudsonAlpha+YY1 |
| K562 | HudsonAlpha+ZBTB33 |
| K562 | HudsonAlpha+ZBTB7A_(SC-34508) |
| PANC-1 | HudsonAlpha+NRSF |
| PANC-1 | HudsonAlpha+Pol2-4H8 |
| PANC-1 | HudsonAlpha+Sin3Ak-20 |
| PFSK-1 | HudsonAlpha+FOXP2 |
| PFSK-1 | HudsonAlpha+NRSF |
| PFSK-1 | HudsonAlpha+Sin3Ak-20 |
| PFSK-1 | HudsonAlpha+TAF1 |
| SK-N-MC | HudsonAlpha+FOXP2 |
| SK-N-MC | HudsonAlpha+Pol2-4H8 |
| SK-N-SH | HudsonAlpha+NRSF |
| SK-N-SH | HudsonAlpha+V04+NRSF |
| SK-N-SH | HudsonAlpha+Pol2-4H8 |
| SK-N-SH | HudsonAlpha+Sin3Ak-20 |
| SK-N-SH | HudsonAlpha+TAF1 |
| SK-N-SH_RA | HudsonAlpha+CTCF |
| SK-N-SH_RA | HudsonAlpha+p300 |
| SK-N-SH_RA | HudsonAlpha+Rad21 |
| SK-N-SH_RA | HudsonAlpha+USF1_(SC-8983) |
| SK-N-SH_RA | HudsonAlpha+YY1_(SC-281) |
| T-47D | DMSO_0.02pct+HudsonAlpha+CTCF_(SC-5916) |
| T-47D | BPA_100nM+HudsonAlpha+ERalpha_a |
| T-47D | Genistein_100nM+HudsonAlpha+ERalpha_a |
| T-47D | Estradiol_10nM+HudsonAlpha+ERalpha_a |
| T-47D | DMSO_0.02pct+HudsonAlpha+FOXA1_(SC-6553) |
| T-47D | DMSO_0.02pct+HudsonAlpha+GATA3_(SC-268) |
| T-47D | DMSO_0.02pct+HudsonAlpha+p300 |
| U87 | HudsonAlpha+NRSF |
| U87 | HudsonAlpha+Pol2-4H8 |
| A549 | Stanford+BHLHE40 |
| A549 | Stanford+CEBPB |
| A549 | Stanford+Max |
| A549 | Stanford+Pol2(phosphoS2) |
| A549 | Stanford+Rad21 |
| GM08714 | USC+ZNF274 |
| GM10847 | TNFa+Stanford+NFKB |
| GM10847 | Stanford+Pol2 |
| GM12878 | Stanford+BHLHE40_(NB100-1800) |
| GM12878 | Stanford+BRCA1_(A300-000A) |
| GM12878 | Yale+c-Fos |
| GM12878 | Stanford+CHD1_(A301-218A) |
| GM12878 | Stanford+CHD2_(AB68301) |
| GM12878 | Stanford+COREST_(sc-30189) |
| GM12878 | Stanford+CTCF_(SC-15914) |
| GM12878 | Stanford+E2F4 |
| GM12878 | Stanford+EBF1_(SC-137065) |
| GM12878 | Stanford+ELK1_(1277-1) |
| GM12878 | USC+IKZF1_(IkN)_(UCLA) |
| GM12878 | Yale+JunD |
| GM12878 | Stanford+Max |
| GM12878 | Stanford+MAZ_(ab85725) |
| GM12878 | Stanford+Mxi1_(AF4185) |
| GM12878 | Stanford+NF-E2_(SC-22827) |
| GM12878 | TNFa+Stanford+NFKB |
| GM12878 | Harvard+NF-YA |
| GM12878 | Harvard+NF-YB |
| GM12878 | Stanford+Nrf1 |
| GM12878 | Stanford+p300 |
| GM12878 | Stanford+p300_(SC-584) |
| GM12878 | Stanford+Pol2 |
| GM12878 | Yale+Pol2 |
| GM12878 | Stanford+Pol2(phosphoS2) |
| GM12878 | Yale+Pol3 |
| GM12878 | Stanford+Rad21 |
| GM12878 | Stanford+RFX5_(200-401-194) |
| GM12878 | Stanford+SIN3A_(NB600-1263) |
| GM12878 | Stanford+SMC3_(ab9263) |
| GM12878 | Stanford+STAT1 |
| GM12878 | Stanford+STAT3 |
| GM12878 | Stanford+TBLR1_(ab24550) |
| GM12878 | Stanford+TBP |
| GM12878 | USC+TR4 |
| GM12878 | Stanford+USF2 |
| GM12878 | Stanford+WHIP |
| GM12878 | USC+YY1 |
| GM12878 | Stanford+Znf143_(16618-1-AP) |
| GM12878 | USC+ZNF274 |
| GM12878 | Harvard+ZZZ3 |
| GM12891 | TNFa+Stanford+NFKB |
| GM12891 | Stanford+Pol2 |
| GM12892 | TNFa+Stanford+NFKB |
| GM12892 | Stanford+Pol2 |
| GM15510 | TNFa+Stanford+NFKB |
| GM15510 | Stanford+Pol2 |
| GM18505 | TNFa+Stanford+NFKB |
| GM18505 | Stanford+Pol2 |
| GM18526 | TNFa+Stanford+NFKB |
| GM18526 | Stanford+Pol2 |
| GM18951 | TNFa+Stanford+NFKB |
| GM18951 | Stanford+Pol2 |
| GM19099 | TNFa+Stanford+NFKB |
| GM19099 | Stanford+Pol2 |
| GM19193 | TNFa+Stanford+NFKB |
| GM19193 | Stanford+Pol2 |
| H1-hESC | Stanford+Bach1_(sc-14700) |
| H1-hESC | Stanford+BRCA1_(A300-000A) |
| H1-hESC | Stanford+CEBPB |
| H1-hESC | Stanford+CHD1_(A301-218A) |
| H1-hESC | Stanford+CHD2_(AB68301) |
| H1-hESC | Stanford+c-Jun |
| H1-hESC | Stanford+c-Myc |
| H1-hESC | USC+CtBP2 |
| H1-hESC | Stanford+GTF2F1_(AB28179) |
| H1-hESC | Stanford+JunD |
| H1-hESC | Stanford+MafK_(ab50322) |
| H1-hESC | USC+Max |
| H1-hESC | Stanford+Mxi1_(AF4185) |
| H1-hESC | Stanford+Nrf1 |
| H1-hESC | Stanford+Rad21 |
| H1-hESC | Stanford+RFX5_(200-401-194) |
| H1-hESC | Stanford+SIN3A_(NB600-1263) |
| H1-hESC | USC+SUZ12 |
| H1-hESC | Stanford+TBP |
| H1-hESC | Stanford+USF2 |
| H1-hESC | Stanford+Znf143_(16618-1-AP) |
| HCT-116 | USC+Pol2 |
| HCT-116 | USC+TCF7L2 |
| HEK293 | USC+ELK4 |
| HEK293 | USC+KAP1 |
| HEK293 | Yale+Pol2 |
| HEK293 | USC+TCF7L2 |
| HEK293-T-REx | USC+ZNF263 |
| HeLa-S3 | USC+AP-2alpha |
| HeLa-S3 | USC+AP-2gamma |
| HeLa-S3 | Stanford+BAF155 |
| HeLa-S3 | Stanford+BAF170 |
| HeLa-S3 | Harvard+BDP1 |
| HeLa-S3 | Stanford+BRCA1_(A300-000A) |
| HeLa-S3 | Harvard+BRF1 |
| HeLa-S3 | Harvard+BRF2 |
| HeLa-S3 | Yale+Brg1 |
| HeLa-S3 | Stanford+CEBPB |
| HeLa-S3 | Yale+c-Fos |
| HeLa-S3 | Stanford+CHD2_(AB68301) |
| HeLa-S3 | Stanford+c-Jun |
| HeLa-S3 | Yale+c-Myc |
| HeLa-S3 | Stanford+COREST_(sc-30189) |
| HeLa-S3 | USC+E2F1 |
| HeLa-S3 | USC+E2F4 |
| HeLa-S3 | USC+E2F6 |
| HeLa-S3 | Stanford+ELK1_(1277-1) |
| HeLa-S3 | USC+ELK4 |
| HeLa-S3 | Stanford+GTF2F1_(AB28179) |
| HeLa-S3 | USC+HA-E2F1 |
| HeLa-S3 | Stanford+Ini1 |
| HeLa-S3 | Stanford+IRF3 |
| HeLa-S3 | Stanford+JunD |
| HeLa-S3 | Stanford+MafK_(ab50322) |
| HeLa-S3 | Stanford+Max |
| HeLa-S3 | Stanford+MAZ_(ab85725) |
| HeLa-S3 | Stanford+Mxi1_(AF4185) |
| HeLa-S3 | Harvard+NF-YA |
| HeLa-S3 | Harvard+NF-YB |
| HeLa-S3 | Stanford+Nrf1 |
| HeLa-S3 | Stanford+p300_(SC-584) |
| HeLa-S3 | Yale+Pol2 |
| HeLa-S3 | Stanford+Pol2(phosphoS2) |
| HeLa-S3 | Stanford+PRDM1_(9115) |
| HeLa-S3 | Stanford+Rad21 |
| HeLa-S3 | Stanford+RFX5_(200-401-194) |
| HeLa-S3 | Harvard+RPC155 |
| HeLa-S3 | Stanford+SMC3_(ab9263) |
| HeLa-S3 | Stanford+SPT20 |
| HeLa-S3 | IFNg30+Yale+STAT1 |
| HeLa-S3 | Stanford+STAT3 |
| HeLa-S3 | Stanford+TBP |
| HeLa-S3 | USC+TCF7L2 |
| HeLa-S3 | USC+TCF7L2_C9B9_(2565) |
| HeLa-S3 | Harvard+TFIIIC-110 |
| HeLa-S3 | USC+TR4 |
| HeLa-S3 | Stanford+USF2 |
| HeLa-S3 | Stanford+ZKSCAN1_(HPA006672) |
| HeLa-S3 | Stanford+Znf143_(16618-1-AP) |
| HeLa-S3 | USC+ZNF274 |
| HeLa-S3 | Harvard+ZZZ3 |
| HepG2 | Stanford+ARID3A_(NB100-279) |
| HepG2 | Stanford+BHLHE40_(NB100-1800) |
| HepG2 | Stanford+BRCA1_(A300-000A) |
| HepG2 | forskolin+Stanford+CEBPB |
| HepG2 | Stanford+CEBPB |
| HepG2 | Stanford+CHD2_(AB68301) |
| HepG2 | Stanford+c-Jun |
| HepG2 | Stanford+COREST_(sc-30189) |
| HepG2 | forskolin+Stanford+ERRA |
| HepG2 | forskolin+Stanford+GRp20 |
| HepG2 | forskolin+Stanford+HNF4A |
| HepG2 | forskolin+Stanford+HSF1 |
| HepG2 | Stanford+IRF3 |
| HepG2 | Stanford+JunD |
| HepG2 | Stanford+MafF_(M8194) |
| HepG2 | Stanford+MafK_(ab50322) |
| HepG2 | Stanford+MafK_(SC-477) |
| HepG2 | Stanford+Max |
| HepG2 | Stanford+MAZ_(ab85725) |
| HepG2 | Stanford+Mxi1_(AF4185) |
| HepG2 | Stanford+Nrf1 |
| HepG2 | Stanford+p300_(SC-584) |
| HepG2 | forskolin+Stanford+PGC1A |
| HepG2 | forskolin+Stanford+Pol2 |
| HepG2 | Stanford+Pol2 |
| HepG2 | Stanford+Pol2(phosphoS2) |
| HepG2 | Stanford+Rad21 |
| HepG2 | Stanford+RFX5_(200-401-194) |
| HepG2 | Stanford+SMC3_(ab9263) |
| HepG2 | insulin+Stanford+SREBP1 |
| HepG2 | Stanford+TBP |
| HepG2 | USC+TCF7L2 |
| HepG2 | USC+TR4 |
| HepG2 | Stanford+USF2 |
| HepG2 | USC+ZNF274 |
| HUVEC | USC+c-Fos |
| HUVEC | Stanford+c-Jun |
| HUVEC | USC+GATA-2 |
| HUVEC | Stanford+Max |
| HUVEC | Stanford+Pol2 |
| IMR90 | Stanford+CEBPB |
| IMR90 | Stanford+CTCF_(SC-15914) |
| IMR90 | Stanford+MafK_(ab50322) |
| IMR90 | Stanford+Pol2 |
| IMR90 | Stanford+Rad21 |
| K562 | Stanford+ARID3A_(sc-8821) |
| K562 | Harvard+ATF1_(06-325) |
| K562 | Harvard+ATF3 |
| K562 | Stanford+Bach1_(sc-14700) |
| K562 | Harvard+BDP1 |
| K562 | Stanford+BHLHE40_(NB100-1800) |
| K562 | Harvard+BRF1 |
| K562 | Harvard+BRF2 |
| K562 | Stanford+Brg1 |
| K562 | Harvard+CCNT2 |
| K562 | Stanford+CEBPB |
| K562 | Yale+c-Fos |
| K562 | Stanford+CHD2_(AB68301) |
| K562 | IFNa30+Stanford+c-Jun |
| K562 | IFNa6h+Yale+c-Jun |
| K562 | IFNg30+Yale+c-Jun |
| K562 | IFNg6h+Yale+c-Jun |
| K562 | Yale+c-Jun |
| K562 | IFNa30+Yale+c-Myc |
| K562 | IFNa6h+Yale+c-Myc |
| K562 | IFNg30+Stanford+c-Myc |
| K562 | IFNg6h+Yale+c-Myc |
| K562 | Stanford+c-Myc |
| K562 | Yale+c-Myc |
| K562 | Stanford+COREST_(ab24166) |
| K562 | Stanford+COREST_(sc-30189) |
| K562 | Stanford+CTCF_(SC-15914) |
| K562 | USC+E2F4 |
| K562 | USC+E2F6 |
| K562 | Stanford+ELK1_(1277-1) |
| K562 | USC+GATA-1 |
| K562 | USC+GATA-2 |
| K562 | Harvard+GTF2B |
| K562 | Stanford+GTF2F1_(AB28179) |
| K562 | Harvard+HMGN3 |
| K562 | Stanford+Ini1 |
| K562 | IFNa30+Stanford+IRF1 |
| K562 | IFNa6h+Stanford+IRF1 |
| K562 | IFNg30+Stanford+IRF1 |
| K562 | IFNg6h+Stanford+IRF1 |
| K562 | Stanford+JunD |
| K562 | USC+KAP1 |
| K562 | Stanford+MafF_(M8194) |
| K562 | Stanford+MafK_(ab50322) |
| K562 | Stanford+Max |
| K562 | Stanford+MAZ_(ab85725) |
| K562 | Stanford+Mxi1_(AF4185) |
| K562 | Harvard+NELFe |
| K562 | Yale+NF-E2 |
| K562 | Stanford+NF-YA |
| K562 | Stanford+NF-YB |
| K562 | Stanford+Nrf1 |
| K562 | Stanford+p300 |
| K562 | IFNa30+Yale+Pol2 |
| K562 | IFNa6h+Yale+Pol2 |
| K562 | IFNg30+Stanford+Pol2 |
| K562 | IFNg6h+Yale+Pol2 |
| K562 | Stanford+Pol2 |
| K562 | Yale+Pol2 |
| K562 | Stanford+Iggrab+Pol2(phosphoS2) |
| K562 | Stanford+Pol2(phosphoS2) |
| K562 | Stanford+Pol3 |
| K562 | Yale+Rad21 |
| K562 | Stanford+RFX5_(200-401-194) |
| K562 | Harvard+RPC155 |
| K562 | MNaseD+USC+SETDB1 |
| K562 | USC+SETDB1 |
| K562 | Harvard+SIRT6 |
| K562 | Stanford+SMC3_(ab9263) |
| K562 | IFNa30+Yale+STAT1 |
| K562 | IFNa6h+Yale+STAT1 |
| K562 | IFNg30+Stanford+STAT1 |
| K562 | IFNg6h+Stanford+STAT1 |
| K562 | IFNa30+Yale+STAT2 |
| K562 | IFNa6h+Yale+STAT2 |
| K562 | Stanford+TAL1_(SC-12984) |
| K562 | Stanford+TBLR1_(ab24550) |
| K562 | Stanford+TBLR1_(NB600-270) |
| K562 | Stanford+TBP |
| K562 | Harvard+TFIIIC-110 |
| K562 | USC+TR4 |
| K562 | Stanford+UBF_(sc-13125) |
| K562 | Stanford+UBTF_(SAB1404509) |
| K562 | Stanford+USF2 |
| K562 | USC+YY1 |
| K562 | Stanford+Znf143_(16618-1-AP) |
| K562 | USC+ZNF263 |
| K562 | USC+ZNF274 |
| K562 | USC+ZNF274_(M01) |
| MCF10A-Er-Src | EtOH_0.01pct+Harvard+c-Fos |
| MCF10A-Er-Src | 4OHTAM_1uM_12hr+Harvard+c-Fos |
| MCF10A-Er-Src | 4OHTAM_1uM_4hr+Harvard+c-Fos |
| MCF10A-Er-Src | 4OHTAM_1uM_36hr+Harvard+c-Fos |
| MCF10A-Er-Src | EtOH_0.01pct+Harvard+c-Myc |
| MCF10A-Er-Src | 4OHTAM_1uM_4hr+Harvard+c-Myc |
| MCF10A-Er-Src | 4OHTAM_1uM_36hr+Harvard+E2F4 |
| MCF10A-Er-Src | EtOH_0.01pct+Harvard+Pol2 |
| MCF10A-Er-Src | 4OHTAM_1uM_36hr+Harvard+Pol2 |
| MCF10A-Er-Src | EtOH_0.01pct+Harvard+STAT3 |
| MCF10A-Er-Src | EtOH_0.01pct_4hr+Harvard+STAT3 |
| MCF10A-Er-Src | EtOH_0.01pct_12hr+Harvard+STAT3 |
| MCF10A-Er-Src | 4OHTAM_1uM_12hr+Harvard+STAT3 |
| MCF10A-Er-Src | 4OHTAM_1uM_36hr+Harvard+STAT3 |
| MCF-7 | USC+GATA3_(SC-268) |
| MCF-7 | USC+GATA3_(SC-269) |
| MCF-7 | USC+HA-E2F1 |
| MCF-7 | USC+TCF7L2 |
| MCF-7 | USC+ZNF217 |
| NB4 | Stanford+c-Myc |
| NB4 | Stanford+Max |
| NB4 | Yale+Pol2 |
| NT2-D1 | USC+SUZ12 |
| NT2-D1 | USC+YY1 |
| NT2-D1 | USC+ZNF274 |
| PANC-1 | USC+TCF7L2 |
| PBDE | USC+GATA-1 |
| PBDE | USC+Pol2 |
| PBDEFetal | USC+GATA-1 |
| Raji | USC+Pol2 |
| SH-SY5Y | USC+GATA-2 |
| SH-SY5Y | USC+GATA3_(SC-269) |
| U2OS | USC+KAP1 |
| U2OS | USC+SETDB1 |
| K562 | UChicago+eGFP-FOS |
| K562 | UChicago+eGFP-GATA2 |
| K562 | UChicago+eGFP-HDAC8 |
| K562 | UChicago+eGFP-JunB |
| K562 | UChicago+eGFP-JunD |
| A549 | UT-A+CTCF |
| A549 | UT-A+Pol2 |
| Fibrobl | UT-A+CTCF |
| Gliobla | UT-A+CTCF |
| Gliobla | UT-A+Pol2 |
| GM12878 | UT-A+c-Myc |
| GM12878 | UT-A+CTCF |
| GM12878 | UT-A+Pol2 |
| GM12891 | UT-A+CTCF |
| GM12892 | UT-A+CTCF |
| GM19238 | UT-A+CTCF |
| GM19239 | UT-A+CTCF |
| GM19240 | UT-A+CTCF |
| H1-hESC | UT-A+c-Myc |
| H1-hESC | UT-A+CTCF |
| H1-hESC | UT-A+Pol2 |
| HeLa-S3 | UT-A+c-Myc |
| HeLa-S3 | UT-A+CTCF |
| HeLa-S3 | UT-A+Pol2 |
| HepG2 | UT-A+c-Myc |
| HepG2 | UT-A+CTCF |
| HepG2 | UT-A+Pol2 |
| HUVEC | UT-A+c-Myc |
| HUVEC | UT-A+CTCF |
| HUVEC | UT-A+Pol2 |
| K562 | UT-A+c-Myc |
| K562 | UT-A+CTCF |
| K562 | UT-A+Pol2 |
| MCF-7 | estrogen+UT-A+c-Myc |
| MCF-7 | serum_stimulated_media+UT-A+c-Myc |
| MCF-7 | serum_starved_media+UT-A+c-Myc |
| MCF-7 | vehicle+UT-A+c-Myc |
| MCF-7 | estrogen+UT-A+CTCF |
| MCF-7 | serum_stimulated_media+UT-A+CTCF |
| MCF-7 | serum_starved_media+UT-A+CTCF |
| MCF-7 | UT-A+CTCF |
| MCF-7 | vehicle+UT-A+CTCF |
| MCF-7 | serum_stimulated_media+UT-A+Pol2 |
| MCF-7 | serum_starved_media+UT-A+Pol2 |
| MCF-7 | UT-A+Pol2 |
| NHEK | UT-A+CTCF |
| ProgFib | UT-A+CTCF |
| ProgFib | UT-A+Pol2 |
| A549 | UW+CTCF |
| AG04449 | UW+CTCF |
| AG04450 | UW+CTCF |
| AG09309 | UW+CTCF |
| AG09319 | UW+CTCF |
| AG10803 | UW+CTCF |
| AoAF | UW+CTCF |
| BE2_C | UW+CTCF |
| BJ | UW+CTCF |
| Caco-2 | UW+CTCF |
| GM06990 | UW+CTCF |
| GM12801 | UW+CTCF |
| GM12864 | UW+CTCF |
| GM12865 | UW+CTCF |
| GM12872 | UW+CTCF |
| GM12873 | UW+CTCF |
| GM12874 | UW+CTCF |
| GM12875 | UW+CTCF |
| GM12878 | UW+CTCF |
| HAc | UW+CTCF |
| HA-sp | UW+CTCF |
| HBMEC | UW+CTCF |
| HCFaa | UW+CTCF |
| HCM | UW+CTCF |
| HCPEpiC | UW+CTCF |
| HCT-116 | UW+CTCF |
| HEEpiC | UW+CTCF |
| HEK293 | UW+CTCF |
| HeLa-S3 | UW+CTCF |
| HepG2 | UW+CTCF |
| HFF | UW+CTCF |
| HFF-Myc | UW+CTCF |
| HL-60 | UW+CTCF |
| HMEC | UW+CTCF |
| HMF | UW+CTCF |
| HPAF | UW+CTCF |
| HPF | UW+CTCF |
| HRE | UW+CTCF |
| HRPEpiC | UW+CTCF |
| HUVEC | UW+CTCF |
| HVMF | UW+CTCF |
| K562 | UW+CTCF |
| MCF-7 | UW+CTCF |
| NB4 | UW+CTCF |
| NHDF-neo | UW+CTCF |
| NHEK | UW+CTCF |
| NHLF | UW+CTCF |
| RPTEC | UW+CTCF |
| SAEC | UW+CTCF |
| SK-N-SH_RA | UW+CTCF |
| WERI-Rb-1 | UW+CTCF |
| WI-38 | UW+CTCF |
